# Integrated Modeling of BCR/TCR Repertoire Diversity Reveals the Mechanistic Basis of Immune Imprinting and Chronic Infection Control

**DOI:** 10.64898/2026.02.26.708225

**Authors:** Zhaobin Xu, Shiyong Wen, Cheng Liu, Weiqi Zheng, Qiangcheng Zeng, Guangyu Yang, Jian Song, Hongmei Zhang, Lu Liu, Chunhua Li, Liyan Wang, Baidong Hou, Dongqing Wei

**Author notes:** Correspondence (ZX).

## Abstract

The adaptive immune system orchestrates a complex interplay between humoral and cellular responses to resolve pathogen invasion and suppress tumorigenesis [1,91]. However, a unified quantitative framework integrating lymphocyte repertoire diversity with system-level kinetics remains elusive. Here, we present a multi-scale mathematical model that explicitly incorporates B-cell receptor (BCR) and T-cell receptor (TCR) clonotypic diversity into a discrete agent-based simulation. This framework mechanistically elucidates pivotal immunological phenomena, including the affinity-dependent selection in germinal centers [25], the kinetic constraints of immune imprinting (“original antigenic sin”) [54], and the bifurcation between acute and chronic infection trajectories [15,59]. We identify that chronic infections are sustained by specific kinetic thresholds of viral replication relative to antibody affinity maturation. Furthermore, our simulations of cancer-immune interactions demonstrate that therapeutic efficacy is non-linearly dependent on neoantigen presentation [78] and T-cell exhaustion thresholds [79]. This work bridges the gap between reductionist data and systems biology, providing a “quantitative immunology” platform to optimize vaccine dosing schedules and design precision immunotherapies.

## Main

The historical expansion of human life expectancy is inextricably linked to our growing mastery of immunology, evidenced by the strong correlation between global population surges and the universal implementation of vaccination programs [1–2]. The 20th century marked a paradigm shift in dissecting the adaptive immune system, particularly through the elucidation of the stochastic mechanisms governing B-cell receptor (BCR) and T-cell receptor (TCR) diversification. This diversity is the bedrock of immunity: BCR variation generates a vast antibody repertoire capable of recognizing infinite antigens, while TCR polymorphism dictates the specificity of helper (CD4^+^) and cytotoxic (CD8^+^) T cell responses [3–5].

In the modern era, the advent of high-throughput technologies—ranging from flow cytometry to single-cell sequencing—has granted us unprecedented access to the static landscapes of immune receptor sequences [6–8]. Furthermore, spatial omics platforms now allow for the detailed mapping of immune cell architectures, resolving the cellular composition and gene expression profiles within critical structures such as germinal centers [9–10]. However, a fundamental gap remains: while we can now catalogue the components of the immune system with high fidelity, understanding the complex, non-linear dynamics that orchestrate these components during viral infection or tumorigenesis remains a formidable challenge. Purely empirical methods struggle to capture the system-level temporal evolution, and data-driven approaches, such as deep learning, often function as “black boxes,” insufficient for deciphering the causal, mechanistic logic of immune regulation. Consequently, there is an urgent imperative for a unified mathematical framework that bridges the gap between reductionist data and systems biology. Such a model must integrate humoral and cellular immunity and be grounded in rigorous biophysical principles to preclude the risks of overfitting. A mechanism-based approach is essential to accurately recapitulate the host response to pathogenic invasion and to provide quantitative insights into emergent immunological phenomena, including the kinetics of secondary infection [11–12], the maintenance of long-term memory [13– 14], the trajectory of CD8^+^ T cell exhaustion [15–16], and the establishment of chronic infections [17–18].

Yet, a generalized, systems-level model that explicitly couples lymphocyte repertoire diversity with post-infection interaction dynamics is largely absent from the current literature. To address this, we developed a comprehensive computational framework that incorporates the stochasticity of BCR and TCR polymorphism, enabling a high-fidelity simulation of adaptive immune responses. By integrating the core mechanisms of cellular and humoral immunity into a discrete agent-based environment, our model is designed to simulate immune dynamics across diverse biological contexts—from acute pathogen challenges to the chronic equilibrium of cancer development. Here, we leverage this platform to elucidate complex immunological behaviors, offering a theoretical basis to optimize vaccine regimens and design precision immunotherapies for chronic infections and malignancies.

## Results

### 2.1 Mechanistic Modeling of Humoral Dynamics and Repertoire Evolution Core Biophysical Framework (Figure 1A)

We first established a stochastic, agent-based framework to simulate the initiation of humoral immunity, as delineated in Figure 1A. Distinct from classical compartmental models, our approach mechanistically couples viral binding kinetics with antigen presentation efficiency. The model postulates that B-cell proliferation is not merely a function of antigen availability but is strictly affinity-dependent: B cells expressing high-affinity BCRs capture and internalize virions more efficiently, thereby presenting a higher density of peptide–MHC-II complexes to CD4^+^ T cells. This enhanced cognate interaction amplifies T-cell help, driving preferential clonal expansion. While dendritic cells are canonically viewed as the primary antigen-presenting cells (APCs), our model integrates recent paradigm-shifting evidence that activated B cells serve as the dominant APCs during the established phase of the humoral response [19–21], creating a positive feedback loop grounded in BCR–antigen binding energy [22–24].

**Figure 1.**
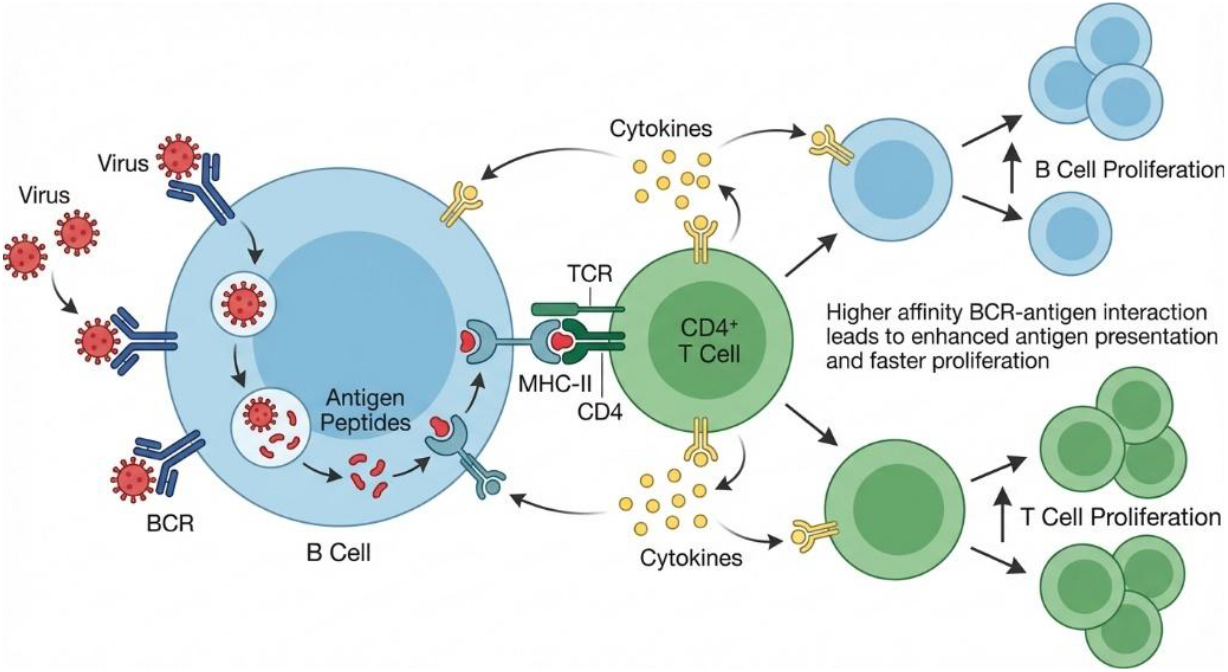
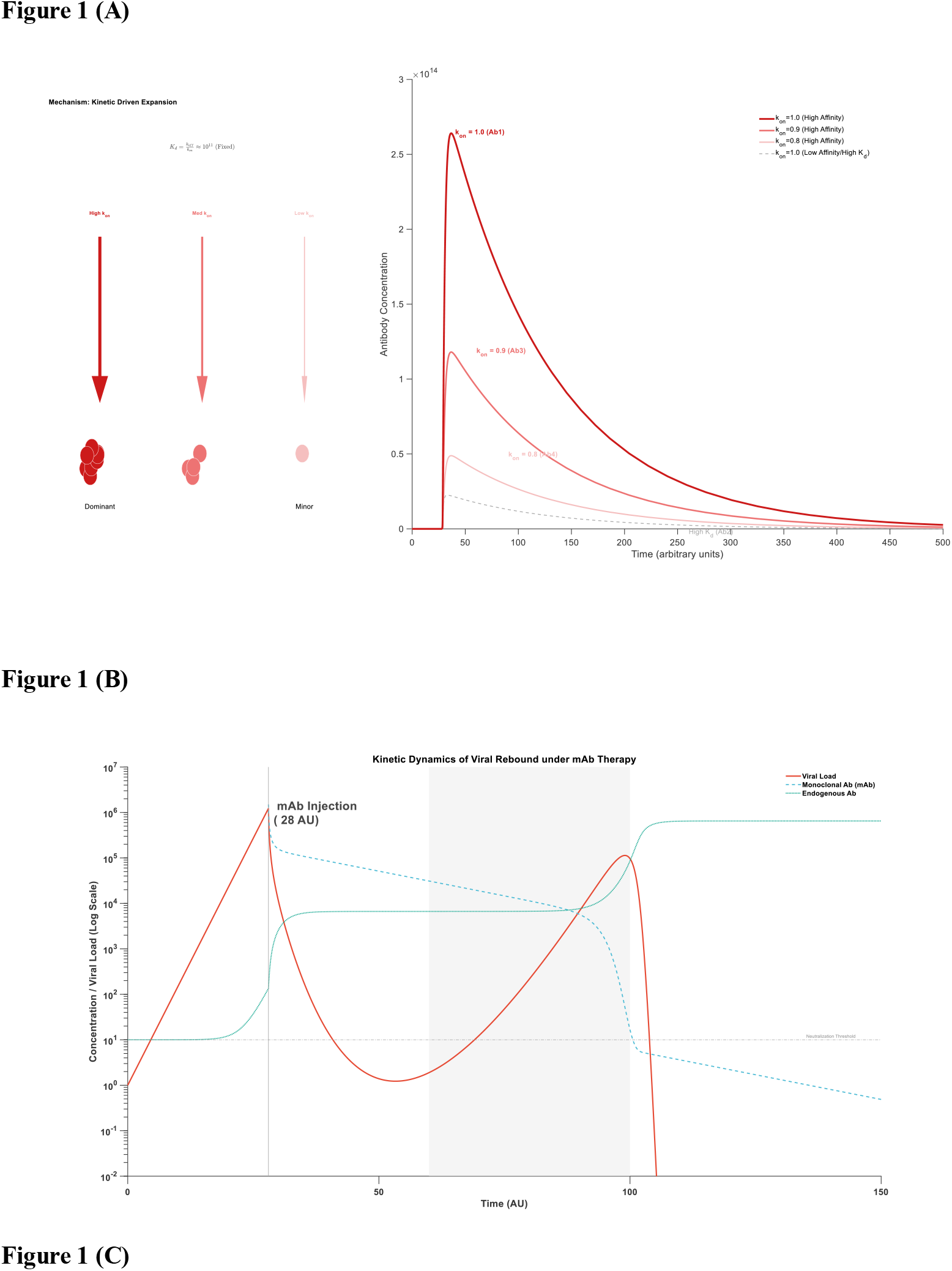
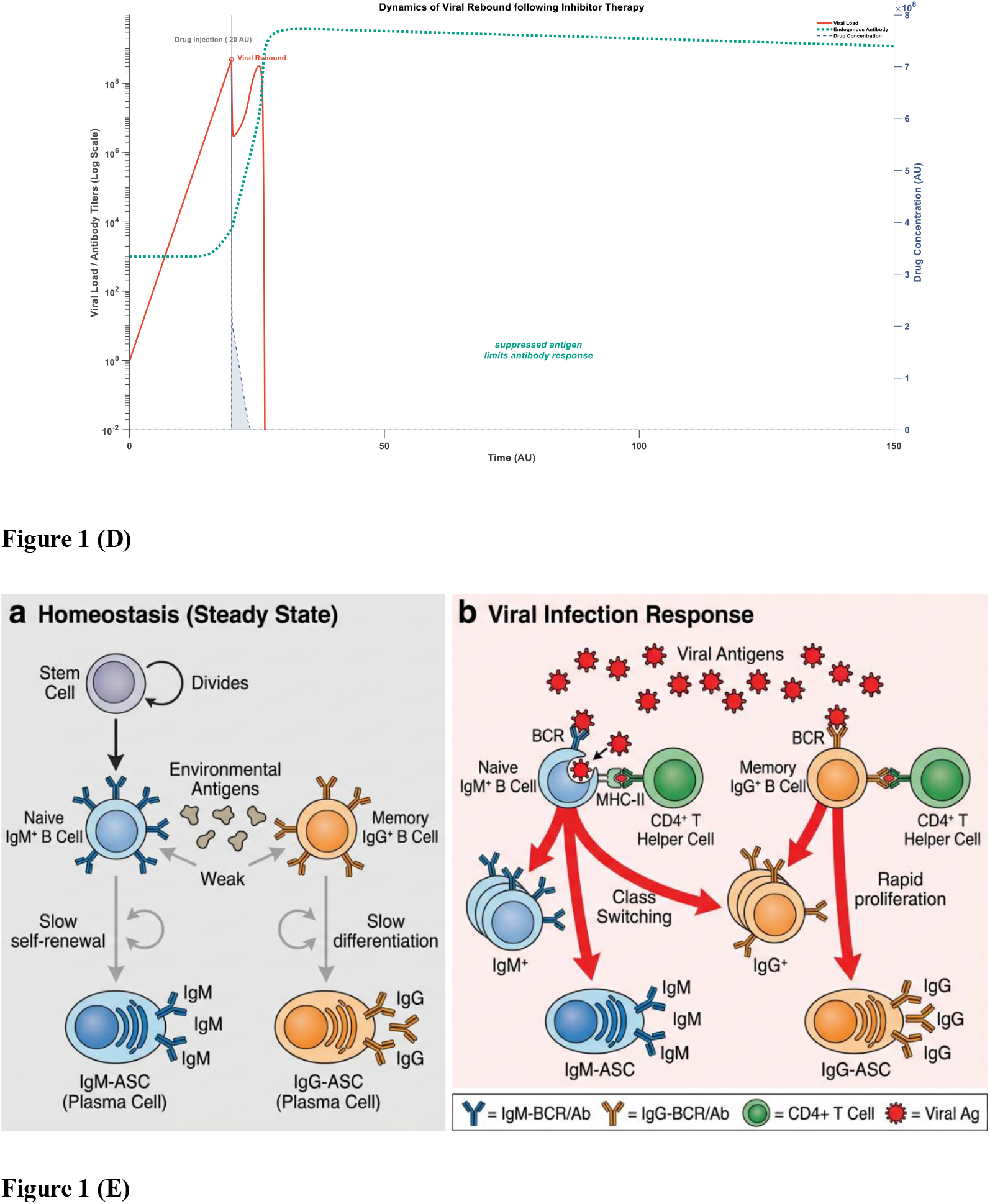
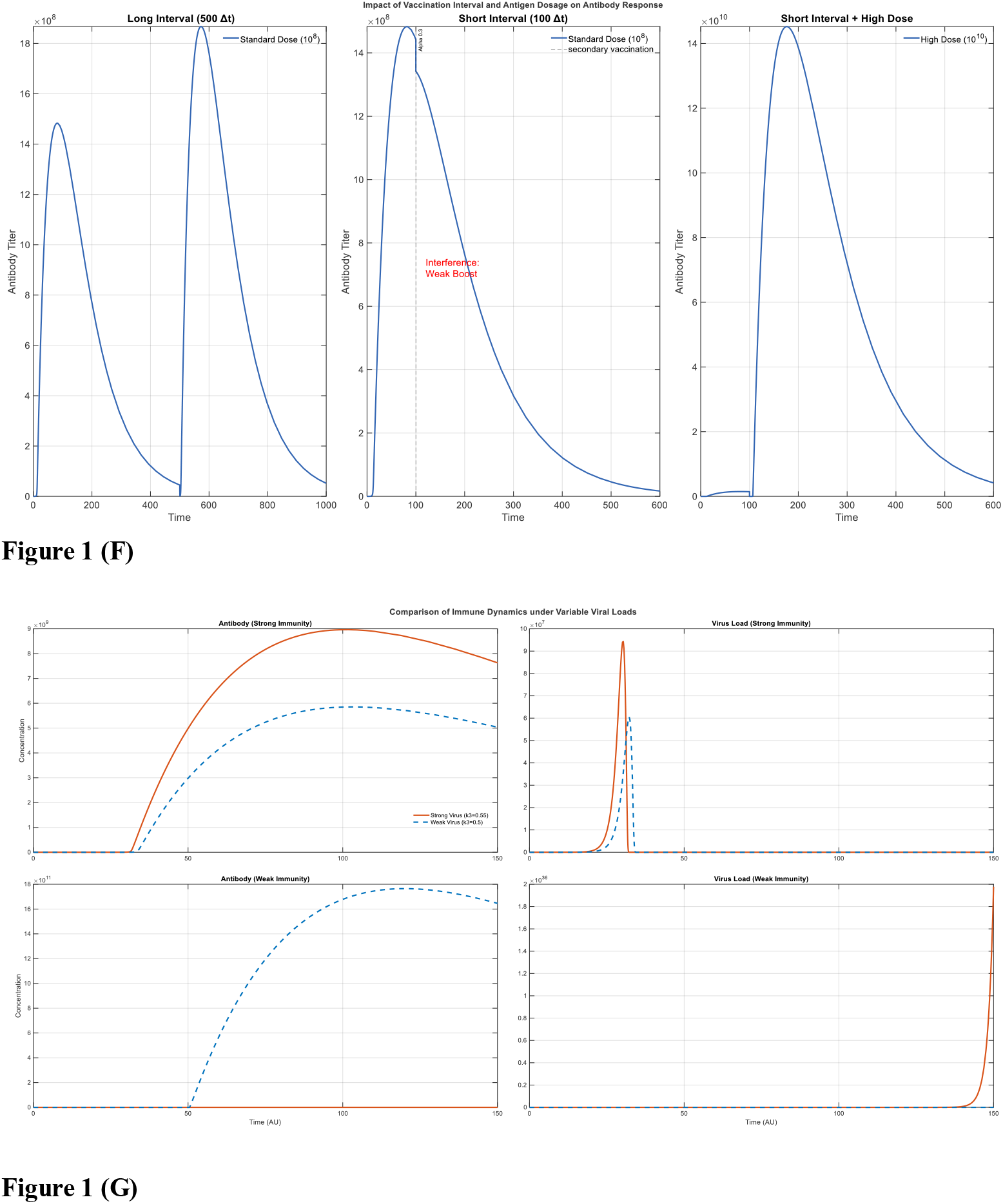
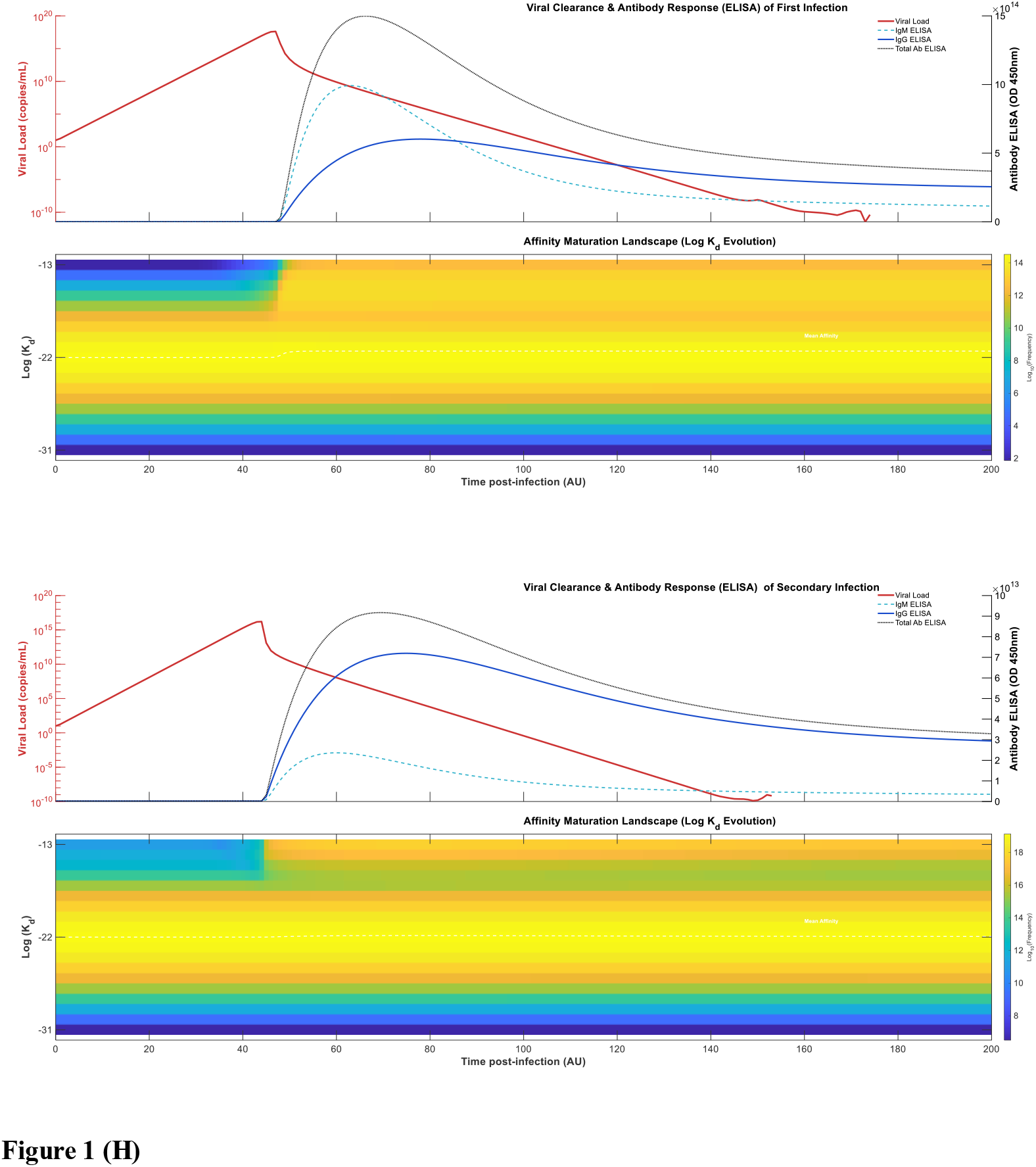
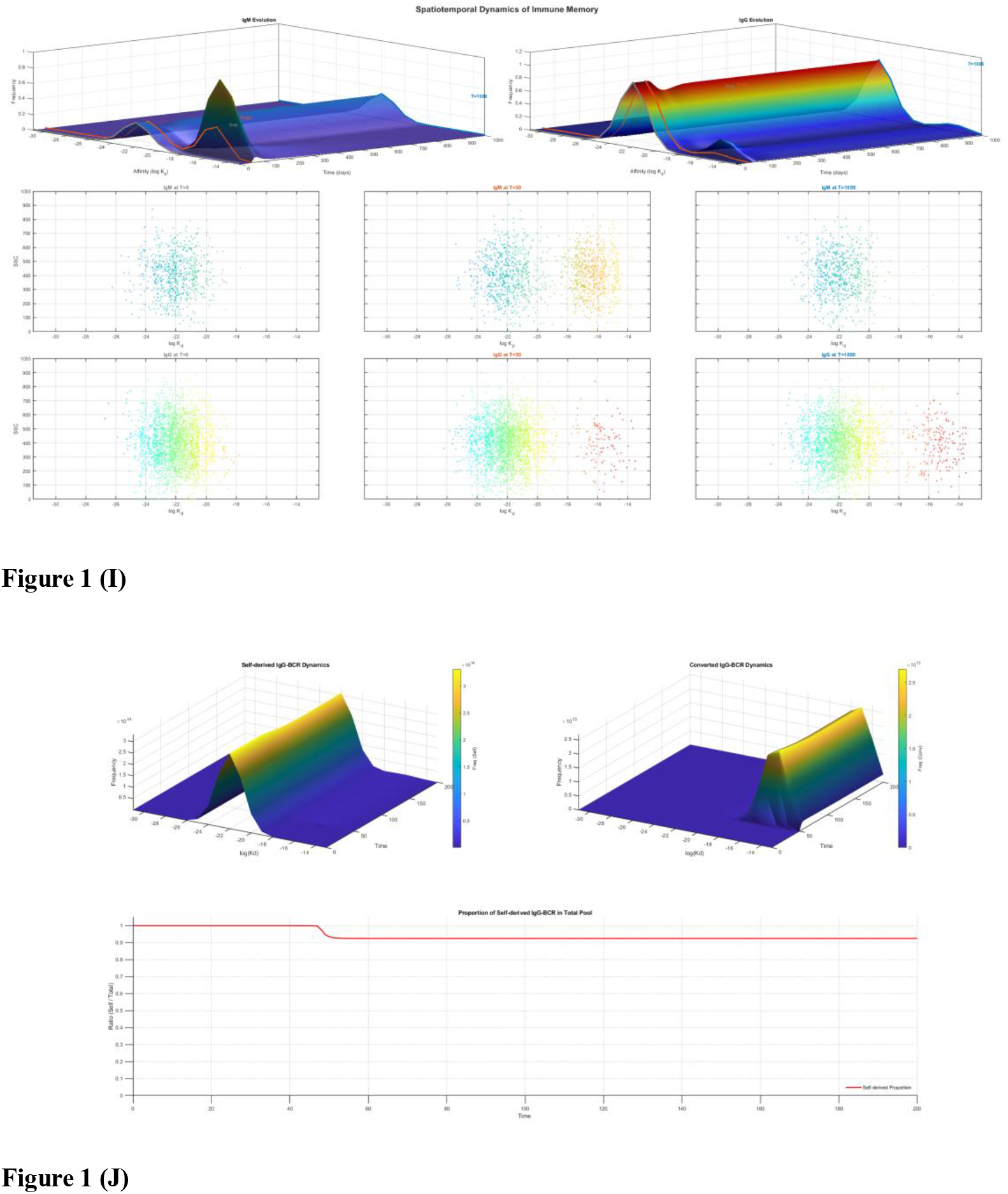
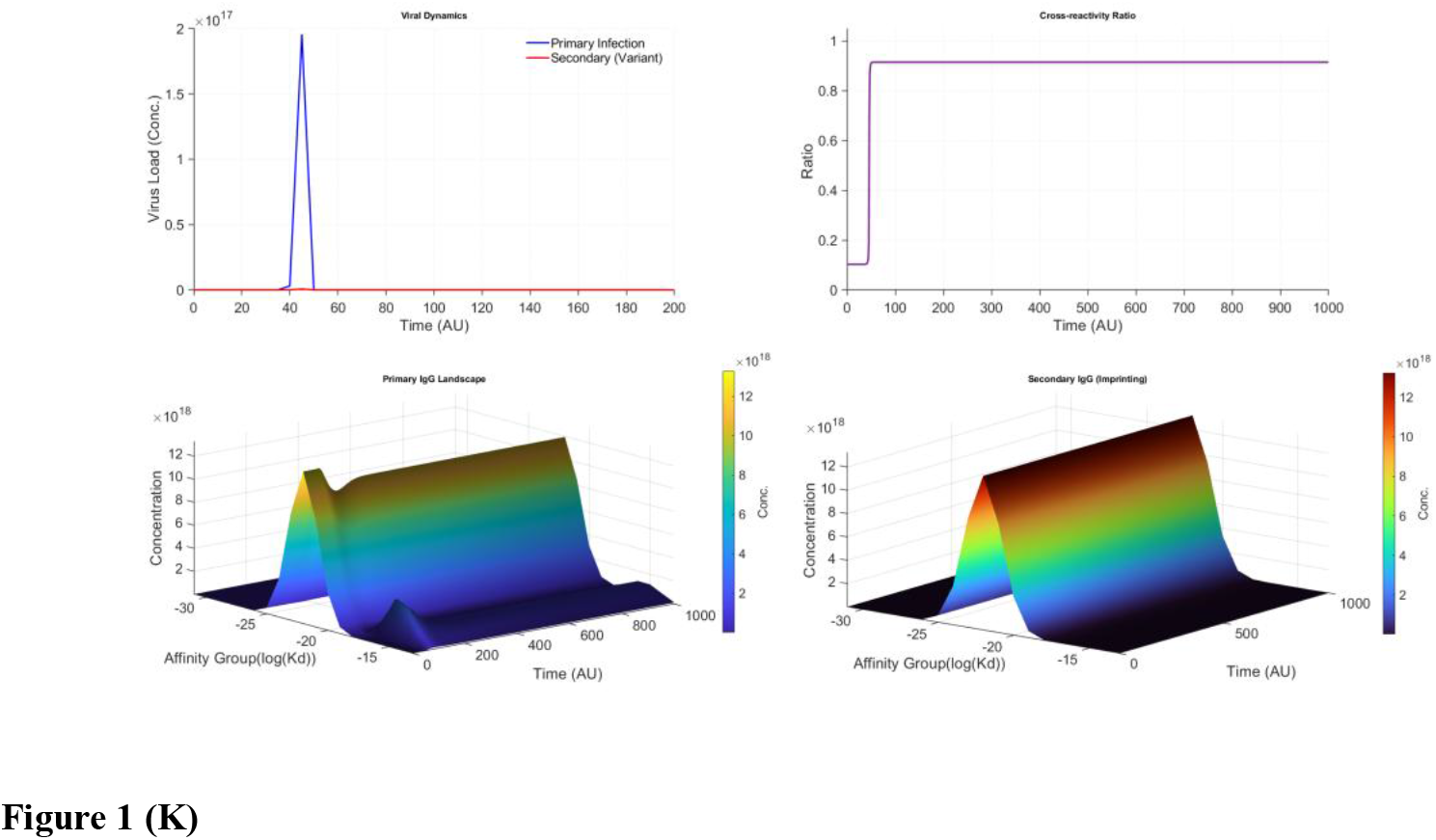
Stochastic modeling of affinity-dependent humoral responses and antibody repertoire evolution. **(A)** Schematic representation of the humoral immune response framework. The model integrates virus-BCR binding kinetics, antigen internalization, and MHC-II presentation to CD4+ T cells, driving affinity-dependent B-cell clonal expansion. **(B)** Dynamics of specific antibody clonal expansion following primary infection. The curves illustrate the selection of high-affinity clones over time. **(C)** Mechanism of viral rebound following monoclonal antibody (mAb) therapy. The dashed cycle highlights the resurgence of viral load occurring when mAb concentrations decay below a critical neutralization threshold before endogenous germinal center re-activation. **(D)** Viral rebound dynamics following small-molecule inhibitor treatment (e.g., Paxlovid analogue). **(E)** Compartmental diagram of the humoral response quantifying the differentiation pathways of IgM+ and IgG+ B cells into respective antibody-secreting plasma cells (ASCs). **(F)** Impact of vaccination scheduling and dosage on antibody kinetics. Comparative analysis of short (100 time-steps) vs. long (500 time-steps) intervals and varying dosages (10^8^ vs 10^10^) reveals that extended intervals favor higher affinity maturation. **(G)** Germinal center (GC) stress response in immunocompromised hosts. Simulation shows that excessive antigen stimulation can lead to early GC exhaustion. **(H)** Differential evolutionary dynamics of IgM and IgG antibody repertoires in primary and secondary infections. Heatmaps illustrating the temporal evolution of the IgG antibody repertoire distribution (binding affinity landscape) throughout the course of infection. During the primary infection, the response is dominated by a rapid, high-amplitude fluidity of IgM-BCRs, driven by the superior binding kinetics of polymeric IgM-BCR structures on the cell surface. In contrast, the IgG response exhibits a significant kinetic lag due to the requirement for isotype switching from IgM and initially lower distinct binding affinities. While the IgM peak is transient and decays rapidly without forming long-term memory, the heatmaps show the progressive consolidation and affinity maturation of the IgG population. This shift establishes a stable reservoir of high-affinity IgG-BCRs, which serves as the basis for immunological memory. Consequently, in a secondary infection, this pre-existing high-affinity IgG pool allows for a dominant, rapid, and robust IgG response that supersedes the IgM response. The color gradient represents the density of IgG clones across the affinity spectrum over time. **(I)** Long-term maintenance of immune memory by high-affinity IgG-BCR subsets. The panels display the temporal evolution of antigen-specific B cell dynamics, projecting the three-dimensional evolutionary landscape onto two-dimensional flow cytometry profiles (bottom rows). The simulation reveals distinct kinetic fates for the two isotypes over an extended timeline: while high-affinity IgM-BCR populations appear transiently and undergo rapid decay to baseline levels, high-affinity IgG-BCR populations are selectively preserved. Crucially, these IgG-BCRs are maintained above a specific homeostatic threshold within the high-affinity region long after the resolution of infection. This persistent high-affinity IgG-BCR reservoir, unlike the transient IgM response, serves as the structural basis for immunological memory. **(J)** Mechanism of discontinuous, stepwise decline in virus A-specific IgG following heterologous virus B infection. Schematic and quantitative analysis of the impact of IgM-to-IgG isotype switching on pre-existing antiviral IgG repertoires. Upon infection with virus B, the de novo IgG response arises from two sources: *de novo* expansion of virus B-specific IgG-BCRs (*p*) and conversion of virus B-primed IgM-BCRs to IgG-BCRs (1-*p*) (denoted by dashed transition arrows). This redirection of the B cell repertoire reduces the proportional contribution of virus A-specific IgG to the total IgG pool, causing virus A-specific IgG levels to decline discontinuously from an initial proportion *q* to a new steady state of *q*×*p*. While total IgG levels transiently increase (*r*) during acute virus B infection (leading to a temporary spike in virus A-specific IgG as *q*×*p*×*r*), this is followed by a resolution phase where virus A-specific IgG stabilizes at the lower *q*×*p* baseline. The lower panel quantifies the magnitude of this decline: each subsequent heterologous infection exacerbates the erosion of virus A-specific IgG, with the total decrement directly proportional to the number of heterologous exposures. This phenomenon reflects the competitive remodeling of the IgG repertoire, where new antigenic challenges prioritize de novo IgM-to-IgG conversion over maintenance of pre-existing pathogen-specific IgG memory. **(K)** Quantitative dynamics of immune imprinting and BCR cross-reactivity. Simulation of the mutant virus kinetics and the concurrent reshaping of the antibody repertoire during infection with a variant strain (variation coefficient = 5). The layout visualizes the amplitude of immune imprinting by quantifying the proportion of B cells exhibiting cross-binding activity (BCR cross-linking ratio) relative to viral homology. The model illustrates that the antibody affinity profile is dictated by the degree of resemblance between the primary and secondary pathogens: high homology maintains the bimodal distribution (shifted toward high-affinity regions) established by the initial infection—characteristic of “original antigenic sin”—whereas divergence shifts the profile toward a normal distribution. This analysis allows for the quantitative determination of the imprint magnitude and the potential for secondary infection by mutant strains.

#### Affinity vs. Clonal Selection (Figure 1B)

Validating the model against fundamental selection principles, we explored the tension between thermodynamic affinity and kinetic association rates (Model 3.1.3). While traditional views emphasize equilibrium dissociation constants (*K*_*d*_), our simulations reveal that the immune system acts as a kinetic filter. As shown in Figure 1B, among antibody clones with identical binding free energies, those with faster on-rates (*k*_*on*_) achieve superior clonal dominance over those with slower off-rates (*k*_*off*_). This suggests that the germinal center reaction selects not just for tight binding, but for rapid antigen capture—a critical feature for neutralizing fast-replicating pathogens.

#### Mechanisms of Viral Rebound (Figure 1C–D)

We next applied this framework to resolve the paradox of viral rebound following antiviral interventions. Contrary to the hypothesis that rebound takes place solely due to viral resistance mutations [28, 30], our model identifies a “kinetic decoupling” mechanism (Model 3.1.4 & 3.1.5). Monoclonal Antibodies (mAbs): We demonstrate that while exogenous mAbs effectively neutralize virions, they do not contribute to antigen presentation or endogenous B-cell priming (Figure 1C). If mAbs are cleared before the endogenous repertoire has matured, the host remains immunologically naive, allowing residual virus to resurge.

Small-Molecule Inhibitors: Similarly, direct viral replication inhibitors suppress the antigen load required to drive affinity maturation (Figure 1D). Premature withdrawal leaves the system in a “sub-threshold” state, leading to rebound.

Collectively, these simulations underscore that successful therapy must balance viral suppression with sufficient antigen exposure to prime endogenous immunity.

#### Differentiation Dynamics and Homeostasis (Figure 1E)

To capture long-term repertoire maintenance, the model integrates distinct B-cell lineages: continuously replenished IgM^+^ pools versus self-renewing IgG^+^ memory compartments (Figure 1E). A novel feature of our framework is the inclusion of “environmental antigens”—a broad spectrum of background stimuli that maintain basal BCR signaling and steady-state immunoglobulin levels in the absence of specific pathogen challenge. This homeostatic mechanism ensures that the rapid decay of short-lived plasma cells is balanced by de novo differentiation, stabilizing the immune landscape.

#### Vaccine Optimization and Germinal Center Collapse (Figure 1F–G)

Leveraging Model 3.1.6, we provided a mechanistic rationale for vaccination scheduling. Our simulations recapitulate the observation that short-interval dosing attenuates booster responses [31– 33]. We attribute this to “epitope masking”: high titers of pre-existing antibodies from the prime dose sequester antigen, preventing BCR engagement on memory B cells. Extending the interval allows antibody decay to “unmask” epitopes, restoring robust activation (Figure 1F).

Furthermore, the model elucidates the pathophysiology of severe infections (e.g., severe COVID-19). We found that an overwhelming antigen load can trigger a “terminal differentiation catastrophe,” where GC B cells effectuate premature mass differentiation into plasma cells, exhausting the proliferative precursor pool (Figure 1G). This kinetic exhaustion leads to GC collapse [34–35], distinguishing functional immunity from the dysregulated responses seen in critical illness. Repertoire Remodeling: The Transient vs. The Persistent (Figure 1H–I)

We further expanded the computational framework to resolve humoral heterogeneity by stratifying IgM-BCR, IgG-BCR, secreted antibodies, and their producing plasma cells into 100 distinct subtypes based on their specific forward association and reverse dissociation rate constants. The kinetic profiles and initial concentrations for each subtype are detailed in Extended Data Figure 1. We longitudinally tracked the evolution of BCR affinities (Model 3.1.7). The model reveals a fundamental bifurcation in lineage fate: the IgM response appears as a transient, high-magnitude wave that rapidly recedes due to the turnover of short-lived plasma cells; conversely, the IgG response establishes a persistent footprint in the high-affinity landscape (Figure 1H). This simulation quantitatively explains why IgM dominates early kinetics yet fails to confer long-term memory [38–39]. The resulting “memory state” is not static but a dynamic equilibrium, where specific IgG clone contract proportionally but maintain a supranormal baseline due to the integration of environmental and specific survival signals (Figure 1I).

#### Theory of Discontinuous Antibody Decay (Figure 1J)

A striking prediction of our model is the mechanism of long-term antibody attrition. We challenge the assumption of simple exponential decay, proposing instead a “discontinuous attrition” model driven by heterologous interference. Each subsequent infection with a distinct pathogen creates a competitive bottleneck in the memory pool (diluting the fraction of pre-existing clones). Our simulations show that specific antibody titers undergo step-wise reductions following heterologous challenges (Figure 1J). This theoretical prediction aligns precisely with serological surveillance data showing varying declines in pre-existing immunity to respiratory viruses following the SARS-CoV-2 pandemic and vaccination campaigns [44–45].

#### Immune Imprinting and the “Homology Threshold” (Figure 1K)

Finally, we offer a quantitative definition of immune imprinting (“Original Antigenic Sin”). Our model demonstrates that primary infection reshapes the BCR repertoire into a bimodal distribution, heavily biased toward high-affinity clones for the ancestral strain. Upon challenge with a variant, the activation of cross-reactive memory cells suppresses the de novo naïve response. We introduce a quantitative metric—the “homology threshold”—governed by the cross-reactivity ratio (Figure 1K). The model predicts that chronicity or susceptibility to variants is determined by whether the viral mutation distance exceeds the breadth of the imprinted repertoire [46–48], providing a calculable basis for vaccine strain selection.

### 2.2 Mechanistic Coupling of Humoral-Cellular Crosstalk

While dendritic cells (DCs) are canonically viewed as the primary orchestrators of cellular immunity, our model necessitates a revised framework that explicitly integrates the regulatory role of humoral components. We delineated two distinct immunological topologies based on pathogen intrinsic immunogenicity (Figure 2A).

**Figure 2.**
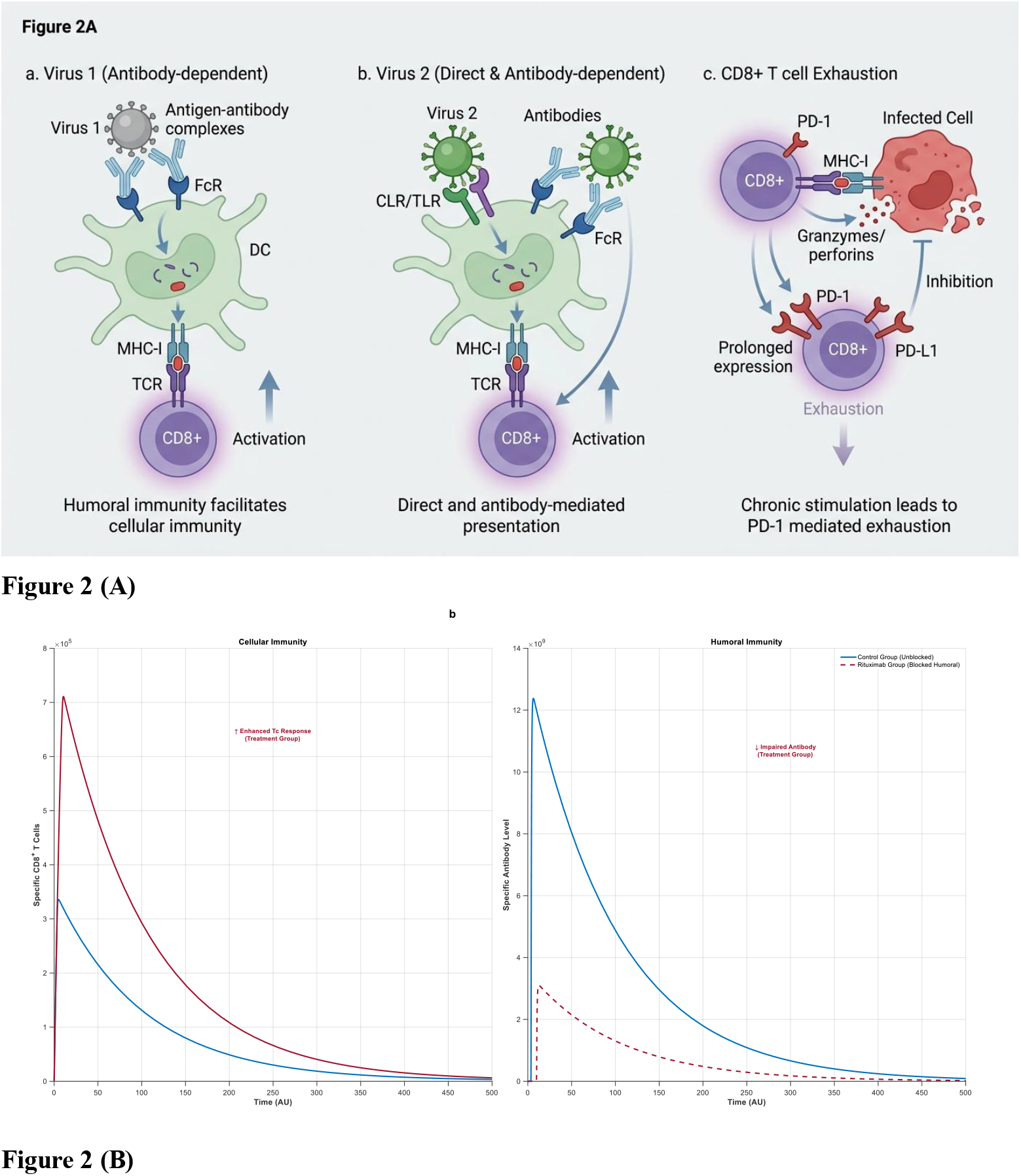
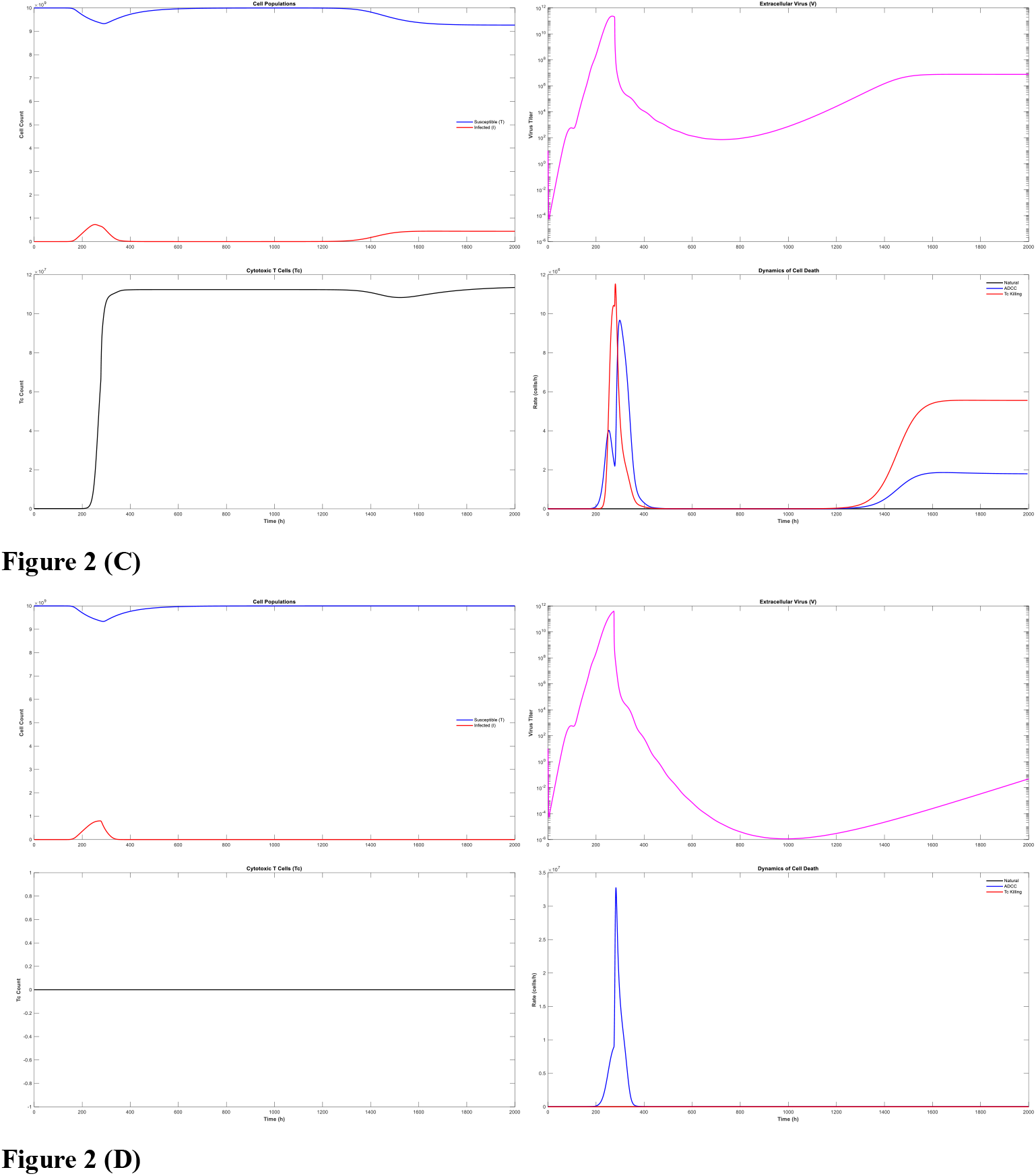
Mechanistic topology of cellular-humoral crosstalk and kinetics of T-cell mediated viral control. **(A)** Schematic of the integrated Cellular-Humoral Interface. The model defines two distinct activation pathways based on viral immunogenicity. Left (Virus 1, e.g., LCMV/Neoantigens): Illustrates an “Obligate Dependency” pathway where pathogens lack intrinsic DC-activating ligands (PAMPS). CTL priming is strictly gated by the formation of antigen-antibody immune complexes (ICs) and subsequent Fc-receptor mediated cross-presentation. Right (Virus 2, e.g., SARS-CoV-2): Illustrates a “Redundant” pathway where heavy surface glycosylation/PAMPs trigger direct DC activation (via TLRs/CLRs), operating in parallel with IC-mediated priming. The bottom panel depicts the T-cell Exhaustion module, modeling the progressive upregulation of inhibitory receptors (e.g., PD-1) and functional decay proportional to cumulative antigen stimulation. **(B)** Bifurcation of T cell Responses under B-cell Depletion. Simulation of CD8^+^ T cell magnitude in the presence (control) vs. absence (depletion) of B cells/Antibodies. For Virus 1 (low immunogenicity), B cell depletion collapses the T cell response (blue vs. red lines), conforming to the obligate dependency model. For Virus 2 (high immunogenicity), B cell depletion preserves or paradoxically enhances the T cell response (purple vs. green lines), consistent with the removal of antigen-sequestering antibodies in a redundant activation system [52]. **(C)** Cellular immunity predisposes the system to chronic infection by limiting antigen availability. Dynamics of infection in the presence of CD8+ T cell-mediated immunity with a weak antibody regeneration coefficient (equal to 2). Top panel: Temporal evolution of susceptible and infected cell populations, showing a progression from acute infection to a latent phase, followed by a transition to chronicity. Second panel: Kinetics of free virus (*V*). Note that while the presence of cytotoxic T lymphocytes (CTLs) suppresses the peak viral load (lowering acute severity compared to Figure 2D), the viral load rebounds after latency, settling into a lower-level steady state (chronic infection). Third panel: Proliferation kinetics of antigen-specific CD8+ T cells. Bottom panel: Decomposition of cell death mechanisms over time, showing the rates of antibody-dependent cellular cytotoxicity (ADCC), CTL-mediated lysis, and natural viral lysis. A rebound in lysis rates is observed in the late phase, consistent with persistent infection. **(D)** Ablation of cellular immunity enables viral clearance despite increased acute severity. Dynamics of infection in the complete absence of CD8+ T cells (*Tc* regeneration rate = 0; initial *Tc* = 0). Top panel: Susceptible and infected cell populations demonstrate a trajectory toward complete recovery following the acute phase. Second panel: Free virus kinetics. The absence of early T cell-mediated control results in a significantly higher peak viral load compared to (Figure 2C) ; however, this facilitates a robust humoral response leading to complete viral clearance (*V*→0). Third panel: Absence of CD8+ T cell proliferation (flatline at 0). Bottom panel: Cell death dynamics showing only ADCC-mediated and natural lysis events, which approach zero as the infection is cleared, contrasting with the persistent lysis observed in the chronic scenario.

#### Obligate vs. Redundant Activation Pathways

We first modeled pathogens with low intrinsic immunogenicity (“Virus 1”), typified by LCMV or poorly glycosylated tumor neoantigens that evade direct recognition by C-type lectin (CLR) or Toll-like receptors (TLR). For this class, we posit that humoral immunity acts as an obligate “gating” mechanism for cellular activation. In our model, the formation of antigen-antibody immune complexes induces Fc region conformational changes, which are requisite for Fc-receptor mediated uptake by DCs and subsequent cross-presentation via MHC-I to CD8^+^ T cells. This topology creates a critical dependency: failure in the humoral arm precludes effective cytotoxic T cell (CTL) priming. This mechanistic coupling provides a mathematical derivation for empirical observations where antibody depletion abrogates CD8^+^ T cell responses in LCMV models [49], and conversely, explains how B-cell depletion therapies ameliorate autoimmune pathology by dampening autoantibody-dependent CTL cytotoxicity [50–51].

In contrast, highly immunogenic pathogens (“Virus 2,” e.g., SARS-CoV-2) possess extensive surface pattern recognition motifs (PAMPs) that trigger DCs directly. This creates a “redundant” topology where CTL priming drives parallelly through direct viral recognition and Fc-mediated uptake. Consequently, our simulations predict that humoral blockade in this context does not abolish cellular immunity. Strikingly, under specific parameters **(Model 3.2.2)**, B-cell deficiency may paradoxically amplify T cell responses due to reduced antigen sequestration by antibodies, a phenomenon recapitulating clinical reports of robust T cell expansion in Rituximab-treated patients’ post-vaccination [52] (Figure 2B).

#### Temporal Dynamics of Cytotoxicity and the “Immune Eclipse” Phase

Beyond activation, effector function is governed by the interplay between TCR affinity and the spatiotemporal density of peptide–MHC-I complexes. Unlike the static neoantigen landscape of tumor cells, viral infection imposes a time-dependent antigen accumulation. We employed Model

#### 3.2.1 to simulate this non-linear progression

Critically, our model identifies an “immune eclipse phase” during early infection, where intracellular viral load is sufficient for replication but insufficient to generate surface MHC-I densities required for CTL recognition. This delay is a key determinant of infection outcomes. We demonstrate that chronicity arises from a kinetic mismatch: if infected cells lyse and release virions before neutralizing antibody titers peak, the released virus infects new targets which immediately enter the eclipse phase, rendering them temporarily invisible to CTLs. This cycle creates a self-sustaining reservoir of infection.

Cellular immunity attenuates acute viremia but compromises humoral-mediated viral clearance. To delineate the mechanistic interplay between cellular and humoral responses, we simulated viral dynamics under varying immune contexts. We observed a critical trade-off where CD8+ T cell-mediated cytotoxicity controls infection severity at the expense of antibody quality. In the presence of functional cellular immunity (**Fig. 2C**), the rapid lysis of infected cells significantly suppresses the peak viral load compared to T cell-deficient conditions. However, this early containment restricts the antigenic threshold required to drive robust clonal expansion and affinity maturation of B cells. Consequently, the host fails to achieve sterilizing immunity, resulting in a transition from latency to a rebound of viral replication, establishing a stable chronic infection characterized by persistent T cell proliferation and ongoing cytolysis. Conversely, the ablation of the CD8+ T cell compartment (**Fig. 2D**) leads to a significantly higher acute viral peak; yet, this elevated antigen availability triggers a potent humoral response that successfully drives the system toward complete viral clearance. These results suggest that while cellular immunity is essential for mitigating acute pathology, its suppressive effect on antigen load can paradoxically hinder the establishment of the sterilizing humoral immunity necessary to prevent chronic infection.

### 2.3 Synergistic Principles of B-T Cell Cognate Interactions and Memory Maintenance Kinetic Amplification via Cognate Help

Adaptive immunity fundamentally relies on the cognate interaction between antigen-presenting B cells and helper CD4^+^ T cells. We systematized these interactions into a state-space model encompassing eight distinct cellular contexts defined by B-cell isotype (IgM/IgG), antigen source (environmental/viral), and T-cell differentiation state (effector/memory) (Figure 3A; see Model 3.3.1).

**Figure 3.**
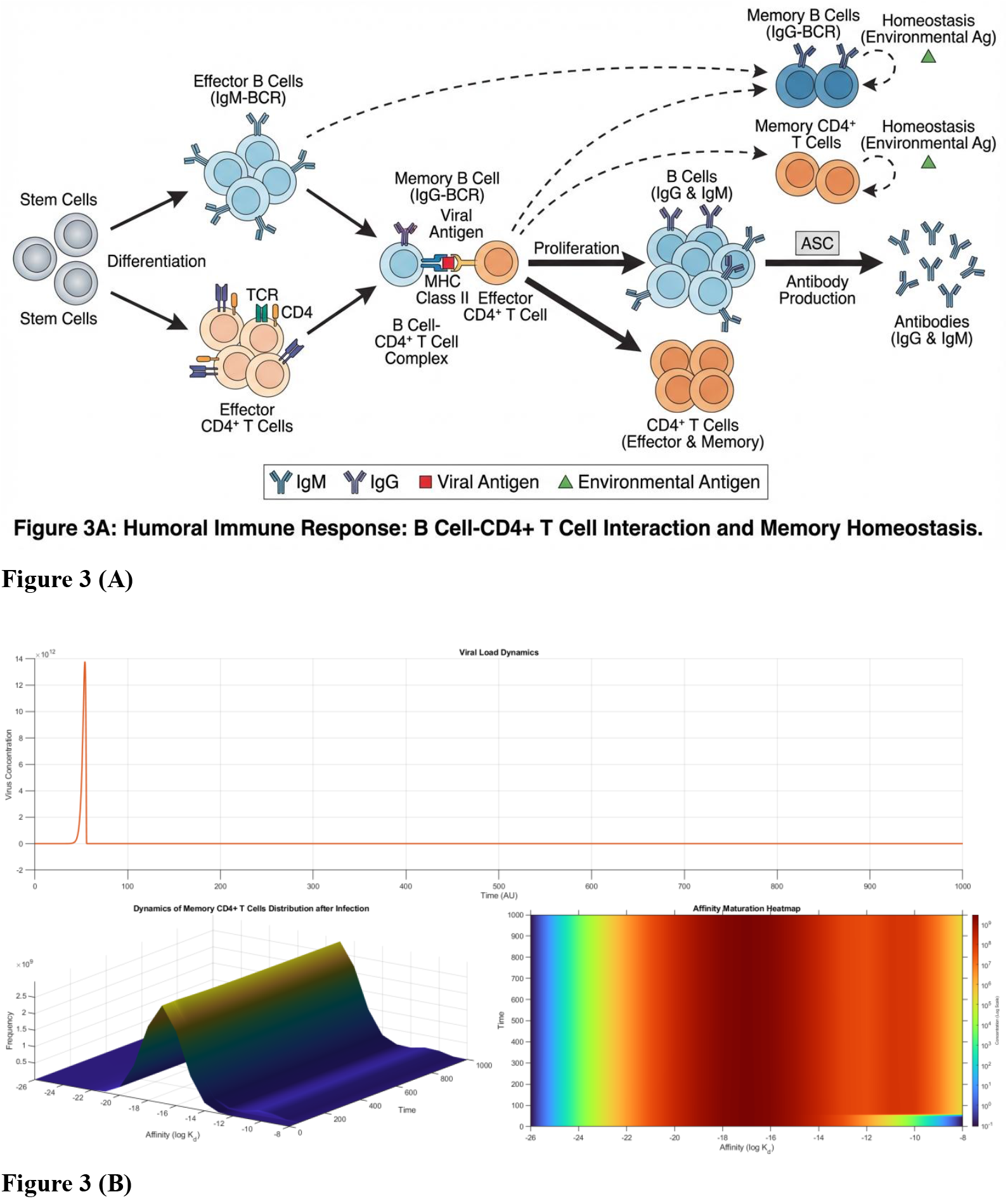
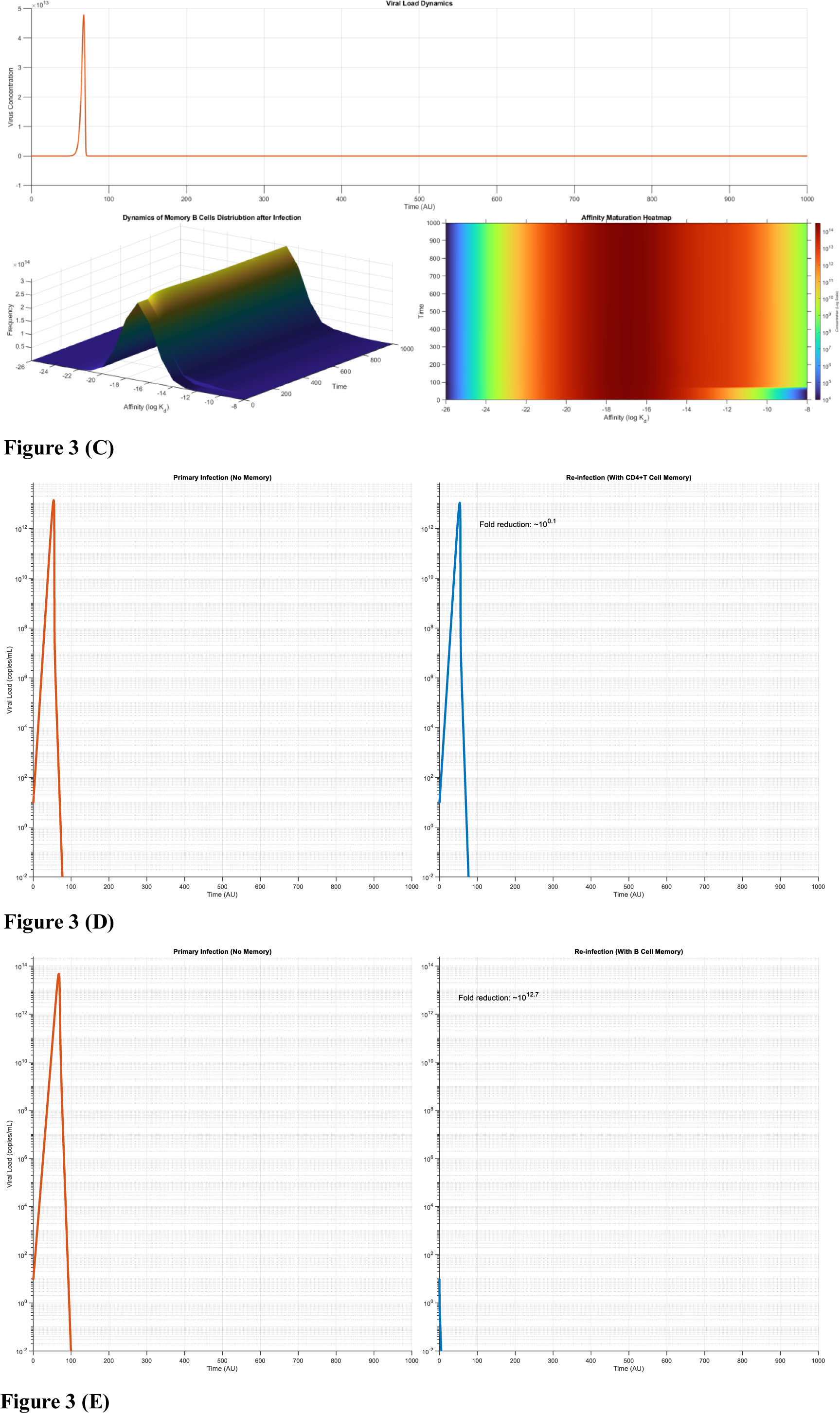
Principles of B-T cell synergy, repertoire evolution, and the functional dichotomy of immune memory. **(A)** Topology of Cognate Interactions. Schematic of the eight-compartment interaction matrix modeling antigen presentation. Contexts are defined by B-cell lineage (naïve IgM vs. memory IgG), T-cell state (effector vs. memory), and antigen source (viral vs. tonic environmental signals). This framework underpins the “Supra-linear Amplification” law, where ternary complex formation accelerates antibody production to the power of ∼1.2. **(B–C)** Repertoire Evolution Landscapes. Heatmaps illustrating the temporal evolution of receptor binding affinities post-infection. **(B)** TCR Diversity Dominance: Simulation with high TCR/low BCR diversity shows robust selection of high-affinity CD4^+^ clones. **(C)** BCR Diversity Dominance: Simulation with high BCR/low TCR diversity demonstrates the establishment of a persistent high-affinity IgG pool (Memory), compared to the transient high-affinity peak of IgM (Acute). Note: Both memory compartments are maintained by background tonic signaling from environmental antigens. **(D)** Efficacy of CD4^+^ T Cell Memory (Broad but Non-Sterilizing). Viral load kinetics upon rechallenge when only T cell memory is preserved. The response significantly reduces peak viral load (attenuation) via accelerated helper functions but fails to prevent productive infection. This mirrors the clinical phenotype of “breakthrough infections” with mild severity. **(E)** Efficacy of B Cell Memory (Specific and Sterilizing). Viral load kinetics upon rechallenge when only B cell memory is preserved. The presence of high-affinity BCRs/Antibodies completely suppresses viral replication (Sterilizing Immunity). Synthesis: The comparison between (D) and (E) illustrates the trade-off between high-potency/low-breadth (Humoral) and moderate-potency/high-breadth (Cellular) protection against drifting pathogens.

While previous models often treat T-cell help as a binary gatekeeper, our simulations reveal a quantitative law of “Supra-linear Kinetic Amplification.” In a B-cell-autonomous system (Model 3.1.1), antibody output scales linearly with antigen–BCR complex density. However, the integration of ternary complexes (Antigen–BCR–TCR) introduces a non-linear scalar, amplifying production approximately to the power of 1.2. This kinetic boost is not merely an accelerator but a functional requirement: in our agent-based simulations, this amplification is the critical determinant that pushes the system past the “sterilizing threshold.” Without this CD4^+^ synergy, the immune response consistently fails to outpace viral replication, causing the system to bifurcate toward a chronic infection state rather than clearance.

#### Divergent Repertoire Evolution (Figure 3B–C)

To disentangle the stochastic evolution of lymphocyte repertoires, we simulated scenarios isolating TCR constraint (High BCR/Low TCR diversity) versus BCR constraint (Low BCR/High TCR diversity). A universal topological feature emerged: both IgG and CD4^+^ T cell compartments evolve toward persistent high-affinity peaks, establishing the basis of long-term memory. Crucially, we identified that memory maintenance for both lineages shares a common mechanistic dependency on tonic signaling derived from environmental antigens. In contrast, the IgM compartment, despite forming transient high-affinity clusters, undergoes rapid decay due to the lack of self-renewal capacity, confirming its role as a “first-responder” wave rather than a memory reservoir.

Functional Dichotomy: Sterilizing Specificity vs. Broad Mitigation.

We next interrogated the distinct protective roles of humoral versus cellular memory by simulating secondary challenges under conditions of selective memory preservation.

B-Cell Memory (The Sterilizing Wall): Preservation of the post-infection B-cell repertoire confers “sterilizing immunity” against homologous viral strains. Due to the immediate availability of high-affinity antibodies, viral replication is suppressed instantaneously (Figure 3E).

T-Cell Memory (The Mitigation Buffer): In contrast, preservation of CD4^+^ T-cell memory alone does not prevent re-infection. Instead, it accelerates the de novo antibody response via cognate priming, resulting in a reduced peak viral load and attenuated disease severity (Figure 3D).

#### The Structural Trade-off: Fragility vs. Conservation

Our model provides a mechanistic rationale for the clinical observations seen in rapidly evolving pathogens like SARS-CoV-2 [53–55]. B-cell memory targets tertiary conformational epitopes, which are highly susceptible to structural disruption by single point mutations (antigenic drift). Consequently, B-cell mediated protection is “fragile”—highly potent against homologous strains but essentially abrogated by significant drift. Conversely, CD4^+^ T cells recognize linear peptide sequences derived from intracellular processing, which are evolutionarily conserved across variants. This structural dichotomy creates a fundamental trade-off: Humoral memory offers sterilizing but strain-specific protection, while cellular memory provides non-sterilizing but cross-reactive breadth. This explains why vaccine protection against infection (humoral) wanes rapidly with viral drift, while protection against severe disease (cellular) remains robust [56–58].

#### Implications for Vaccine Design

These findings necessitate a paradigm shift in evaluating vaccine efficacy for mutating viruses. “Protection duration” is an insufficient metric; the quality of “imprinted immunity” is paramount. We propose that optimal vaccines must leverage B-cell antigens not solely for antibody production, but to reshape the upstream CD4^+^ helper landscape. However, we also identify a critical limitation in therapeutic settings: high-affinity circulating antibodies can sequester B-cell antigens, blocking BCR access and dampening the very memory reactivation they are intended to boost—a mechanism (“Isotope Masking”) that we explore further in the context of cancer neoantigen vaccines (Section 5).

### 2.4 Integrated Agent-Based Modeling: Spatiotemporal Bifurcation and Therapeutic Thresholds

To resolve the kinetic discrepancies between serum viral loads and tissue-level pathology, we engineered a discrete agent-based framework that introduces a critical topological constraint: spatiotemporal shielding. As delineated in Figure 4A, we model infection as a discontinuous process where intracellular viral replication is temporarily sequestered from neutralizing antibodies. Lysis events—triggered either by reaching a viral capacity threshold or via immune-mediated cytotoxicity (ADCC and CD8^+^ CTL)—release virions into the extracellular milieu, creating a bottleneck where virions must compete between antibody neutralization and de novo infection of susceptible cells.

**Figure 4.**
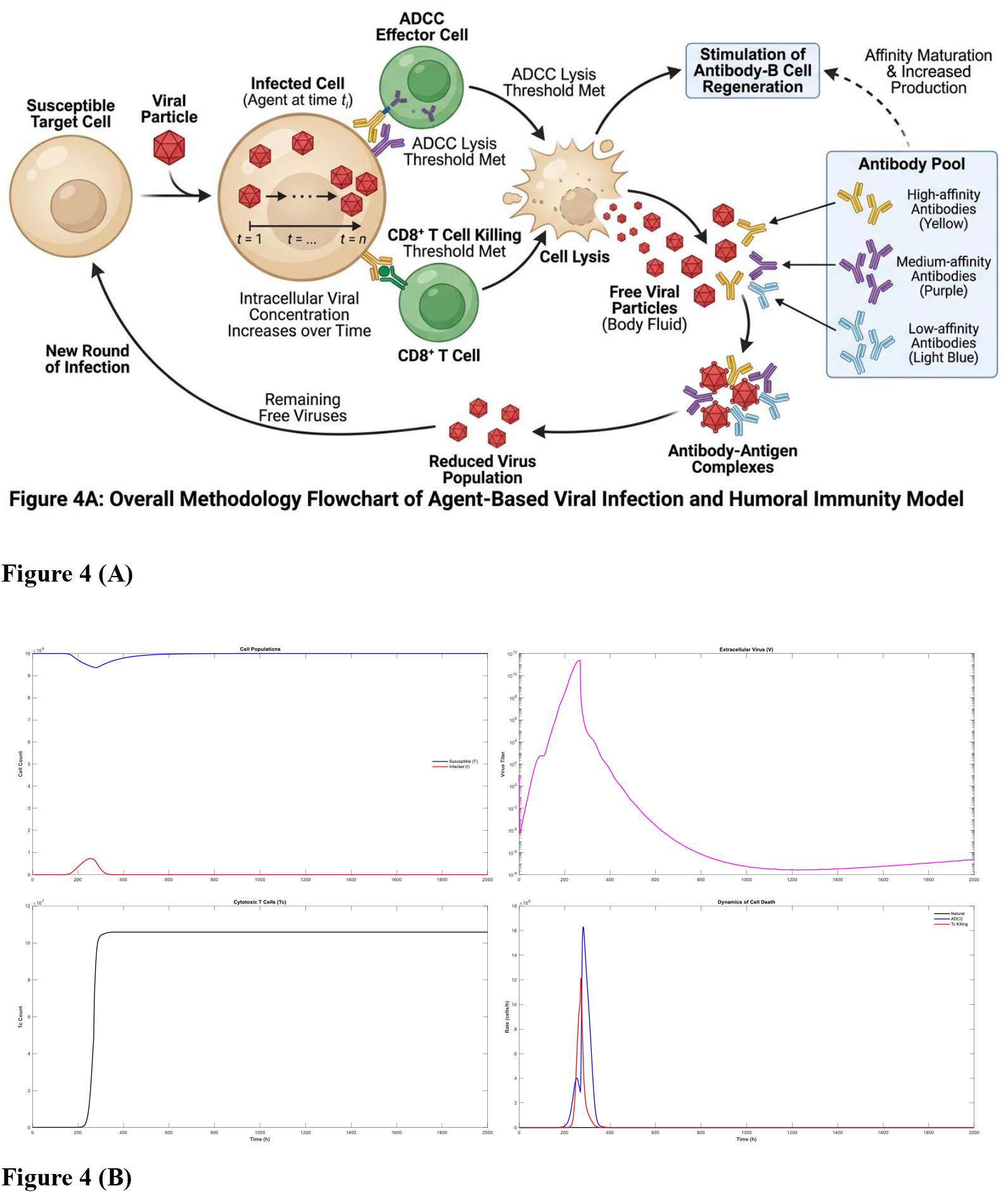
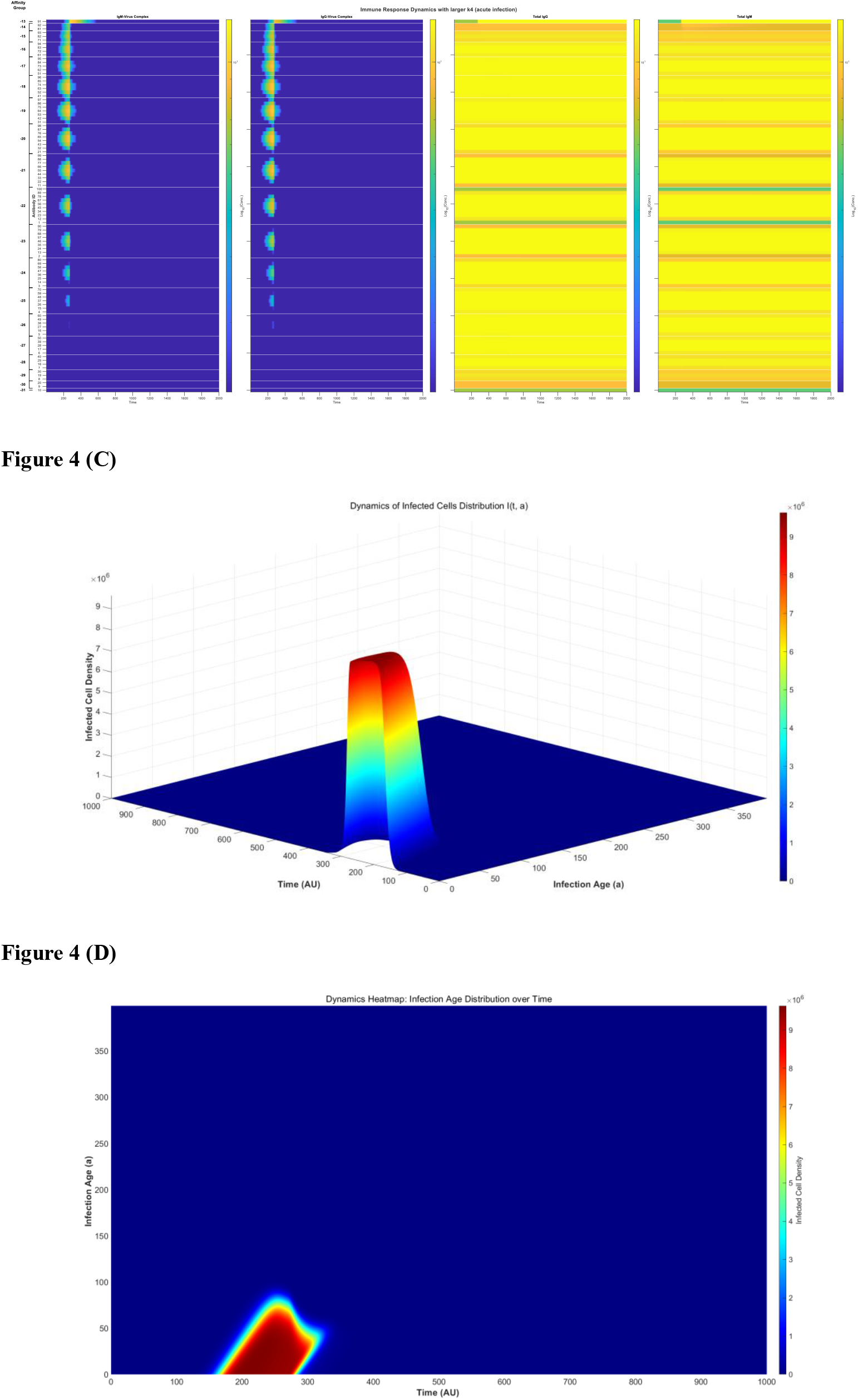
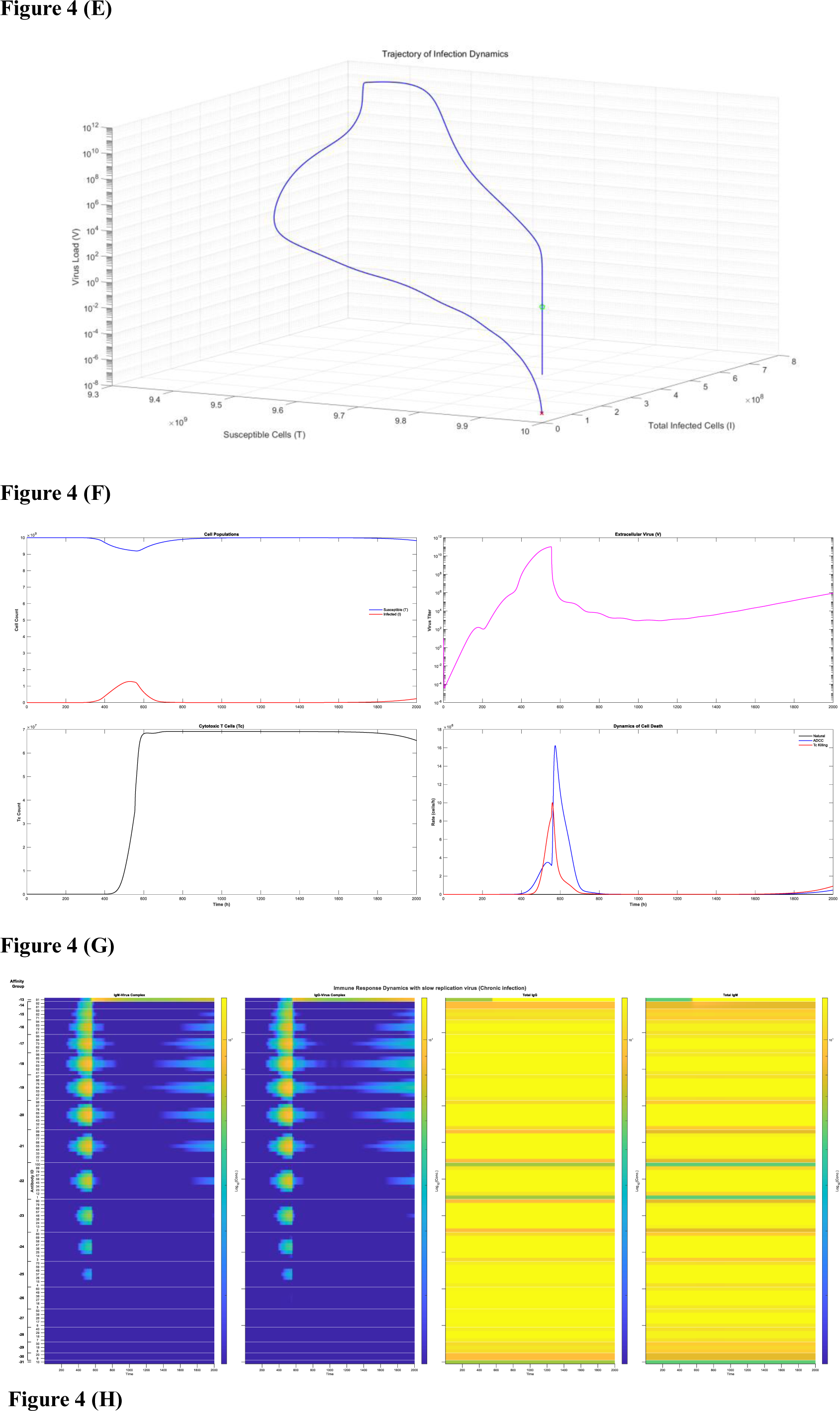
Agent-based simulation of infection bifurcation and non-linear therapeutic responses. **(A)** Schematic representation of the stochastic viral infection and antibody affinity maturation model. Flowchart illustrates the iterative dynamics between viral replication and the humoral immune response. The model initiates with a viral population (red particles) encountering a polyclonal antibody pool characterized by varying binding affinities: high (yellow), medium (purple), and low (blue). In the neutralization phase, antibodies bind to viruses based on their affinity; high-affinity binding effectively neutralizes viral particles, while surviving viruses infect somatic cells (acting as protective compartments) to initiate a new replication cycle. Crucially, the formation of antigen-antibody complexes triggers a positive feedback loop, stimulating the selective proliferation of corresponding B-cell clones. This is modeled via a regeneration coefficient (e.g., a factor of 2, as depicted), which preferentially amplifies high-affinity antibodies. Over successive time steps (i*i*), this mechanism shifts the antibody landscape toward high-binding variants, ultimately achieving a threshold where circulating antibodies are sufficient to neutralize all viral progeny released from cell lysis, thereby terminating the infection. **(B)** Dynamics of viral eradication driven by high-affinity antibody responses. Simulation results for an individual with robust immunity (antibody regeneration coefficient increased from 2 to 3 compared to Fig. 2C). The panels illustrate the temporal evolution of (from left to right): susceptible (blue) and infected (red) cell populations showing no resurgence of infection; free virus load (V) demonstrating complete clearance without rebound; expansion of antigen-specific CD8+ T cells; and the rates of cell lysis mediated by ADCC, cytotoxic T cells (Tc), and viral burst. Note the absence of cytolytic activity in the late phase, indicating successful resolution of infection. **(C)** Kinetics of immune complex formation and free antibody levels. Temporal profiles of antibody-virus complexes (IgM-V, IgG-V) and free immunoglobulins. Early peaks in IgM-V and IgG-V complexes (left two panels) conform to viral load dynamics and decline upon clearance. The right panels demonstrate that while specific IgM levels are transient, high-affinity specific IgG is maintained at a high plateau, indicative of established protective immunity. **(D)** Three-dimensional distribution of infected cell population structural dynamics. The density of infected cells *I*(*t,a*) is plotted against time (*t*) and infection age (a, ranging from 1 to 400 AU). The data reveals that infected cells do not survive to advanced infection ages due to rapid clearance by ADCC and Tc responses. The absence of infected cells after *t*=350 confirms the termination of the acute infection phase. **(E)** Heatmap projection of bacterial infection age dynamics. A two-dimensional projection of the data in Figure 4 D, visualizing the density of infected cells with varying infection ages over time. The heat map clearly delineates the cessation of new infection events and the elimination of existing infected cells in the late stages of the immune response. **(F)** Phase space trajectory of complete viral clearance. A three-dimensional phase diagram illustrating the system’s trajectory from the initial state (green circle: V=0, Susceptible cells =10^10^, Infected cells =0) to the terminal state (red cross). The trajectory converges back to the virus-free equilibrium (V=0) with full recovery of susceptible cells, confirming that the system does not settle into a chronic steady state. **(G)** The ‘slow replication trap’ facilitates chronic infection establishment. Simulation results where the viral proliferation coefficient is reduced to 0.05 (compared to 0.1 in Fig. 4B), representing a slowly replicating virus (e.g., HBV-like kinetics). The panels show (from left to right): the temporal dynamics of susceptible and infected cell populations, where infected cells exhibit late-phase recrudescence rather than elimination; the rebound of free viral load (V) indicating the formation of a persistent chronic state; CD8+ T cell expansion; and the rates of cell lysis. Unlike the clearance observed in acute infection, significant cytolytic activities (ADCC, Tc) persist into the late phase, reflecting ongoing localized infection. **(H)** Dysregulated antibody kinetics and immune complex persistence. Temporal profiles of immune complexes (IgM-V, IgG-V) and free immunoglobulins (IgM, IgG) corresponding to the chronic scenario in Figure 4 G. In contrast to the resolution seen in Fig. 4C, immune complexes (left two panels) initially decline but display a significant secondary rise, mirroring viral resurgence. The right panels reveal that the induction of high-affinity specific IgG is markedly delayed compared to the acute response (Fig. 4C) and reaches a lower plateau, suggesting that slowly replicating viruses evade the threshold required to trigger a robust, sterilizing humoral response.

Crucially, this framework introduces an affinity-dependent positive feedback loop (λ=2). High-affinity antibodies (yellow agents) not only neutralize virions but form immune complexes that drive preferential B-cell clonal expansion. This creates a “winner-take-all” dynamic required for sterilizing clearance (Figure 4B-E). Integrating this spatiotemporal granularity reveals that chronicity is not merely a failure of clearance, but a quasi-stable equilibrium driven by specific kinetic failures. We identified three distinct bifurcation pathways toward this persistent state:

I. The Feedback Failure Trap (Figure 4B-E compared to Figure 2C). In hosts with intrinsic immune compromise (reduced feedback coefficient k4), the system fails to reach the critical antibody titer required to intercept all released virions. This establishes a “smoldering” state: a high burden of infected cells persists despite low extracellular viral loads, maintaining chronic T-cell stimulation that drives exhaustion [59–60].
II. The Imprinting Paradox (Extended Figure 4). We elucidate a counter-intuitive mechanism driving chronicity following reinfection with antigenic variants [64]. Pre-existing cross-reactive immunity suppresses the initial exponential phase of viral replication. Paradoxically, this “efficient” early control acts as a kinetic trap: it deprives the naïve B-cell compartment of the antigenic mass required to trigger robust de novo affinity maturation. The system effectively “starves” the adaptive response, stabilizing at a sub-threshold equilibrium implicated in long-COVID sequelae [61–63].
III. The “Stealth” of Slow Replication (Figure 4F-G). Contrary to the intuition that virulence correlates with persistence, our simulations identify slow viral replication kinetics as a potent driver of chronicity. Rapidly replicating pathogens (e.g., Influenza) trigger an immediate, high-magnitude immune alarm. Conversely, slow-replicating viruses (e.g., HBV, HCV) expand below the immune activation threshold, establishing a deep cellular reservoir before effector mechanisms are fully engaged [59–60].

#### Non-Linear Therapeutic Dynamics

We leveraged this platform to simulate perturbations, revealing that therapeutic responses in chronic settings are governed by strict thresholds and non-linearities:

- Small-Molecule Inhibitors: While effective for symptom control [65–67], inhibitors alone typically fail to achieve sterilization because they do not reverse the underlying immune dysregulation or threshold deficits.
- Threshold Effects in mAb Therapy (Extended Figure 5): Monoclonal antibody outcomes exhibit bistability. Sub-therapeutic doses provide transient suppression followed by rebound (hysteresis), as they fail to outlast the intracellular viral eclipse phase. Only high-dose regimens achieving “sterilizing saturation” can force the system out of the chronic attractor [68–70].
- Metastable Polyclonal Boosting (Extended Figure 6): Investigating physical therapies (e.g., Gua Sha) or peptidoglycan stimulation [71], we find that modulating environmental antigens induces a transient “polyclonal lift” in basal antibody titers. While this perturbs the system—potentially acting as an adjuvant—it typically returns to the chronic saddle point absent specific affinity maturation.

The “Antigen Sink” Effect in Therapeutic Vaccination (Extended Figure 7).

Most critically, our model identifies a dangerous non-monotonic dose-response in therapeutic vaccination [72–74]. In the chronic plateau phase, low-dose antigen administration acts as an “antibody sink”—sequestering existing neutralizing antibodies without surpassing the threshold to trigger B-cell clonal bursts. This paradoxically exacerbates viral load. Our model dictates that effective therapeutic vaccination requires a massive antigen bolus—significantly exceeding prophylactic doses—to break the tolerance plateau and drive high-affinity clone expansion.

### 2.5 The “Immunogenic Visibility” Hypothesis

While viral kinetics are well-characterized, the quantitative laws governing cancer-immune equilibration remain under-explored. We established a novel system of ordinary differential equations (ODEs) (Model 3.5.1) predicated on the axiom that immune recognition is strictly a function of “immunogenic visibility.”

Standard single-nucleotide polymorphisms (SNPs) often fail to breach the tolerance threshold due to poor proteasomal processing. In contrast, our model focuses on “high-quality” neoantigens derived from aberrantly expressed retroelements (e.g., HERVs) [75, 76] or frameshift mutations via alternative splicing [77, 78]. These sources generate structurally novel peptides that ensure robust presentation by both MHC-I and MHC-II, driving the synergistic activation of CD8^+^ cytotoxic T lymphocytes (CTLs) and antibody-dependent cellular cytotoxicity (ADCC).

#### The Theory of Pre-Equilibrium Expansion

A rigorous mathematical analysis of our framework (see Supplementary Materials) reveals a universal topological feature: the cancer-immune system inevitably tends toward a homeostatic equilibrium where malignant cells persist at subclinical levels. The clinical bifurcation between “health” and “disease” is therefore defined not by the final state, but by the magnitude of “Pre-equilibrium Expansion.”

- Dormancy: Under robust surveillance, cancer cells reach equilibrium with minimal expansion.
- Overt Disease: If surveillance is compromised or proliferation rates are high, the tumor undergoes massive expansion before stabilizing, manifesting as clinical cancer. This theoretical insight shifts the therapeutic goal from “eradicating equilibrium” to “attenuating the pre-equilibrium trajectory” (Figure 5B).

**Figure 5.**
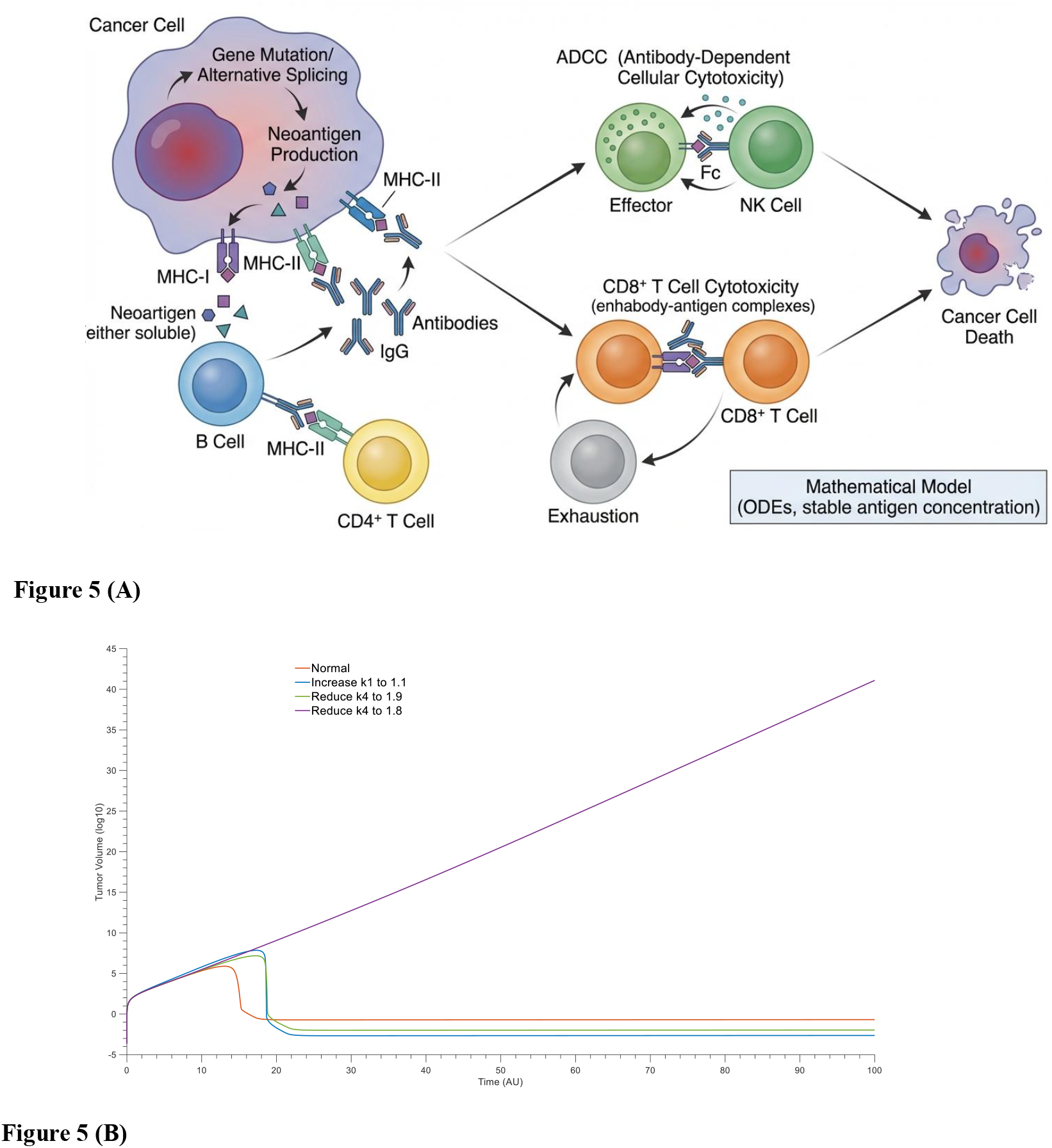
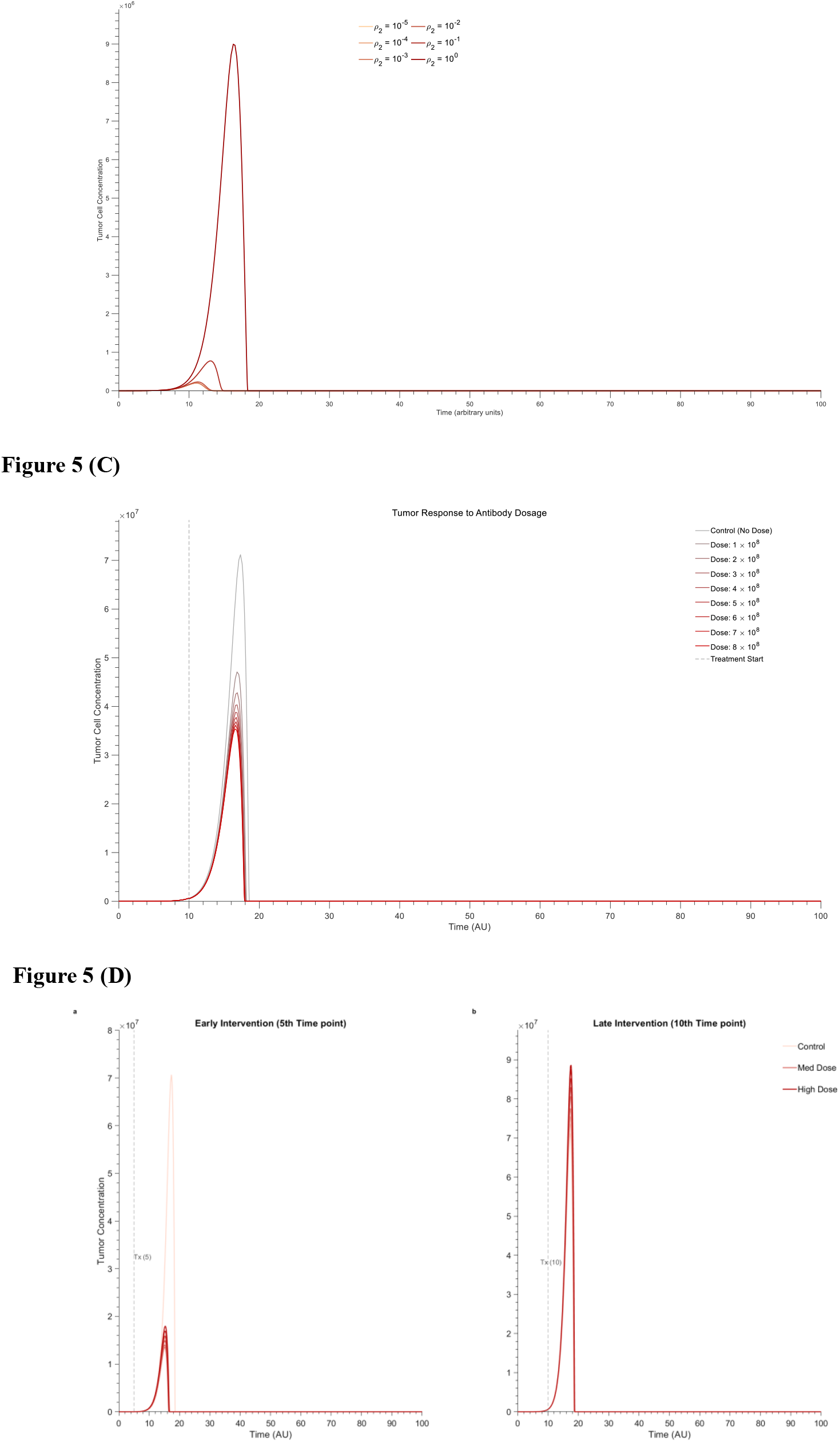
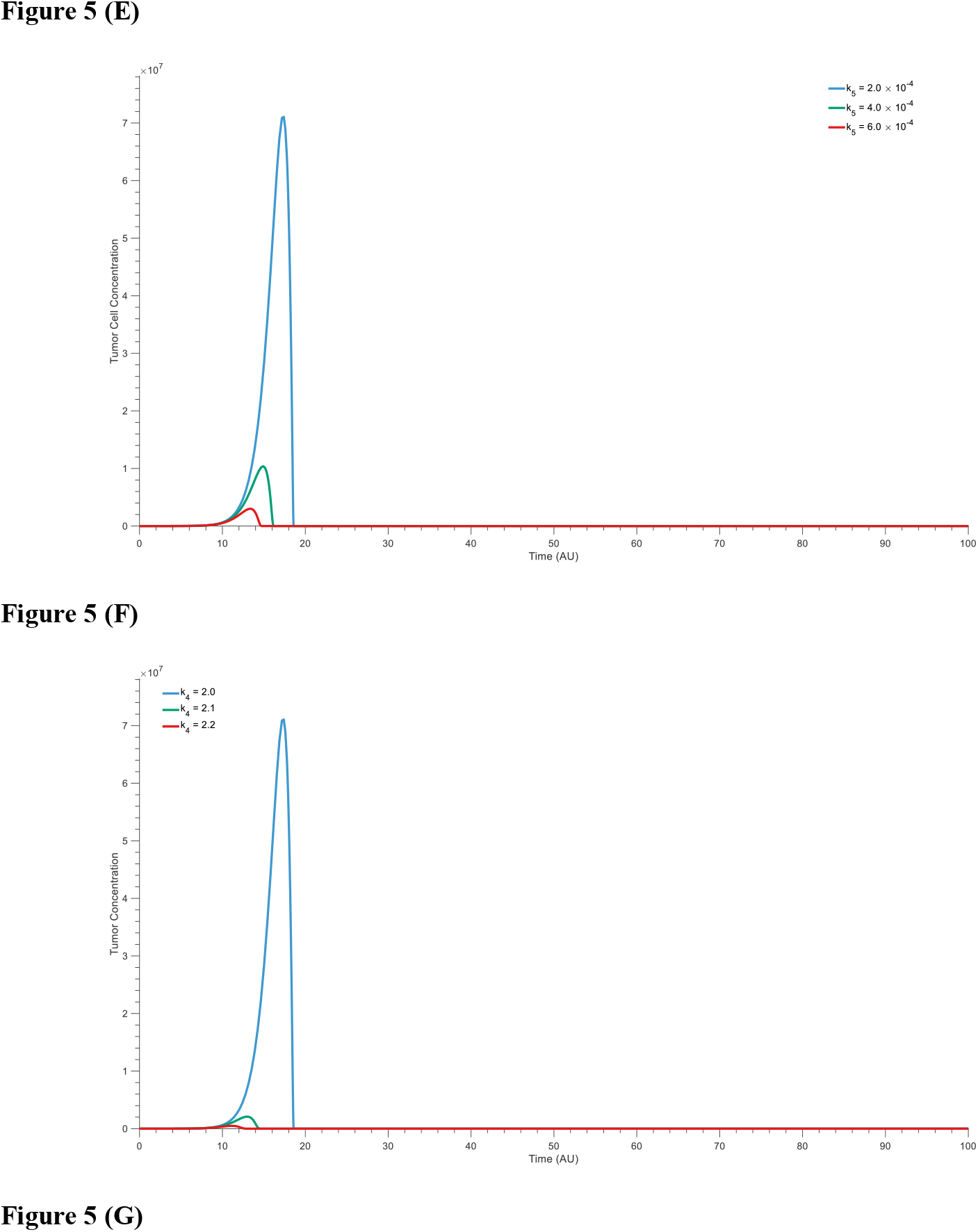
Dynamics of the Cancer-Immune Equilibrium and Mechanism-Based Evaluation of Immunotherapies. **(A)** Topology of the Cancer Immunity Model. Schematic flow integrating dual effector arms: (1) CD8^+^ T cell-mediated lysis modulated by exhaustion (PD-1), and (2) Antibody-Dependent Cellular Cytotoxicity (ADCC). Both arms are coupled via antigen-antibody complex feedback loops. **(B)** The “Pre-Equilibrium Expansion” Theory. Phase-plane trajectories illustrating how clinical outcomes are determined. • Blue Line (Robust Immunity): Cancer cells stabilize rapidly with minimal expansion (Subclinical/Dormant). • Red Line (Compromised Immunity): Weak surveillance allows massive expansion before reaching equilibrium, crossing the “Clinical Detection Threshold” (Overt Cancer). **(C–D)** Standard Interventions. **(C)** PD-1 Blockade: Reducing T-cell exhaustion frequency significantly suppresses tumor growth. **(D)** Neoantigen-Specific mAb: Passive inputs of antibodies drive rapid clearance via ADCC, though efficacy depends on target density. **(E)** The “Antigen Sink” Paradox (B-Cell Antigens). Simulation of whole-antigen/lysate vaccination. Early/Low Dose: Effective control. Late/High Dose: Paradoxical tumor acceleration. Mechanism: Excessive free antigen sequesters circulating antibodies (the “Sink” effect), reducing the effective concentration available for ADCC at the tumor site. **(F–G)** The Advantage of MHC-Restricted Vaccines. **(F)** MHC-I Peptides: Enhance CD8^+^ CTL magnitude directly without consuming antibodies. **(G)** MHC-II Peptides: Boost T-helper function and downstream antibody production coefficient (k_5_). Key Takeaway: Unlike B-cell antigens, MHC-restricted peptides decouple immune stimulation from antibody consumption, avoiding the therapeutic resistance observed in (E).

Therapeutic Simulation: Checkpoints and Antibodies.

We validated the model by simulating established interventions.

- Checkpoint Blockade: By reducing the exhaustion parameter (ρ2ρ2), PD-1 inhibitors mitigate the self-inactivation of CTLs. Our simulations recapitulate clinical data [79], showing that rescuing CTL exhaustaion effectively suppresses pre-equilibrium expansion (Figure 5C).
- Monoclonal Antibodies (mAbs): Targeting dominant neoantigens (e.g., HERV-derived) enhances clearance via a dual mechanism: direct ADCC and the feedback amplification of CTLs via immune complex formation [80, 81]. However, we note that this strategy is economically constrained by the need for highly expressed, shared targets, limiting its personalized application (Figure 5D).

#### The “Antigen Sink” Paradox in B-Cell Vaccines

A critical, counter-intuitive finding emerges from our simulation of B-cell antigen vaccines (e.g., tumor lysates or whole proteins). While effective at low doses in early-stage disease, high-dose administration in advanced stages paradoxically accelerates tumor growth [82]. Our model identifies an “Antigen Sink” mechanism: excessive circulating B-cell antigens sequester neutralizing antibodies away from the tumor surface, thereby abrogating ADCC. This non-monotonic dose-response implies that late-stage administration of whole-protein vaccines may inadvertently shield the tumor from humoral cytotoxicity (Figure 5E).

MHC-Restricted Vaccines: Decoupling Activation from Consumption.

To circumvent the “Antigen Sink,” we simulated strategies targeting MHC-I and MHC-II restricted epitopes.

- MHC-I Peptides: These small ligands bind directly to dendritic cells to boost CTLs [83– 85] but do not engage circulating antibodies. This “decoupling” avoids antibody consumption, creating a pure positive feedback loop (Figure 5F).
- MHC-II Peptides: Investigating helper epitopes, we found significant efficacy [86–88]. By stimulating CD4^+^ help, these antigens enhance downstream antibody production (increasing k_5_) without acting as a sink. The resulting surge in titers amplifies ADCC and subsequent antigen release, establishing a robust feed-forward cycle largely largely devoid of the risks associated with whole-antigen therapies (Figure 5G).

## Methods

### 3.1 Humoral immunity model in adaptive immunity

#### 3.1.1 Kinetic Framework of Viral-Antibody Interaction

To elucidate the fundamental dynamics of humoral immunity, we established a governing system of ordinary differential equations (ODEs) describing the interplay between viral load (V), free antibodies (A), and virus-antibody immune complexes (C). The basal model (Model 3.1.1) is defined as follows:

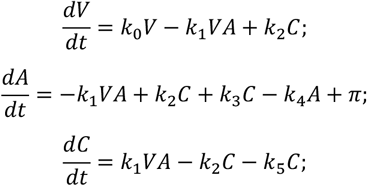

Here, k_0_ denotes the viral replication rate. The association and dissociation rate constants for antibody-virus binding are represented by k_1_ and k_2_, respectively. The term k_3_ quantifies the antigen-dependent stimulation of antibody production (feedback proliferation induced by immune complexes), while k_4_ represents the natural decay rate of free antibodies. The constant π denotes the constitutive rate of antibody production (flux from long-lived plasma cells or background synthesis), and k_5_ is the clearance rate of immune complexes.

#### 3.1.2 Incorporation of Environmental Antigens for Immunological Memory

A critical limitation of the basal framework (Model 3.1.1) is its inability to sustain immunological memory. Specifically, in the absence of viral antigen, the system reverts to its initial equilibrium; despite transient proliferation during infection (driven by the −k_4_A term relative to π), specific antibody titers do not stabilize at elevated levels post-clearance. This trajectory contradicts the established biological phenomenon of long-term memory maintenance.

To address this, we extended the model to incorporate environmental or self-antigens (E), which provide the necessary survival signals for memory maintenance via cross-reactivity or homeostatic tick-over. The refined model (Model 3.1.2) is expressed as:

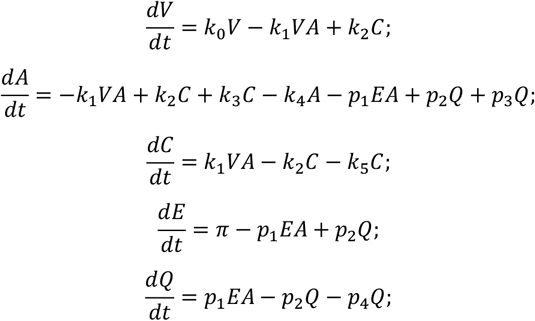

In this formulation, E represents environmental or self-antigenic substrates, and Q denotes the complex formed between antibodies and environmental antigens. The parameter p_1_ is the association rate for the *E*-*A* interaction, p_2_ is the dissociation rate, and p_3_ represents the coefficient of antibody regeneration stimulated by the Q complex. The term p_4_ denotes the degradation rate of the Q complex. This modification allows the system to exhibit bistability or elevated equilibrium states characteristic of immune memory.

#### 3.1.3 Clonal Selection and Affinity Maturation

To validate the mechanism by which the immune system selects for high-affinity clones and maintains diversity, we generalized the core model to simulate a polyclonal response. This framework allows for the analysis of antibody selection under competitive constraints, viral rebound phenomena, and dynamics during therapeutic interventions. The dynamics of immunological screening are modeled as (Model 3.1.3):

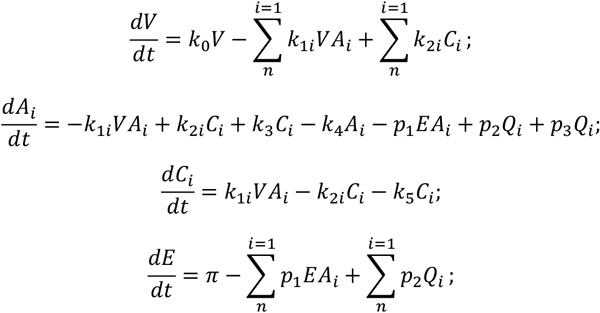

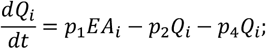

Where subscript *i* denotes a distinct antibody clone. The parameters 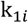 and 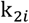represent the clone-specific association and dissociation rate constants with the virus, respectively. The equilibrium dissociation constant is defined as 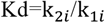, where higher affinity correlates with a larger 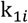 and smaller 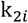. We assume that interactions with environmental antigens ( E) are generic; thus, the binding parameters (p_1_, p_2_) and homeostatic turnover rates (p_3_, p_4_) are constant across clones. This model captures the kinetic evolution of the antibody repertoire, demonstrating not only the acute proliferation of specific clones but also the shift in clonal composition—the mechanism underlying the maintenance of long-term memory quality.

#### 3.1.4 Viral Rebound under Monoclonal Antibody Therapy

Finally, we modeled the dynamics of viral rebound following passive immunization. This scenario introduces an exogenous monoclonal antibody (A_2_) alongside the endogenous response:

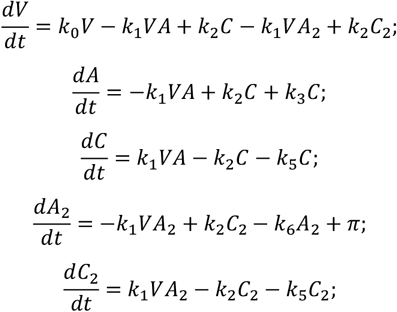

Here, A_2_ represents the therapeutic monoclonal antibody. For comparative rigor, A_2_ is assigned kinetic parameters (k_1_, k_2_) identical to the host’s neutralizing antibodies. Crucially, the exogenous complex (C_2_) does not induce B-cell selection or antibody regeneration (absence of the k_3_C term for A_2_), and free exogenous antibodies are subject to a distinct clearance rate ( k_6_). The term π(t) represents the dosing function: π(t)>0 at the moment of injection and π(t)=0 otherwise.

#### Model 3.1.5: Dynamics of Viral Replication Inhibitor Therapy

To investigate the efficacy of small-molecule antivirals compared to antibody-based interventions, we developed a pharmacodynamic model describing the interaction between the virus (V), antibodies (A), and a specific integration inhibitor (D). The governing equations (Model 3.1.5) are as follows:

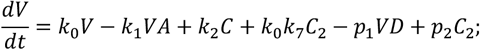

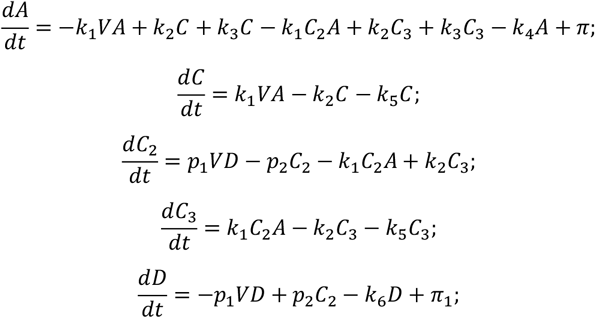

In this compartmental framework, D denotes the concentration of the small-molecule inhibitor, and C_2_ represents the virus-inhibitor complex (V-D). We explicitly model the formation of a ternary complex, C_3_, formed by the binding of antibodies to the virus-inhibitor complex (A-V-D).

The kinetic parameters are defined as follows: p_1_ and p_2_ represent the association and dissociation rate constants for the inhibitor-virus interaction, respectively. The parameter k_7_ ∈[0,1] is a dimensionless coefficient representing the residual replicative capacity of the virus-inhibitor complex (C_2_), quantifying the drug’s inhibitory efficiency (where k_7_=0 implies complete inhibition). The complexes C and C_3_ share a standard immune clearance rate k_5_. The intrinsic decay rates for the antibody and the small molecule are denoted by k_4_ and k_6_, respectively. The term π_1_ represents the dosing function for the inhibitor: it equals the administered dose at the moment of injection and is zero otherwise.

#### Model 3.1.6: Cellular Mechanism of Antibody Production

To provide a mechanistic understanding of the humoral response beyond phenomenological formulations, we expanded the antibody dynamics to explicitly model the differentiation of B cells into Antibody-Secreting Cells (ASCs). This granular model (Model 3.1.6 ) allows for the differentiation between surface-bound receptors (BCRs) and secreted immunoglobulins:

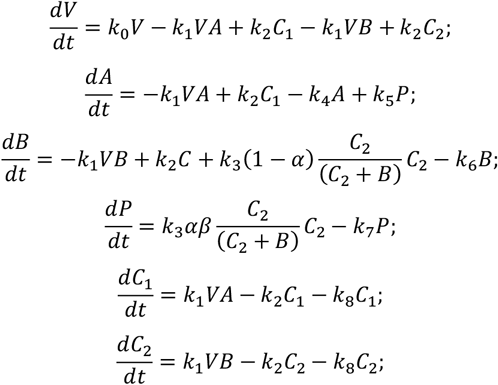

Here, P denotes the population of ASCs (plasma cells), and B represents the density of B-cell receptors (BCRs) on non-plasma B cells, serving as a proxy for the antigen-specific B-cell population. C_1_ and C_2_ represent the Virus-Antibody complex and the Virus-BCR complex, respectively.

The proliferation and differentiation dynamics are governed by the antigen-mediated activation term, where k_3_ is the clonal expansion rate derived from BCR signaling. The term (C_2_ + B) models the saturation of receptor occupancy or signaling efficiency. The parameter α defines the fraction of activated B cells differentiating into ASCs, while β is a scaling conversion factor between BCR density and cell number. The secretion rate of antibodies by ASCs is given by k_5_. k_6_ and k_7_ denote the apoptotic rates for non-plasma B cells and ASCs, respectively. Finally, k_8_ represents the degradation rate of immune complexes ( C_1_ and C_2_).

Regarding the viral parameter k_0_: in the context of natural infection, k_0_>0 represents the viral replication rate. Conversely, in vaccination scenarios using non-replicating antigens, k_0_ is assigned a negative value to mathematically represent the natural decay of the antigen bolus.

#### 3.1.7 Modeling Isotype Switching, Differentiation, and Repertoire Diversity

To capture the full temporal complexity of the humoral response, we expanded the model to distinguish between immunoglobulin isotypes (IgM vs. IgG) and cellular states (naïve/memory B cells vs. Antibody-Secreting Cells, ASCs). This high-fidelity framework (Model 3.1.7) incorporates class switching, affinity-dependent activation, and the structural differences between isotypes (e.g., pentameric IgM vs. monomeric IgG).

##### 1. Cellular Dynamics of B-Cell Receptors (BCR)

We model the dynamics of specific B-cell clones, indexed by *i*, identifying IgM-expressing 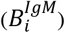 and IgG-expressing 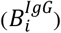 populations.

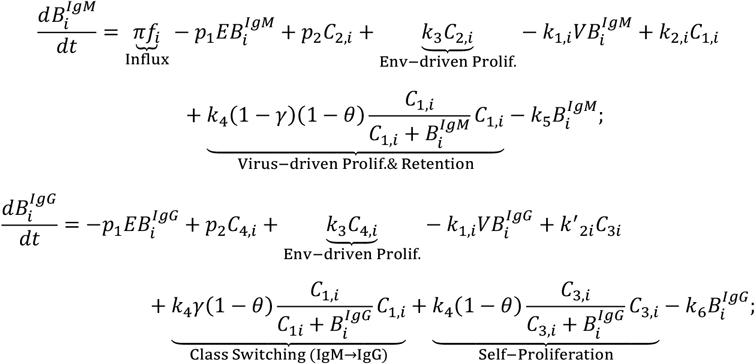

Here, π represents the total homeostatic influx of naïve B cells, distributed across the repertoire according to *f*_*i*_, which follows a Gaussian distribution based on binding affinity. The parameter γ governs the rate of isotype class switching from IgM to IgG, while θ represents the fraction of activated B cells differentiating into terminal ASCs. The saturation term 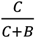 highlights that proliferation signaling is limited by receptor occupancy.

##### 2. Dynamics of Antibody-Secreting Cells (ASCs)

ASCs (plasma cells) are generated via antigen-dependent differentiation of B cells and are responsible for soluble antibody production.

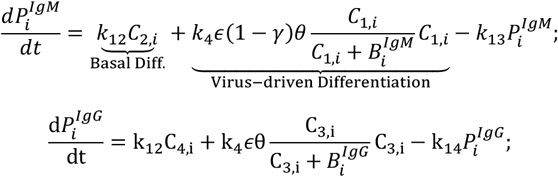

where P_i_ represents the ASC population. ϵ is a scaling factor accounting for the stoichiometric ratio between BCR density and cell number during differentiation. k_12_ denotes basal differentiation driven by environmental antigens, while k_13_ and k_14_ are the apoptotic rates for IgM- and IgG-ASCs, respectively.

##### 3. Soluble Antibody Dynamics

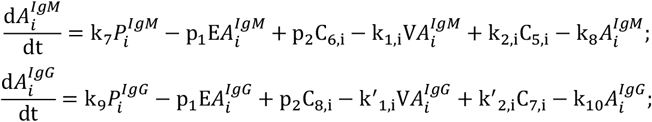

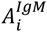and 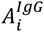denote soluble antibodies. k_7_ and k_9_ represent the secretion rates of IgM and IgG, respectively.

##### 4. Immune Complexes Kinetics

The model tracks eight distinct species of immune complexes (C_1,*i*_…C_8,*i*_), subject to a uniform clearance rate k_11_.

*Note: For brevity, representative equations for viral ( V) and environmental (E) complexes are shown below*.

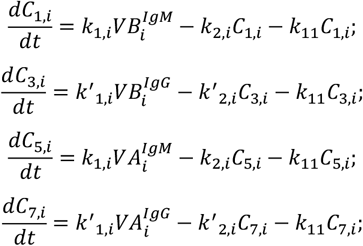

Complexes involving environmental antigens (C_2_,C_4_,C_6_,C_8_) follow an analogous structure determined by parameters p_1_ (association) and p_2_ (dissociation).

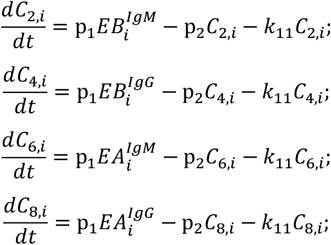

##### 5. Antigen and Global Dynamics

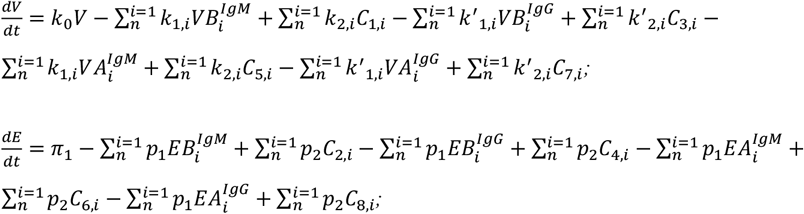

In the expression for viral load (V), the summation accounts for the removal of free virus by binding to all BCRs and soluble antibodies across all clones *i*. Similarly, environmental antigen (E) is maintained by a flux π_1_ and consumed by non-specific binding.

#### Parameter Interpretations and Structural Constraints

##### Avidity vs. Affinity

To accurately reflect biological structure, we distinguish between the kinetic parameters of IgM and IgG. While the variable region sequences (and thus intrinsic affinity) may initially be identical for a given clone *i*, IgM exists predominantly as a pentamer, whereas IgG is monomeric. This results in a significantly higher functional avidity for IgM. Consequently, we impose the following constraints in our simulations:

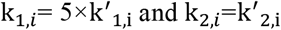

This condition (k_1,*i*_≫k^′^_1,*i*_) provides a mechanistic explanation for the rapid, early-phase proliferation of IgM during acute infection, even prior to affinity maturation.

##### Key Parameters Definition

- k_0_: Viral replication rate (or antigen decay rate for non-replicating vaccines).
- k_3_: Background proliferation coefficient of B cells driven by environmental antigen-BCR cross-linking.
- k_4_: Clonal expansion coefficient driven by specific virus-BCR interaction.
- k_11_: Global clearance/degradation rate of immune complexes.
- k_12_: Rate of B-cell differentiation into ASCs induced by environmental background.
- *γ*: Class-switch recombination rate (IgM to IgG).
- θ: Terminal differentiation rate (B cell to ASC).
- ϵ: Scaling coefficient relating surface receptor dynamics to cellular differentiation stoichiometry.
- p_1_, p_2_ : Association and dissociation rate constants for environmental antigen binding (assumed constant across the repertoire to represent non-specific “tick-over”).

### 3.2 Modeling Cellular Adaptive Immunity

#### 3.2.1 Mechanisms of CD8+ T Cell Priming

We categorized cellular immunity into two distinct pathways based on the mechanism of dendritic cell (DC) priming and antigen presentation. The governing dynamics are primarily driven by the activation and clonal expansion of CD8+ cytotoxic T lymphocytes ( Tc).

##### Type I Antigens (Pathogen-Associated Structural Recognition)

Antigens such as SARS-CoV-2 possess intrinsic pathogen-associated molecular patterns (PAMPs) recognized by Pattern Recognition Receptors (e.g., TLRs and CLRs) on DCs. Consequently, these antigens can induce DC maturation and subsequent MHC-I cross-presentation directly in their free form (V), or via Fc γ receptor-mediated uptake of antigen-antibody immune complexes (C).

##### Type II Antigens (Structure-Independent/Self-Antigens)

Antigens lacking specific PAMPs (e.g., self-antigens or certain strains like LCMV) require opsonization to facilitate uptake. In this context, DC priming is strictly dependent on the formation of immune complexes, enabling entry via Fc receptors.

To describe these dynamics mathematically, we formulated a unified system of ordinary differential equations (Model 3.2.1):

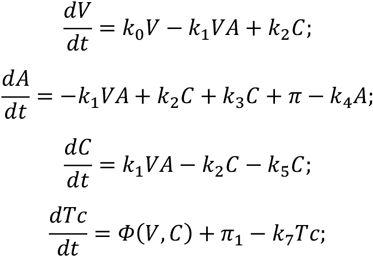

The activation function Φ(V, C) distinguishes the two antigen types:

- **For Type I Antigens:** *ϕ*(*V, C*)=*k*_6_(*C* + *V*), reflecting dual activation pathways.
- **For Type II Antigens:** *ϕ*(*V, C*)=*k*_6_*C*, reflecting strict dependence on immune complex formation.

#### Parameter Interpretation

The parameter *k*_0_ signifies the viral replication rate in natural infection scenarios ( *k*_0_>0). Conversely, in vaccination contexts involving non-replicating antigens, *k*_0_ assumes a negative value to represent in vivo antigen decay. *k*_1_ and*k*_2_ denote the association and dissociation rate constants of the antigen-antibody interaction, respectively. The coefficient *k*_3_ quantifies the positive feedback of immune complexes on antibody regeneration; notably, this parameter is significantly reduced in patients undergoing B-cell depletion therapy (e.g., Rituximab). *k*_5_ represents the clearance rate of immune complexes, which typically exceeds the natural decay of free antigen. Detailed parameterization is provided in the Supplementary Materials.

#### 3.2.2. Antigen Density-Dependent Cytotoxicity and Avidity Maturation

To elucidate the mechanistic link between intracellular antigen density and cytotoxic efficiency, we modeled the formation of the immunological synapse between Tc cells and target cells. Let N_i_ denote a sub-population of infected cells presenting a specific intracellular viral load V_i_. The surface antigen density is defined as the ratio V_i_/N_i_.

The interaction dynamics are governed by the reversible formation of the cytotoxic complex C_i_ (Model 3.2.2):

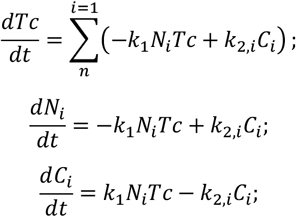

Here, k_1_ represents the forward association rate constant, assumed to be uniform across populations and determined by the collision frequency between T cells and targets. Crucially, the stability of the cytotoxic synapse—and thus the killing efficiency—is governed by the reverse dissociation rate constant 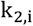, which is modulated by antigen density.

Based on the thermodynamic principle that higher antigen density increases the avidity of the TCR-MHC-I interaction (thereby reducing the off-rate), we derived a log-linear relationship for k_2,*i*_:

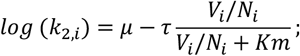

Where:

- μ represents the baseline dissociation constant (log scale) in the absence of antigen flux.
- τ quantifies the maximum gain in binding energy (avidity) achievable at saturation.
- *Km* is the Michaelis-Menten constant, representing the antigen concentration at which the specific biophysical effect reaches half-maximal intensity.

This formulation captures the non-linear enhancement of T-cell dwell time on target cells as intracellular viral load increases, providing a quantitative basis for high-affinity structural recognition.

### 3.3 Spatiotemporal Dynamics of B Cell-CD4+ T Cell Cognate Interactions

To mechanistically capture the T-dependent humoral response, we developed a multi-scale model (Model 3.3.1) that explicitly describes the “linked recognition” process. This model tracks the endosomal processing of BCR-antigen complexes, the surface presentation of peptide-MHC class II (pMHC-II) ligands, and the subsequent cognate interaction with helper T cells (Th).

The system describes the cross-regulation between *n* clones of B cells (indexed by *i*) and *m* clones of CD4+ T cells (indexed by *j*). We distinguish between naïve/effector T cells (T^(1)^) and memory T cells (T^(2)^).

#### 1. B-Cell Clonal Dynamics and Isotype Switching

The expansion of IgM 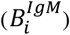 and IgG 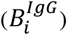 B-cell clones is driven by signals received from immunological synapses formed with CD4+ T cells.

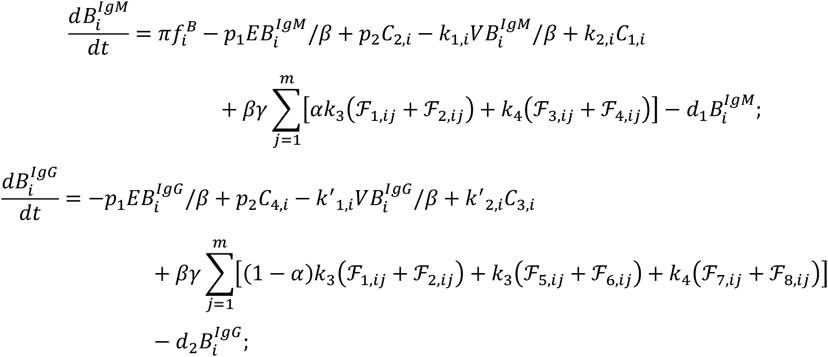

Here, ℱ_*k*,i*j*_ represents the synapse complex formed between B cell clone *i* and T cell clone *j*. The term *β* scales the BCR density per cell, and *γ* quantifies the stoichiometric amplification of B-cell proliferation induced by T-cell help. *α* denotes the probability of IgM-to-IgG class switching upon T-cell priming.

##### 2. CD4+ T Cell Recruitment and Memory

T cells are recruited into synapses by pMHC-II complexes (D) presented on B cells.

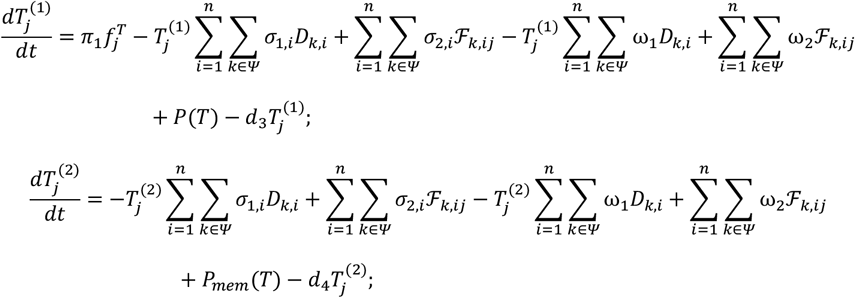

where P(T) represents the T-cell proliferation feedback driven by successful synapse formation.

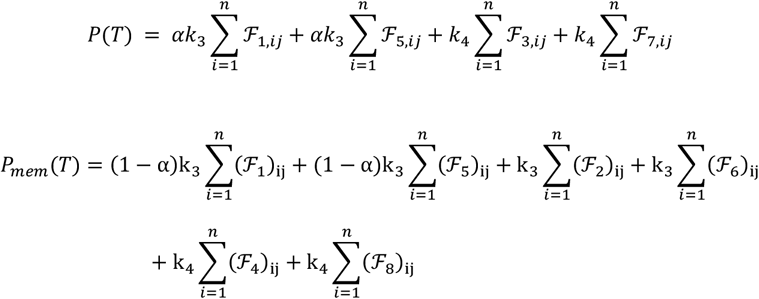

#### 3. Antigen Processing and Presentation Pipeline

The model tracks the lifecycle of the antigen from surface binding ( C*C*) to endosomal degradation and surface presentation as pMHC-II (D*D*).

**Step 1: Surface Binding (Representative for Virus-IgM):**

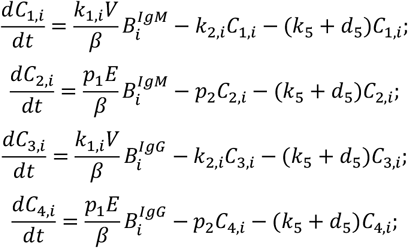

Here, *k*_5_ represents the rate of internalization and processing into pMHC-II, while *d*_5_ denotes non-productive degradation.

**Step 2: pMHC-II Presentation:**

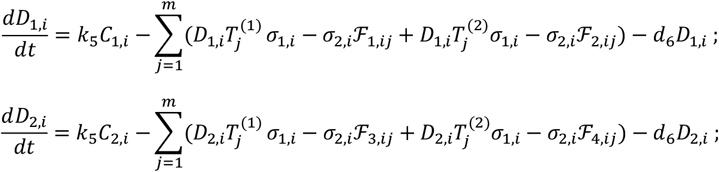

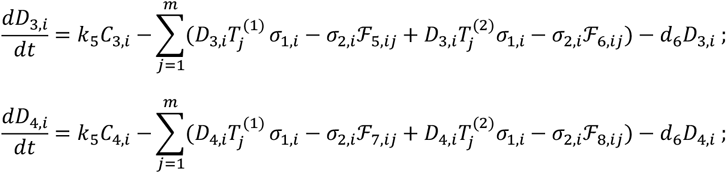

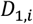 represents the viral peptide-MHC complex derived from IgM-mediated uptake. *D*_2,*i*_, *D*_3,*i*_, *D*_4,*i*_ correspond to complexes derived from environmental antigen (IgM), viral antigen (IgG), and environmental antigen (IgG), respectively.

**Step 3: B-T Synapse Formation (FF):**

The formation of the cognate synapse ℱ_*k*,i*j*_ is governed by:

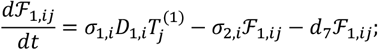

This generic form applies to all 8 permutations of interaction (Naive/Memory T cells × 4 types of Antigen sources).

#### 4. Global Antigen Dynamics

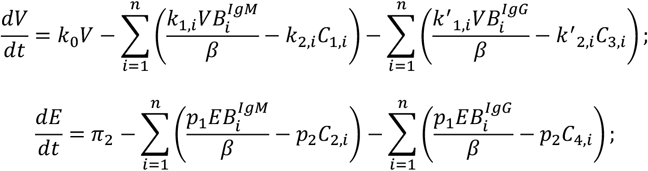

##### Parameter Definitions

- *σ*_1,*i*_, *σ*_2,*i*_: Association and dissociation rates for the TCR–pMHC-II interaction.
- ω_1_, ω_2_: Kinetic constants for interactions involving environmental antigens (assumed to have lower/baseline affinity).
- 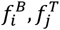: Initial precursor frequencies for B and T cells, distributed normally based on affinity to reflecting the naïve repertoire.
- *d*_6_, *d*_7_: Turnover rates for pMHC-II complexes and immunological synapses, respectively.

### 3.4 Cohort-Based Model of Temporal Viral Infection Dynamics

To capture the heterogeneity of viral replication and immune clearance based on the duration of infection, we utilized a cohort-structured agent-based framework. Unlike standard ODE models that treat all infected cells identically, this model tracks cohorts of cells *I*(*τ,t*) infected at a specific initiation time *τ*, evolving over current time *t*.

#### 3.4.1 Susceptible Cell Dynamics and de novo Infection

The population of susceptible cells, *T*(*t*), evolves according to a replenishment rate *k*_6_, a natural decay rate *k*_7_, and a viral infection term modeled by Hill kinetics. At each time step Δ*t*, the number of newly infected cells (initiating cohort *τ*=*t*) is calculated as:

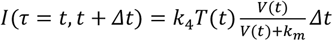

where *k*_4_ is the viral invasion constant, *V*(*t*) is the extracellular viral load, and *k*_*m*_ is the half-maximal infection constant. Consequently, the susceptible population is updated:

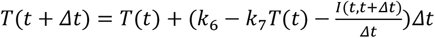

#### 3.4.2 Structured Population Dynamics and Numerical Implementation

To transcend the limitations of numerical stiffness and discretization artifacts inherent in discrete cohort-based or agent-based models (ABMs)—particularly when coupling continuous intracellular viral replication with binary lysis thresholds—we developed a continuous structured population model based on the McKendrick-von Foerster equation. By transforming the discrete “time since infection” (*τ*) into a continuous variable, infection age (*a*), we describe the infected cell population density *i*(*a,t*), where *i*(*a,t*)*da* represents the number of cells at time *t* that have been infected for a duration between *a* and *a*+*da*.

##### Governing Equations

The dynamics of the infected cell population are governed by the following transport equation with a variable sink term:

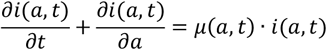

The boundary condition at *a*=0 represents the rate of de novo infection, coupled with the susceptible cell population *T*(*t*):

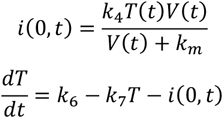

##### Continuous Hazard Functions for Cell Death

A critical modification in our approach is the replacement of binary threshold logic (e.g., if *V*_*in*_ >*θ*,then lysis) with continuous hazard functions. This ensures numerical stability and biological realism. The total instantaneous mortality rate *μ*(*a,t*) is defined as the sum of three distinct mechanisms:

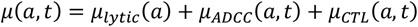

1. **Natural Viral Lysis (***μ*_*lyt*i*c*_): Modeled using a Hill function dependent on intracellular viral load

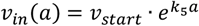

This mimics the probabilistic nature of membrane rupture as viral pressure increases:

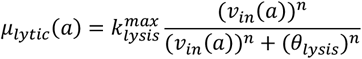

where *n* is the Hill coefficient controlling the steepness of the transition, and *θ*_*lys*i*s*_ is the viral load threshold.
2. **ADCC-Mediated Death (** *μ*_*ADCC*_): Dependent on the formation of antigen-antibody complexes Φ(*a,t*) on the cell surface:

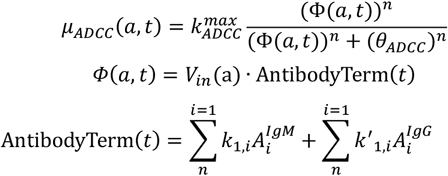
3. **CTL-Mediated Killing (***μ*_*CTL*_): Modeled via mass-action kinetics proportional to the cytotoxic T-lymphocyte count *Tc*(*t*):

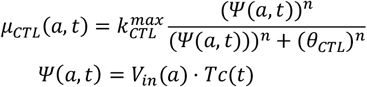

##### Viral Release Dynamics (Integral Formulation)

To calculate the production of extracellular virus *V*(*t*), we integrate the contribution of all dying cells over the infection age domain. The dynamics of free virus are described by:

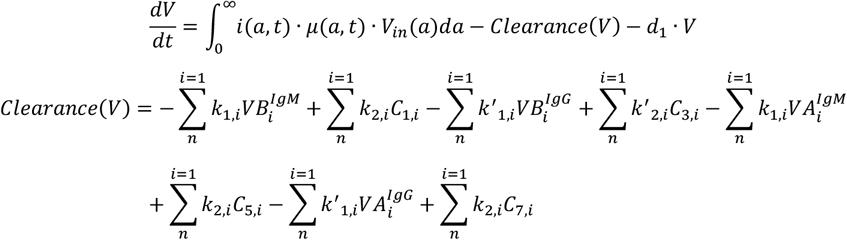

##### Modeling the proliferation and exhaustion of Tc cells

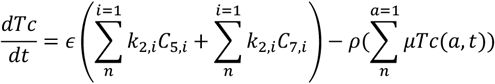

Where *C*_7,*i*_ represents the complexes formed by IgG and the virus, and *C*_5,*i*_ represents the complexes formed by IgM and the virus. *ϵ* is the activation constant for CD8+T cells, *ρ* is the exhaustion constant of CD8+T cells.

The dynamics of other immunological components, such as antibodies, BCR-B cells, plasma cells, and various antibody-antigen complexes, are governed by the description in Model 3.1.7.

##### Numerical Solution: Method of Lines

The partial differential equation (PDE) system was solved using the Method of Lines (MOL). The continuous age domain was discretized into *N*=400 bins with step size Δ*a*=1 hour. The spatial derivative 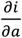 was approximated using a first-order upwind difference scheme to ensure transport stability.

- The infection age a*a* is discretized into the grid points *a*_0_, *a*_1_,…, *a*_*N*_ with a step size of Δ*a* (here, we set Δa=1 and *N*=400).
- Let *I*_*j*_ (*t*) ≈ i(*a*_*j*_, *t*).
- The partial derivative 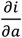 in the PDE is approximated by the difference quotient 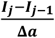 (Upwind scheme).
- Consequently, the original PDE is transformed into a large-scale system of ODEs:

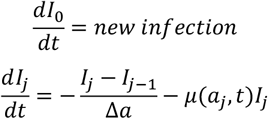

The resulting system of Ordinary Differential Equations (ODEs) was integrated in MATLAB (MathWorks, Inc.) using the ode15s solver to handle the stiffness arising from the rapid activation of Hill functions.

### 3.5 Tumor-Host Interaction Model

#### 3.5.1 System Dynamics

We formulated a deterministic system of ordinary differential equations (ODEs) to quantify the dynamic interplay between tumor progression, neoantigen release, and the dual pressures of humoral (antibody-mediated) and cellular (CD8+ T cell-mediated) immunity. The system tracks five state variables: tumor load (*Tu*), free neoantigen concentration (*N*), antibody-neoantigen complexes (*C*), specific antibody titer (*A*), and cytotoxic CD8++ T cells (*Tc*).

The conceptual framework posits that tumor cells release neoantigens upon death (natural or immune-mediated). These antigens form complexes with antibodies, which subsequently stimulate the recruitment or proliferation of cytotoxic T cells.

The governing equations are defined as follows:

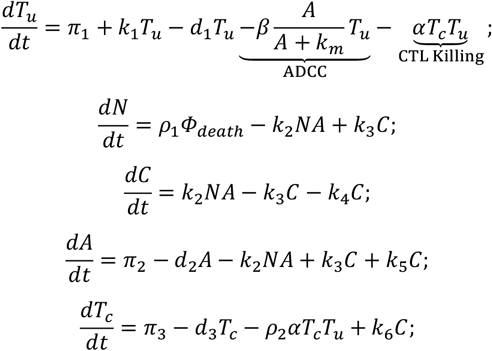

#### 3.5.2 Flux Definitions and Mechanism Description

##### Tumor Growth and Lysis

Tumor dynamics are governed by a constant basal influx *π*_1_ (e.g., stem cell differentiation), an exponential proliferation rate *k*_1_, and a natural death rate *d*_1_. Immune clearance occurs via two pathways:

1. ADCC (Antibody-Dependent Cellular Cytotoxicity): Modeled by a saturation function (Michaelis-Menten kinetics), where *β* is the maximum lysis rate and *k*_*m*_ is the antibody concentration at half-maximal efficacy.
2. CTL Killing: Represented by a mass-action term *αT*_*c*_*T*_*u*_, where *α* is the cytotoxic efficiency of CD8+ T cells.

##### Neoantigen Release (*ϕ*_*death*_)

The total rate of tumor cell death, which drives neoantigen release, is defined as 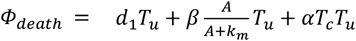. Free neoantigens (*N*) accumulate proportional to cell death (scaled by *ρ*1, the neoantigen load per cell) and are consumed by binding to antibodies. Immune Complex and Effector Dynamics

- Complex Formation (*C*): Reversible binding follows mass-action kinetics with association constant *k*_2_ and dissociation constant *k*_3_. Complexes degrade at rate *k*_4_.
- Antibody Dynamics (*A*): Antibodies are replenished at rate *π*_2_ and decay naturally at rate *d*_2_. The term *k*_5_*C* represents the recycling or regeneration of antibodies following complex processing.
- T Cell Dynamics (*Tc*): CD8+ T cells are maintained by homeostatic influx *π*_3_ and stimulated by the presence of antigen-antibody complexes at rate *k*_6_ (representing cross-presentation mechanics). The term −*ρ*_2_*αTcTu* accounts for T cell exhaustion or loss during the effector phase, where *ρ*_2_ quantifies the exhaustion rate per killing event.

**Table 1:**
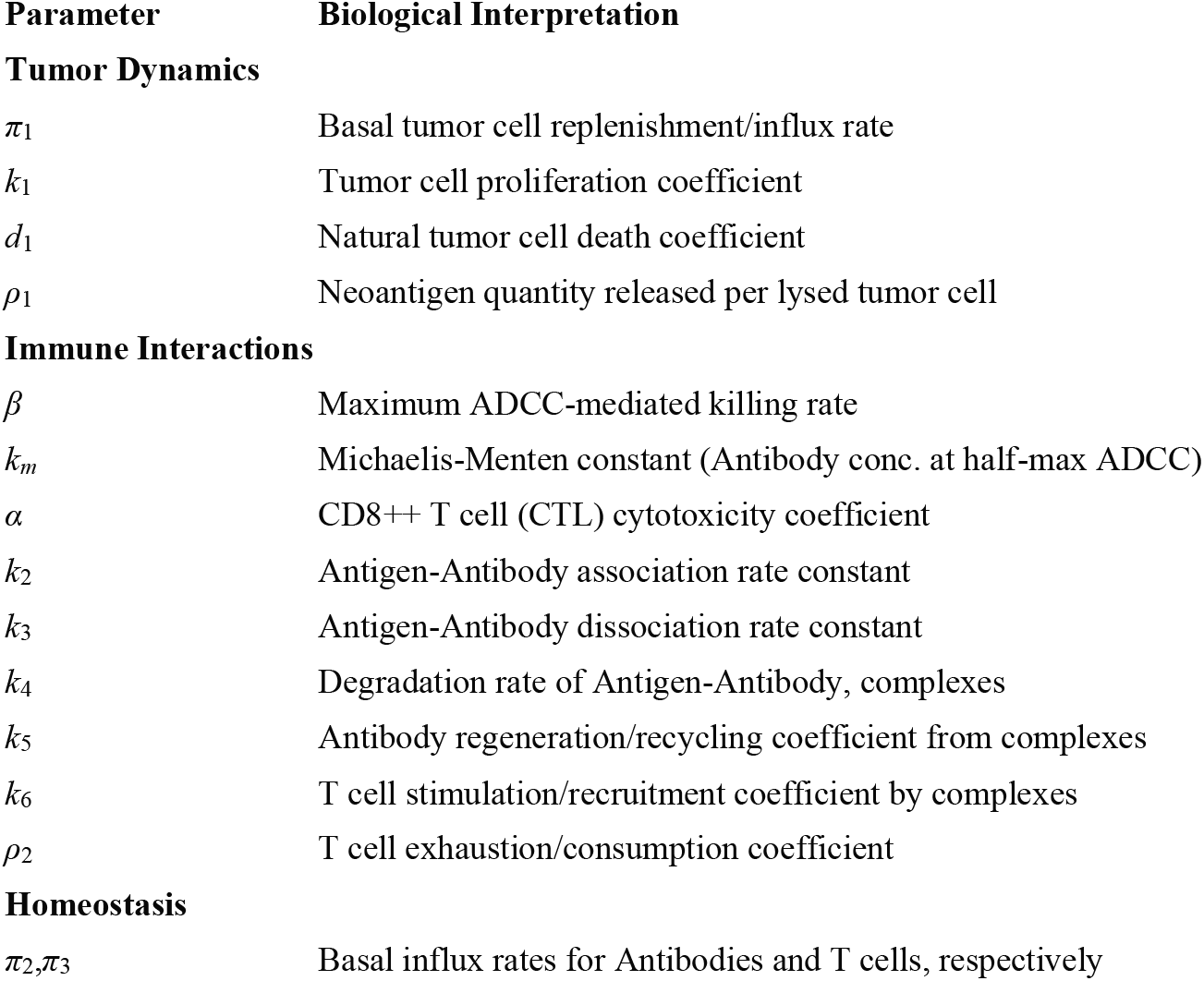
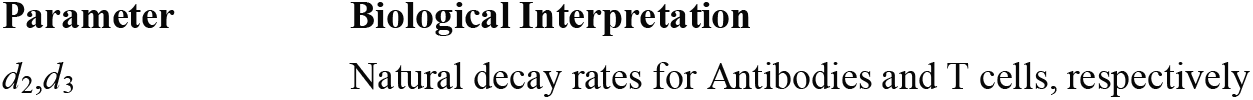
Model Parameters.

## Discussion

Mathematical modeling has long served as a cornerstone for quantifying virus-host dynamics; however, the predictive power of classical models is increasingly constrained by structural oversimplifications. In this study, we introduce a high-fidelity, agent-based framework that bridges the gap between reductionist kinetics and systems immunology. By prioritizing mechanistic accuracy over phenomenological fitting, our model resolves three fundamental limitations inherent in canonical *SIR*-type approaches: distorted infection kinetics, the segregation of immune arms, and the neglect of clonal diversity.

### Reevaluating Infection Kinetics and Immune Activation

A critical departure in our work is the rejection of the standard mass-action bilinear term (*βTV*) for viral infection. While mathematically convenient, this formulation presupposes second-order kinetics between equivalent entities—an assumption that collapses under the vast stoichiometric disparity between virions and cells. As detailed in our Supplementary Materials, biological infection is a saturation process best described by Michaelis-Menten-like kinetics rather than infinite linear approximations. Standard models relying on *βTV* often erroneously predict that infection resolution is driven by the depletion of susceptible cells (*T*). In contrast, our saturation-based approach preserves the susceptible population, demonstrating that viral clearance is predominantly an immune-mediated event.

Furthermore, we reconstructed the governing logic of adaptive immunity. Classical equations often model Cytotoxic T Lymphocyte (CTL) expansion as a direct function of infected cell density (*αIE*). Biologically, however, direct interaction with non-professional antigen-presenting cells often leads to CTL anergy or exhaustion rather than proliferation. Our model corrects this by linking CTL expansion to antigen-antibody complexes and cross-presentation dynamics. This distinction is not merely semantic; it provides a mechanistic basis for the emergence of T cell exhaustion during chronic infections—a phenomenon that phenomenological models fail to capture endogenously.

### Integration of Humoral and Cellular Synergy

Previous modeling efforts have largely treated partial immune responses in isolation. By integrating humoral and cellular arms into a unified system, we reveal emergent properties of immune synergy. We demonstrate that antibodies do not function solely as neutralizing agents but are critical drivers of cellular immunity via complex formation and ADCC (Antibody-Dependent Cellular Cytotoxicity). This coupled framework allows for the investigation of complex clinical scenarios, such as the interplay between neutralizing antibodies and CD8+ T cell priming, offering a theoretical platform to resolve debates regarding the mechanisms of “original antigenic sin” and the specific kinetic profiles of IgM versus IgG in secondary infections.

### The Role of Receptor Diversity in System Dynamics

Perhaps the most significant advance of this framework is the explicit incorporation of BCR and TCR clonal diversity. In the era of single-cell sequencing and high-dimensional flow cytometry, models that rely on bulk “average” lymphocyte populations are becoming obsolete. Our approach simulates the evolutionary pressure on receptor affinity, allowing us to map the trajectory of affinity maturation and the stochastic nature of clonal expansion.

Applying this diversity-aware framework provided several key clinical insights:

1. **Chronic Infection Thresholds:** We identified that the transition from acute to chronic infection is often determined by a mismatch between viral replication rates and the temporal window of specific antibody induction. Notably, high initial titers of low-affinity antibodies (e.g., from heterologous challenges) can paradoxically facilitate chronicity by masking antigens from high-affinity clone selection.
2. **Therapeutic Vaccination:** Our simulations suggest that therapeutic vaccines for established chronic infections require significantly higher antigenic loads than prophylactic interaction to overcome established exhaustion milestones and regulatory feedback loops.
3. **Tumor-Immune Dynamics:** In the context of cancer, the model supports the efficacy of dual-target strategies (MHC-I and MHC-III antigens), predicting that successful suppression of tumor growth requires the synergistic engagement of neoantigen-specific antibodies and CTL infiltration.

### Limitations and Future Directions

While this framework represents a shift toward “quantitative systems immunology,” it is not without limitations. The complexity required to capture biological realism increases the parameter space, posing challenges for identifiability. Currently, our parameterization serves as a theoretical proof-of-concept. The next phase of validation requires not merely scalar clinical data (viral load, ELISA titers) but high-dimensional datasets, such as time-resolved BCR/TCR repertoire sequencing and epitope landscapes.

In conclusion, by moving beyond the “cell depletion” paradigm and explicitly modeling the diverse, synergistic architecture of the immune system, this work provides a rational, quantitative foundation for designing next-generation immunotherapies and understanding the delicate balance between viral clearance and immune pathology.

## Data Availability

The MATLAB codes can be accessed through the following link: https://github.com/zhaobinxu23/adaptive_immunity_new.

## Author Contributions

Conceptualization, S.W. and Z.X.; methodology, Z.X. and D.W.; software, validation, and formal analysis, C.Liu, W.Z., Q.Z., J.S., H.Z., L.L., C.Li and L.W.; investigation, Z.X., C.Liu and W.Z.; visualization, G.Y. and Z.X.; resources, B.H. and D.W.; data curation, C.Liu, W.Z. and Q.Z.; writing— original draft preparation, Z.X.; writing—review and editing, B.H. and D.W.; supervision, D.W.; funding acquisition, Z.X. All authors have read and agreed to the published version of the manuscript.

## Funding

This research was funded by DeZhou University, grant number 30101418.

## Institutional Review Board Statement

Not applicable.

---

**Extended Data Figure 1.**
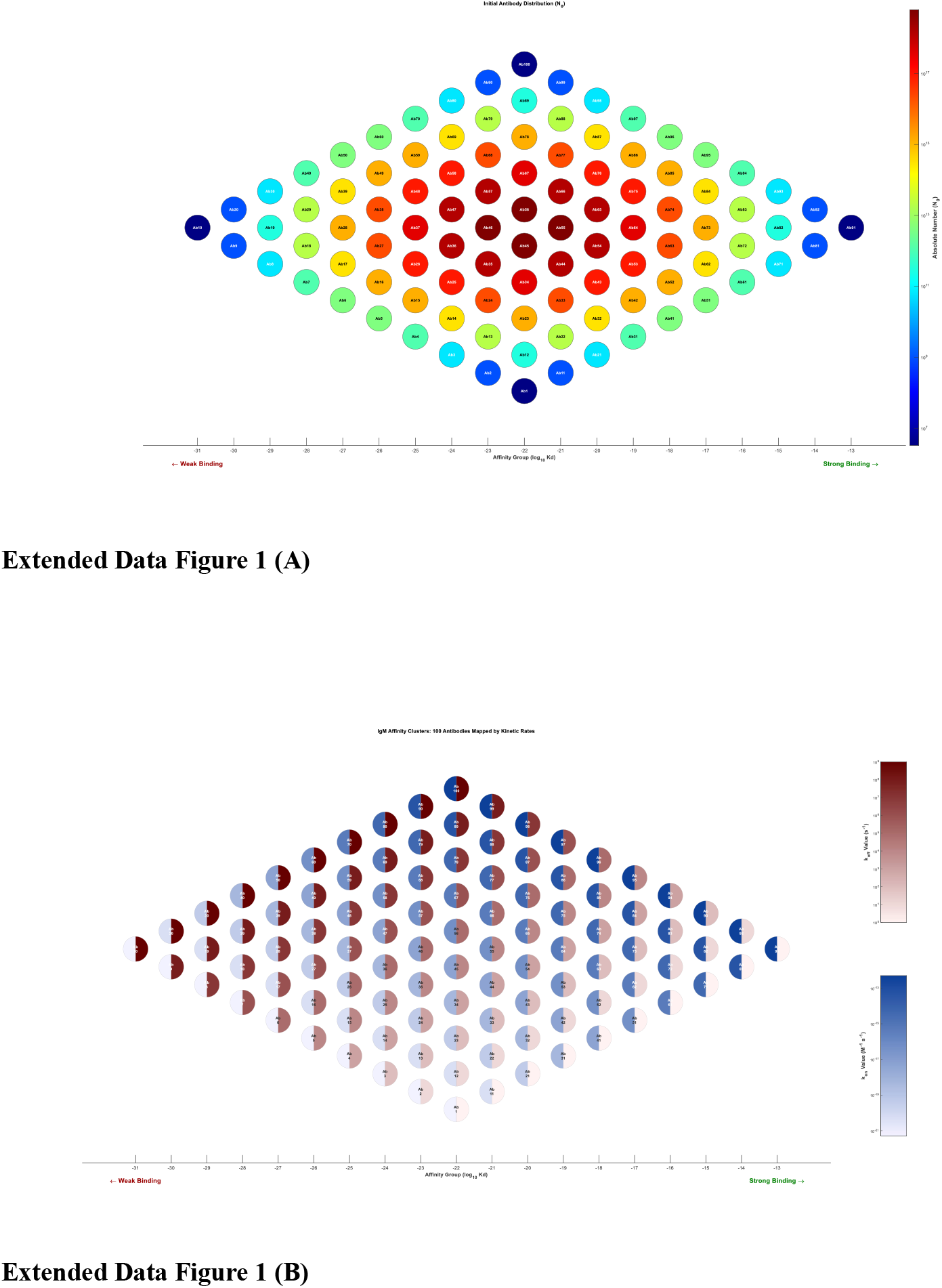
Initialization and kinetic parameterization of the polyclonal antibody repertoire. **(A)** Heatmap illustrating the initial concentration landscape of the modeled IgG-BCR repertoire, comprising 100 distinct clones. To capture immune heterogeneity, the repertoire is structured as a two-dimensional matrix where each clone *i* is uniquely indexed by a pair of integers (*m,n*) representing binding kinetics. The color intensity reflects the initial abundance of each clone, established based on a probability mass function. **(B)** Distribution profiles of the kinetic rate constants governing clonal diversity. The indices *m* and *n* (1≤m,n≤10) determine the forward association (*k*_*on*_) and reverse dissociation (*k*_*off*_) coefficients, respectively. The probability distribution for these kinetic coordinates is derived from a normal distribution (*N* (*μ*=5, *σ*=0.82)) for both variables X (association) and Y (dissociation). This setup ensures the initial repertoire mimics physiological affinity landscapes, where clone abundance follows a log-normal distribution prior to antigen-driven selection.

**Extended Data Figure 2.**
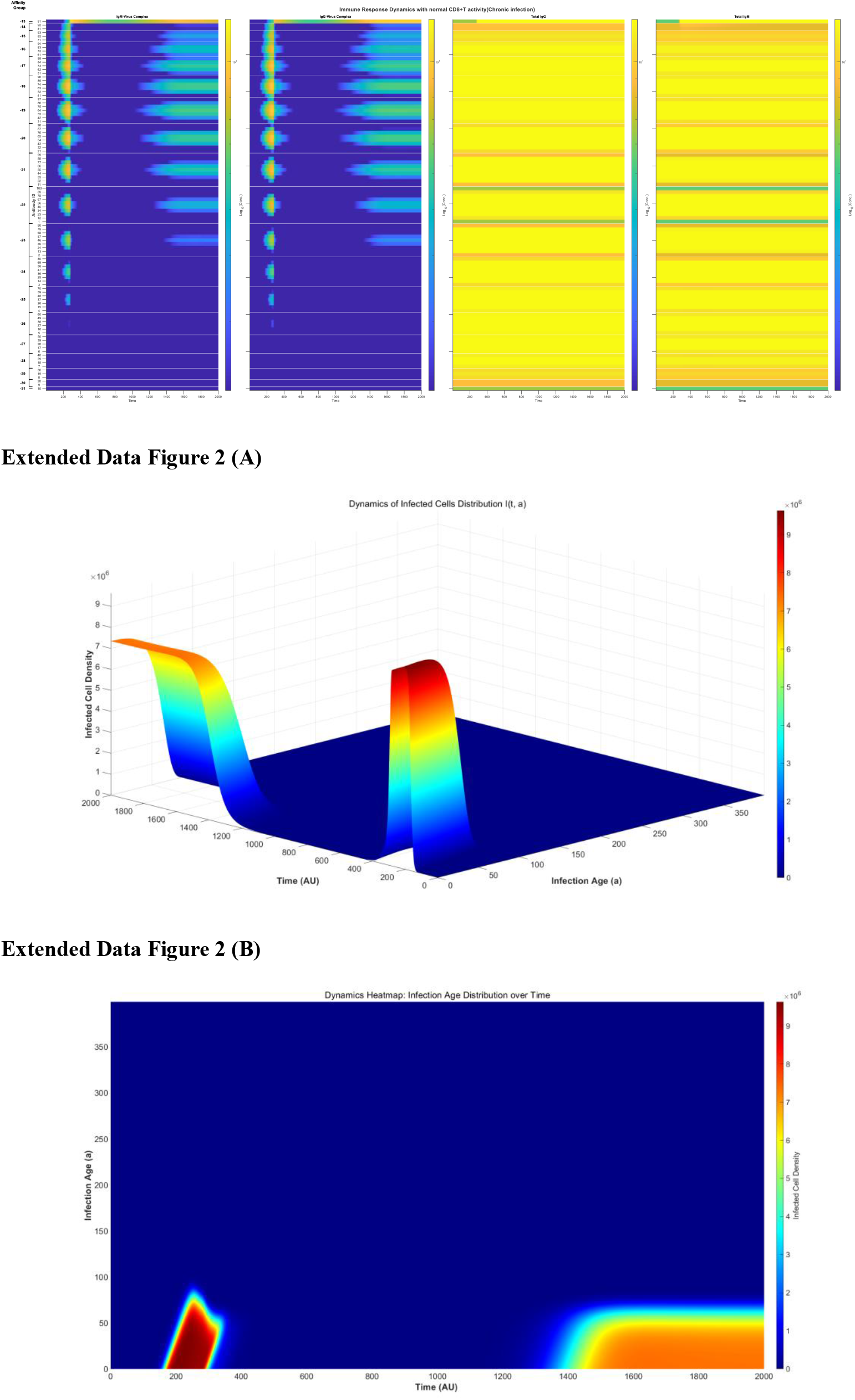
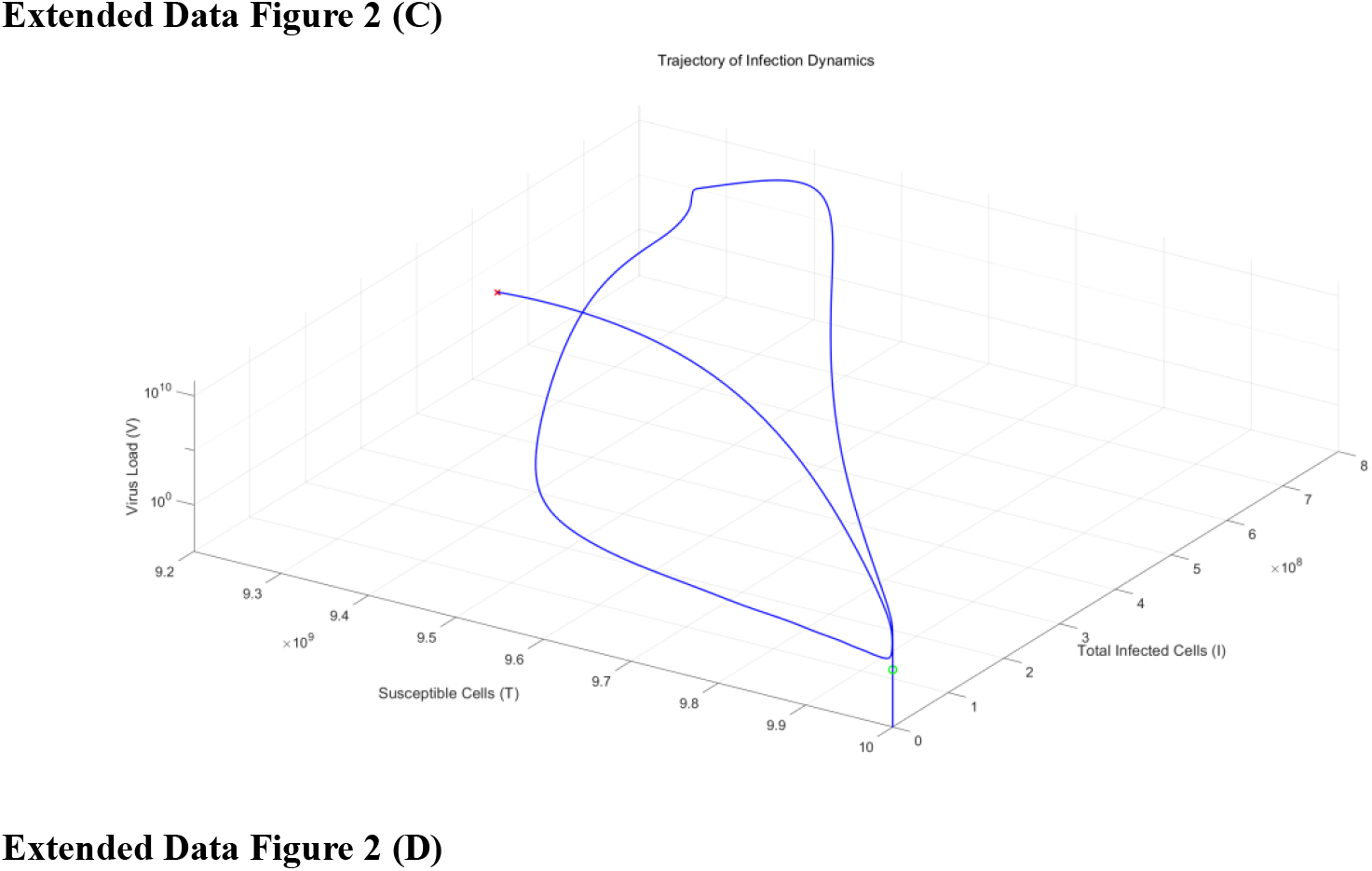
Spatiotemporal dynamics and equilibrium analysis of chronic viral infection. **(A)** Temporal kinetics of immune complexes and free antibodies under conditions of limited antibody regeneration capacity. Panels display (from left to right) the concentration of IgM-Virus complexes, IgG-Virus complexes, free IgM, and free IgG. Following the acute peak, viral complexes diminish during the latent phase (*t*→0) but subsequently rebound to a non-zero steady state, characterizing chronic infection. Notably, while high-affinity specific IgG levels remain elevated during the chronic phase—potentially hindering superinfection—they are insufficient to disrupt the established equilibrium, indicating a failure to achieve sterilizing immunity. **(B)** Three-dimensional evolution of the infected cell density *I*(*t,a*) as a function of time (*t*) and infection age (a*a*, ranging from 1 to 400). The distribution indicates that the infection age remains limited, as cells with high intracellular viral loads are rapidly cleared by ADCC and CTL-mediated cytotoxicity. The topology illustrates the progression from acute burst to latency, culminating in a persistent chronic state. **(C)** Two-dimensional heatmap projection of **(B)**, visualizing the density distribution of infected cells over time and infection age. The color gradient highlights the population shifts, clarifying the transition dynamics from acute instability to a stable, persistent infection pattern. **d**, Three-dimensional phase portrait of the infection dynamics trajectory. The system initiates at the onset of infection (green circle: *V*=10, Susceptible cells ≈10^10^, Infected cells =0) and converges to a stable endemic equilibrium (red cross). At this fixed point, susceptible cells stabilize at ≈9.2×10^9^, while both free virus and total infected cell populations maintain high non-zero levels, mathematically confirming the system’s entrapment in a chronic stable state rather than achieving viral clearance (*V*→0).

**Extended Data Figure 3.**
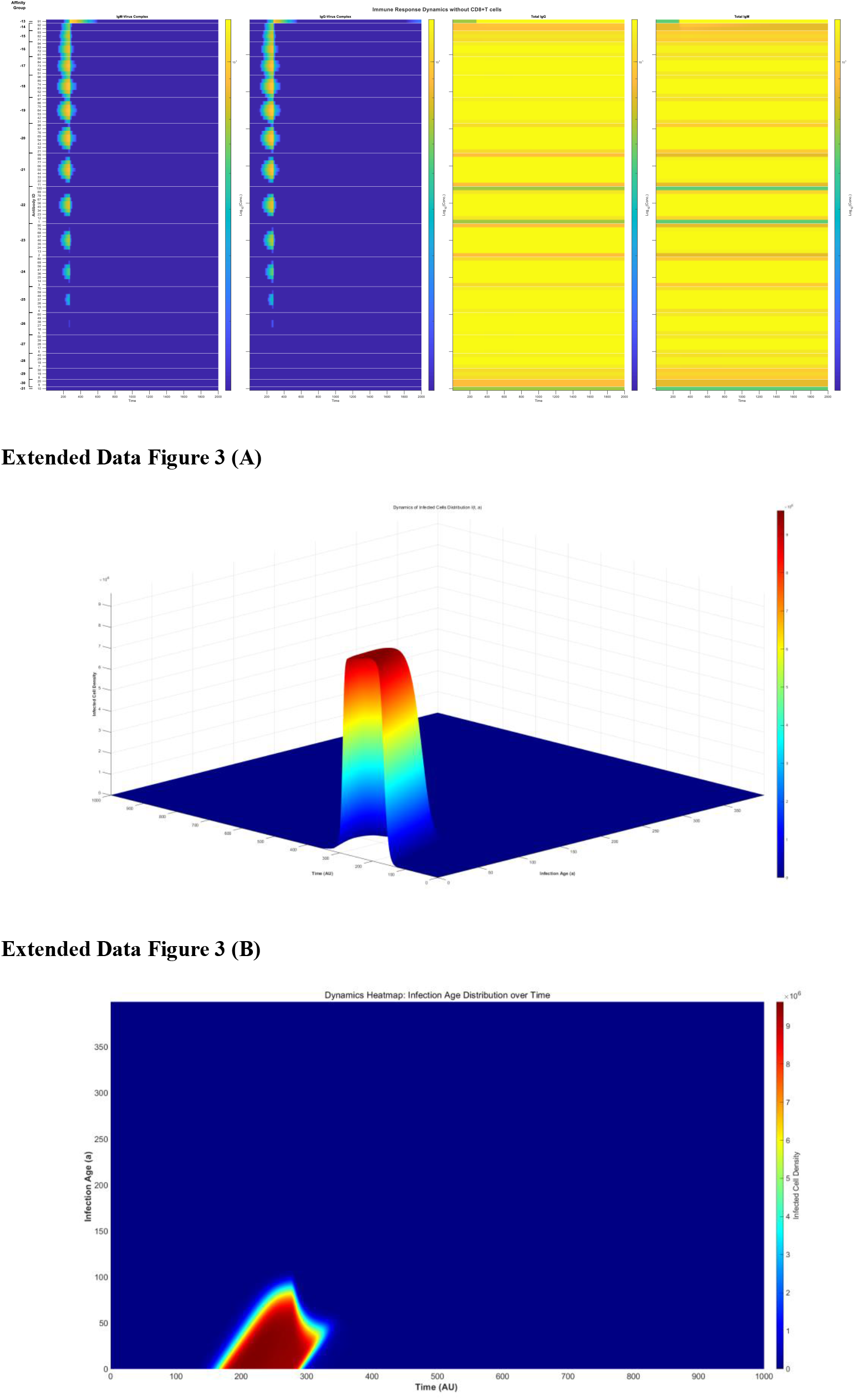
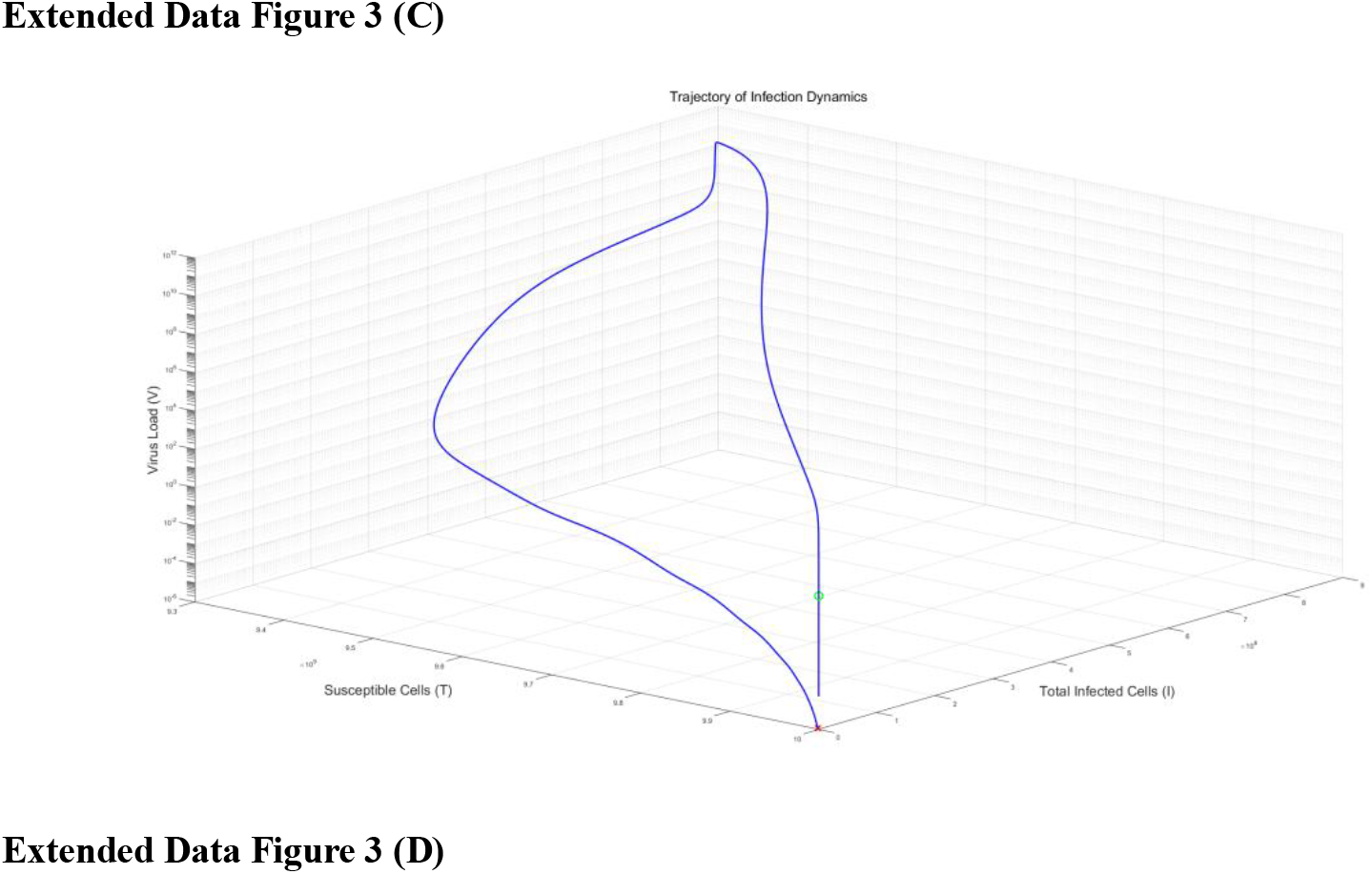
Humoral-mediated viral clearance in the absence of cellular immunity. **(A)** Temporal kinetics of immune complexes and free antibodies in a CD8+ T cell-deficient environment (*Tc*=0). Panels display (from left to right) the concentration of IgM-Virus complexes, IgG-Virus complexes, free IgM, and free IgG. In the absence of early viral containment by cellular immunity, the system experiences a higher antigen burden, which drives a more potent humoral response compared to the T cell-competent scenario (Extended Data Fig. 2A). Consequently, the peak concentrations of immune complexes and free antibodies are significantly elevated, providing sufficient neutralizing capacity to clear the infection and prevent the establishment of a chronic steady state. **(B)** Three-dimensional evolution of infected cell density *I*(*t,a*) as a function of time (*t*) and infection age (*a*). The landscape illustrates a robust acute infection trajectory that is rapidly terminated. Unlike the chronic scenario, the infected cell population is comprehensively eliminated during the acute phase, leaving no residual reservoir to seed persistent infection. **(C)** Two-dimensional heatmap projection of **(B)**, visualizing the population dynamics of infected cells. The intensity gradient highlights the transient nature of the infection, confirming that the elevated antibody response successfully blocks the transition from acute latency to chronicity. **(D)** Three-dimensional phase portrait of the infection dynamics trajectory. The system initiates at the point of infection (green circle: *V*=10, Susceptible cells =10^10^, Infected cells =0) and converges to a stable disease-free equilibrium (red cross). At this terminal state, both total infected cells and free virions return to zero (*V*→0, I→0), while the susceptible cell population recovers to its initial capacity (S≈10^10^), mathematically demonstrating complete viral clearance driven by the compensatory humoral response.

**Extended Data Figure 4.**
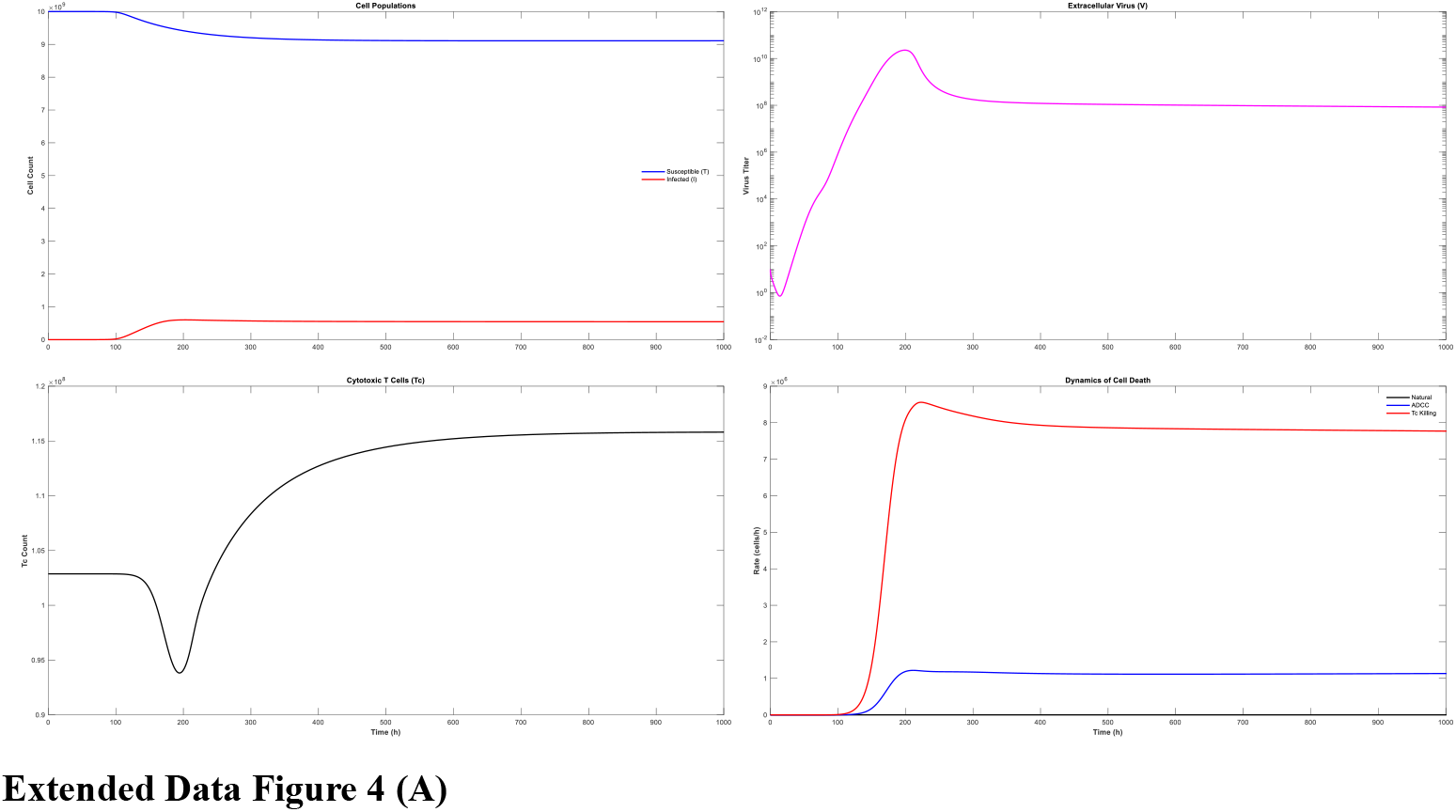
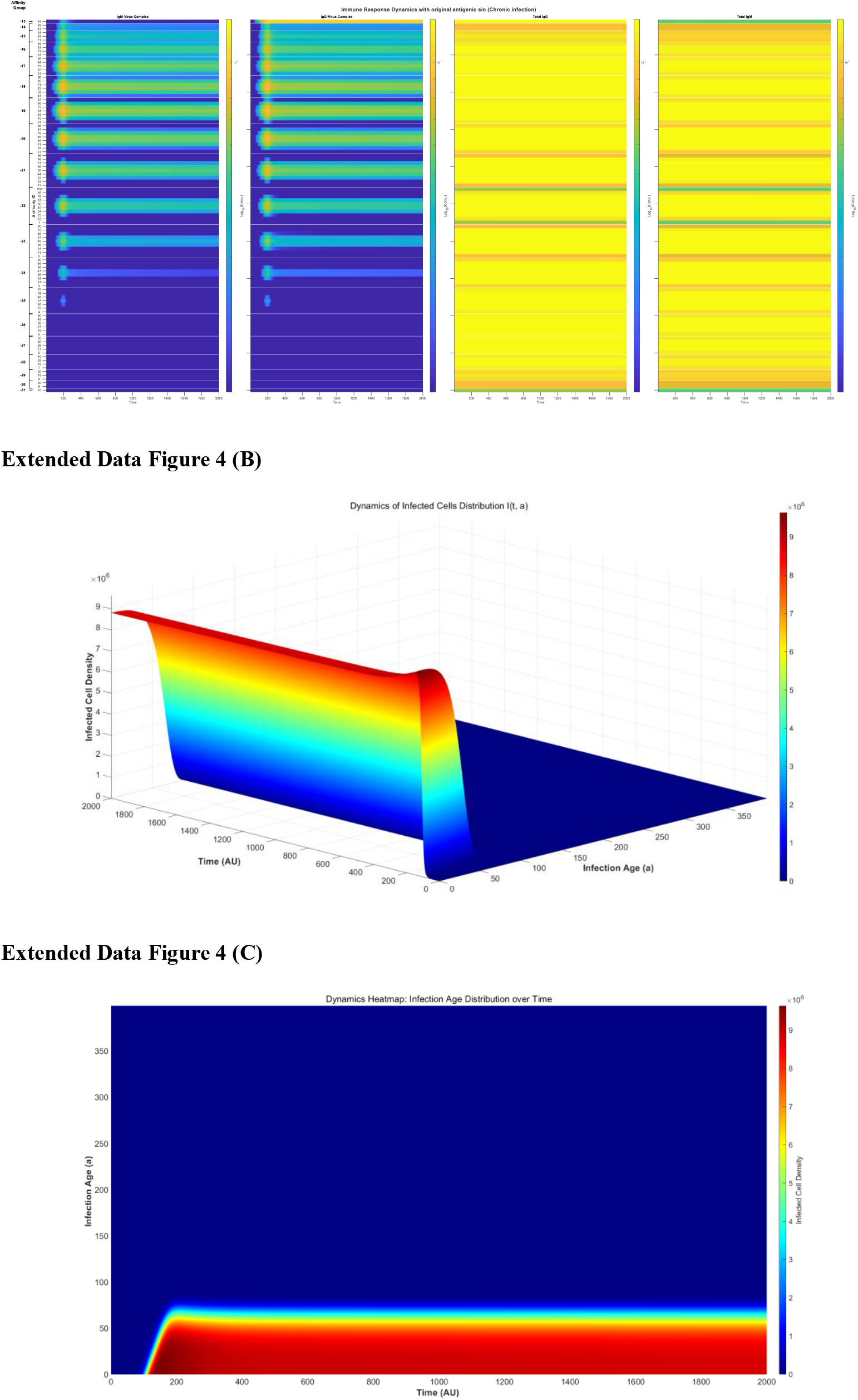
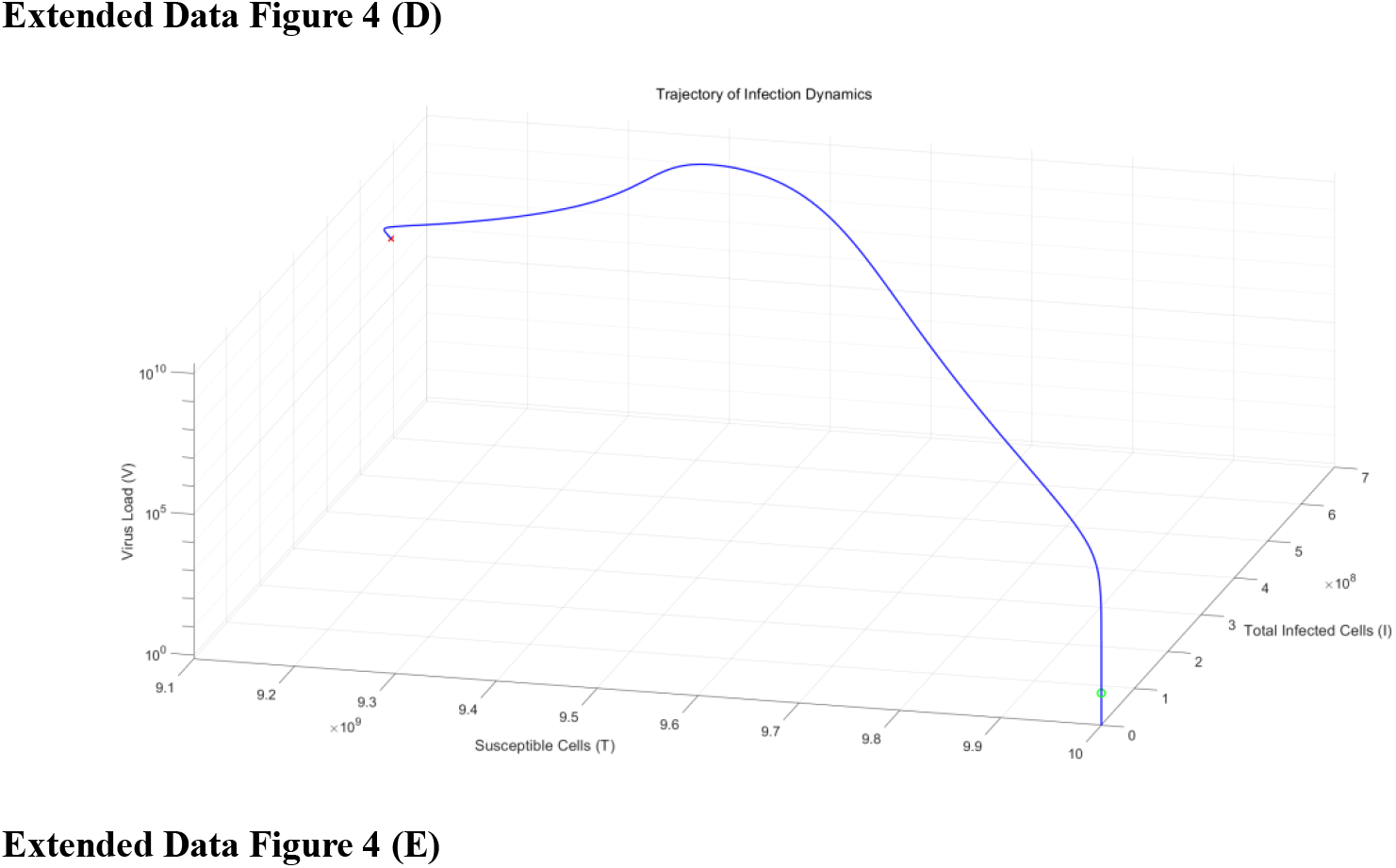
Dynamics of immune imprinting-induced chronic infection during reinfection with a viral variant. **(A)** Temporal dynamics of distinct host and viral compartments during reinfection with a variant strain (variation coefficient = 4) under the influence of immune imprinting. Panels show (from left to right): the evolution of susceptible and infected cells, exhibiting a shift towards a stable infected steady state; free virus (*V*) kinetics, which rebound to maintain a persistent low-level equilibrium (noting that peak viral load is suppressed compared to primary infection due to imprinting); proliferation of specific CD8 + T cells; and decomposition of cell lysis rates mediated by ADCC, Tc cytotoxicity, and viral cytopathic effects, confirming persistent cell death associated with chronic infection. **(B)** Kinetics of humoral immune responses and viral complex formation. Plots display the temporal levels of IgM-Virus complexes, IgG-Virus complexes, free IgM, and free IgG (from left to right). Due to immune imprinting, high-avidity IgG exhibits a high baseline but limited expansion, leading to the stabilization of immune complexes at a steady state rather than clearance. **(C, D)** Spatiotemporal analysis of infected cell populations. **C**, Three-dimensional representation of infected cell density *I*(*t,a*) as a function of time (*t*) and infection age (*a*, range 1–400). **D**, Corresponding two-dimensional heatmap projection. These panels illustrate that while cells with high infection age are cleared (by ADCC/Tc effects), a population of infected cells persists at a stable concentration, establishing a chronic infection reservoir. **(E)** Three-dimensional phase portrait of infection dynamics illustrating the relationship between free virus (*V*), susceptible cells (*T*), and total infected cells. The green circle marks the initial naïve state (*V*=0, *T*=10^10^, Total Infected=0). The red cross (×) denotes the final stable chronic equilibrium, where susceptible cells stabilize at 9.1×10^9^ while viral load and infected cells maintain a non-zero high plateau, indicating a failure of complete viral clearance.

**Extended Data Figure 5.**
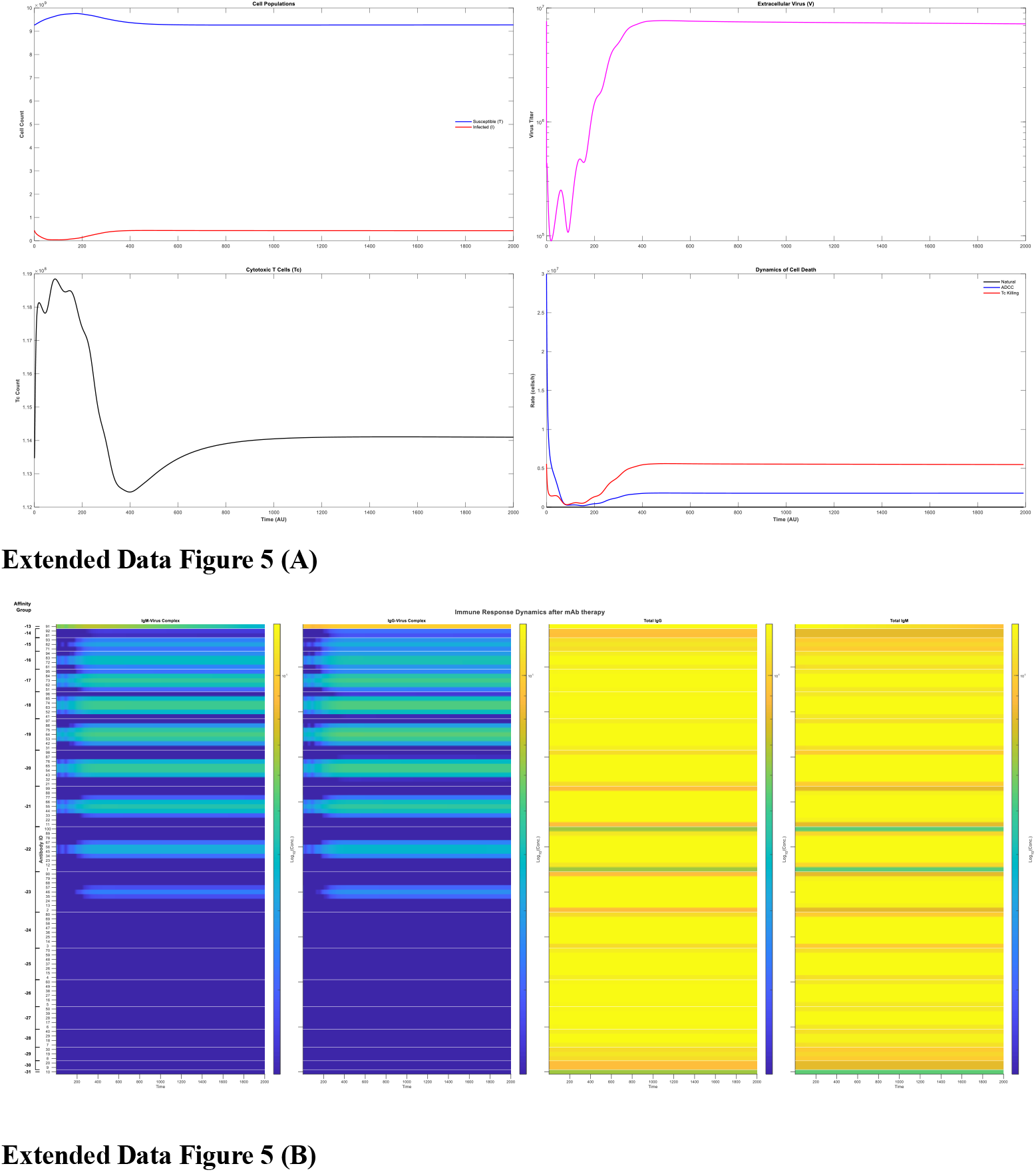
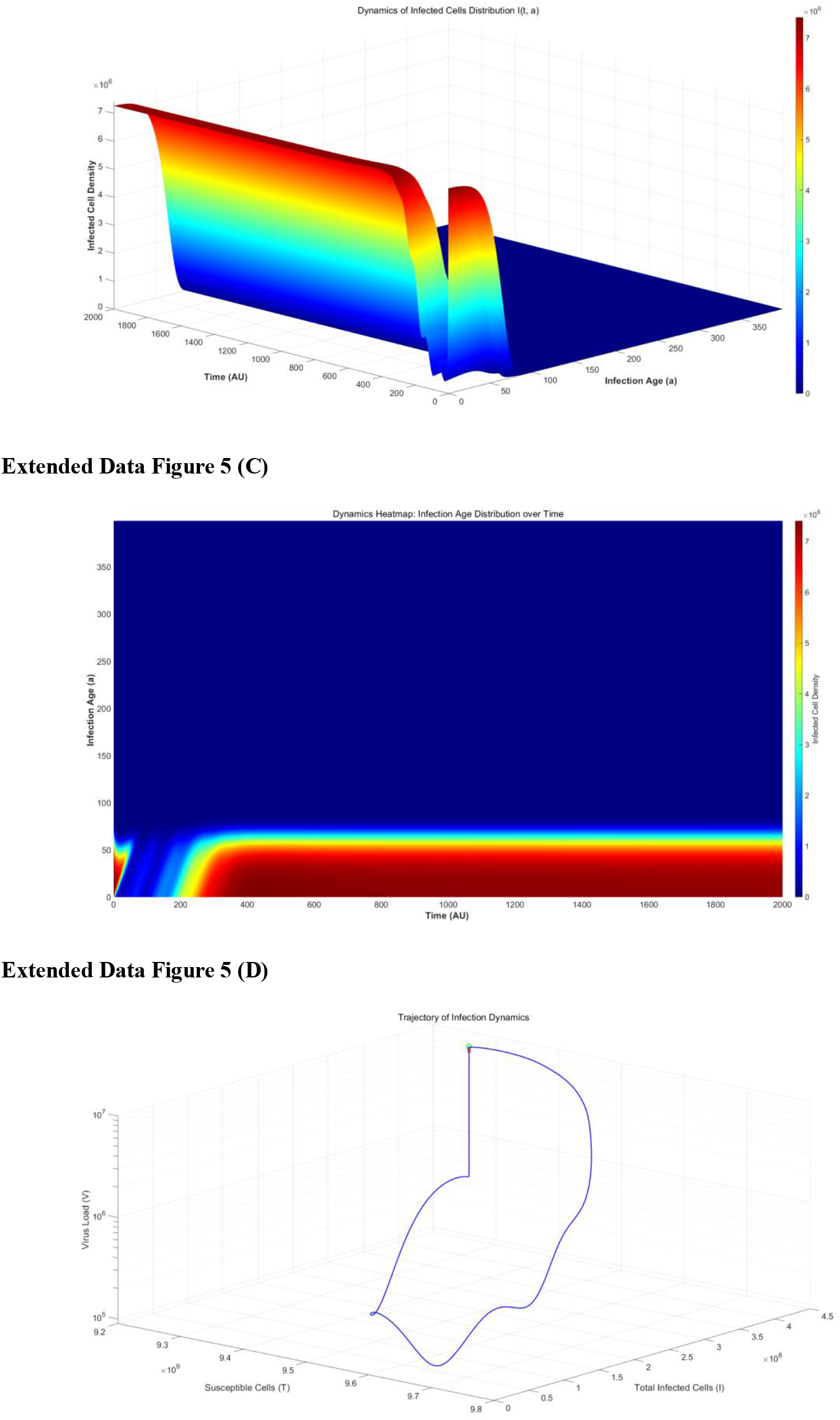
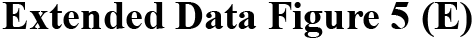
Transient viral suppression and rebound following sub-threshold monoclonal antibody therapy. **(A)** Temporal dynamics of host and viral compartments during administration of a high-affinity monoclonal antibody (mAb, 91st antibody quality parameters) at a dosage of 8×10^17^ molecules (approximately 2% of total host IgG). Panels illustrate (left to right): the trajectory of susceptible and infected cells, showing a transient decline in infected populations followed by a resurgence; free virus (*V*) kinetics, which drop significantly post-treatment but rebound to a persistent low-level steady state as antibody concentration decays; CD8++ T cell proliferation; and the decomposition of cell lysis rates. Notably, ADCC and Tc-mediated lysis peak initially due to mAb activity but subsequently decline, failing to sustain clearance. **(B)** Kinetics of immune complex formation and free antibody levels. Plots display IgM-Virus complexes, IgG-Virus complexes, free IgM, and free IgG. The administration of the specific mAb causes an initial spike in IgG-Virus complexes; however, as the exogenous antibody decays, the system reverts to a state characterized by persistent production of complexes, consistent with the re-establishment of chronic infection. **(C, D)** Spatiotemporal analysis of infected cell dynamics under treatment. **C**, Three-dimensional surface plot and **D**, corresponding two-dimensional heatmap representing infected cell density *I*(*t,a*) as a function of time (*t*) and infection age (*a*). The visualization clearly demarcates a therapeutic window with reduced infected cell density (0 –100 AU), followed by a repopulation of infected cells across age groups, visually confirming the “rebound” phenomenon. **(E)** Three-dimensional phase portrait of the infection trajectory involving free virus ( *V*), susceptible cells (*T*), and total infected cells. The green circle denotes the initial pre-treatment state (a stable chronic infection equilibrium). Upon mAb administration, the system undergoes a phase excursion (reduction in *V* and infected cells) but fails to transition to a cure state. The trajectory eventually returns to the red cross (×), which virtually overlaps with the initial point, indicating that the therapy provided only temporary suppression and the system relaxed back to the original chronic equilibrium.

**Extended Data Figure 6.**
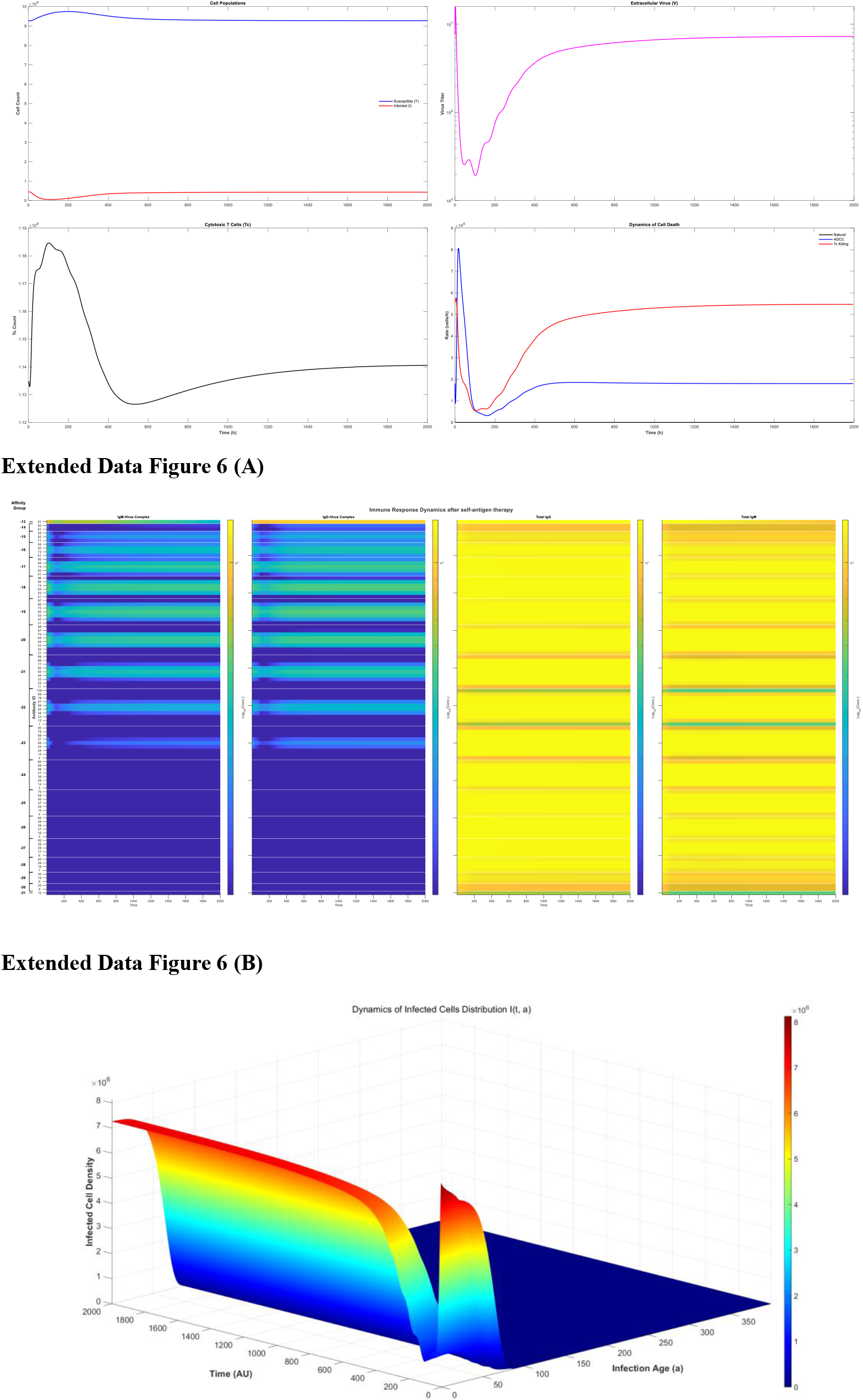
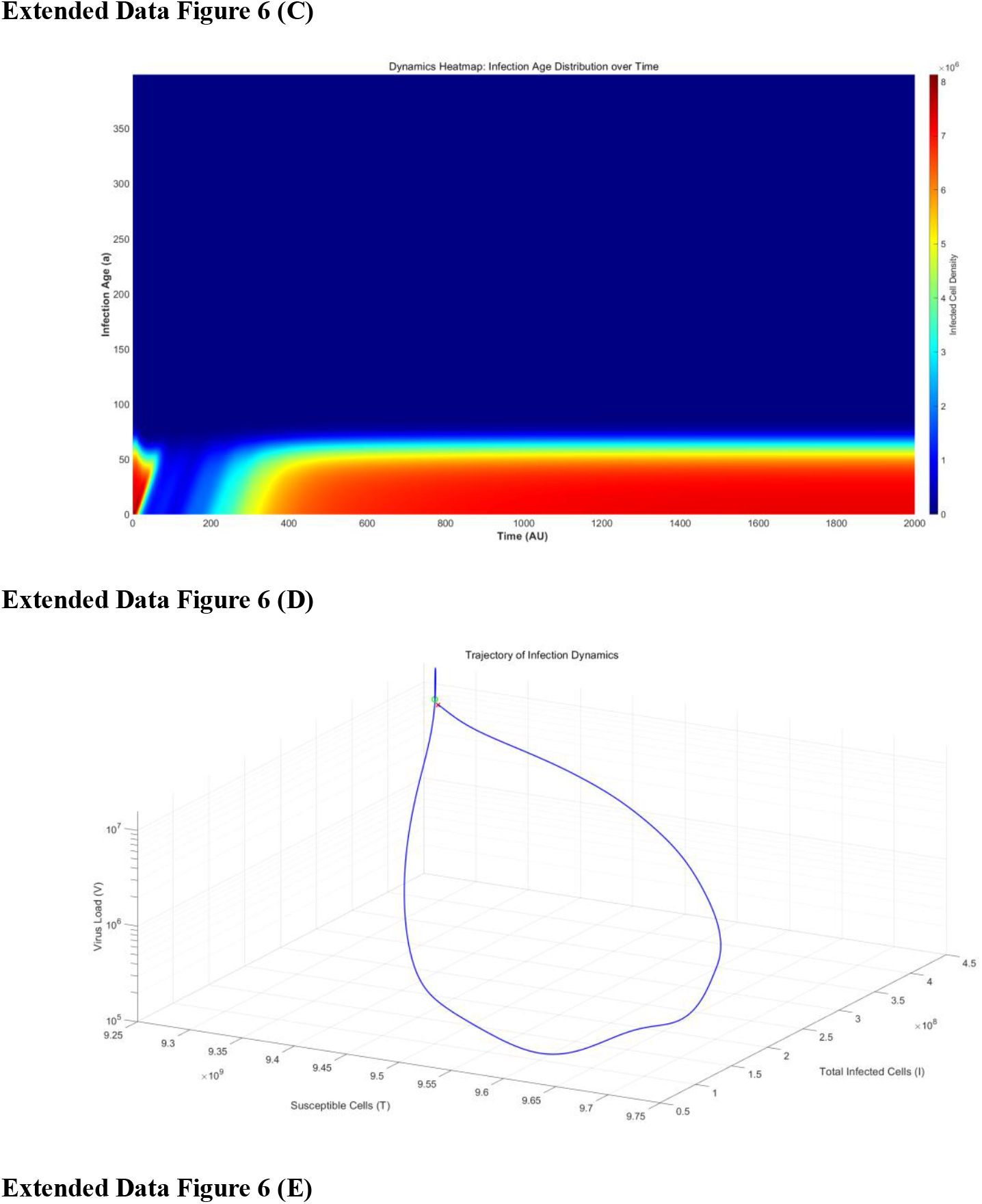
Transient viral suppression and rebound following environmental antigen augmentation. (A) Temporal dynamics of host and viral compartments during environmental antigen augmentation (simulating non-specific polyclonal boosting). The intervention involves a ∼450-fold increase in self-antigen concentration (from 2.2×10^17^ to 1×10^20^ molecules) via methods such as tissue stimulation. Panels display (from left to right): the trajectories of susceptible and infected cells, showing a significant but temporary reduction in infected burden; free virus (*V*) kinetics, which sharply decline upon treatment initiation but subsequently rebound to a low-level steady state; CD8++ T cell proliferation; and the decomposition of cell lysis rates. Note that ADCC and Tc-mediated lysis exhibit a transient spike correlating with antigen elevation but decline as the system relaxes back to the chronic state. **(B)** Kinetics of immune complex formation and free antibody levels. Plots show IgM-Virus complexes, IgG-Virus complexes, free IgM, and free IgG (left to right). Following the perturbation, high-avidity antibody-virus complexes fluctuate—initially increasing before dipping and finally rising again—to stabilize at high levels characteristic of persistent chronic infection. **(C, D)** Spatiotemporal analysis of infected cell dynamics under antigen augmentation. **C**, Three-dimensional surface plot and **D**, corresponding two-dimensional heatmap of infected cell density *I*(*t,a*) as a function of time (*t*) and infection age (*a*). The visualization clearly demarcates a therapeutic window where infected cell density drops (0 –100 AU), followed by a broad resurgence across infection ages, confirming the “rebound” phenomenon. **(E)** Three-dimensional phase portrait of the infection trajectory illustrating the relationship between free virus (*V*), susceptible cells (*T*), and total infected cells. The green circle marks the initial pre-treatment state (a stable chronic equilibrium with non-zero viral load). The therapeutic intervention drives a phase excursion (reduction in *V* and infected cells), but the trajectory ultimately loops back to the red cross ( ×), which overlaps with the initial point. This indicates that the antigen augmentation was insufficient to cross the threshold for eradication, resulting in a return to the original chronic equilibrium.

**Extended Data Figure 7.**
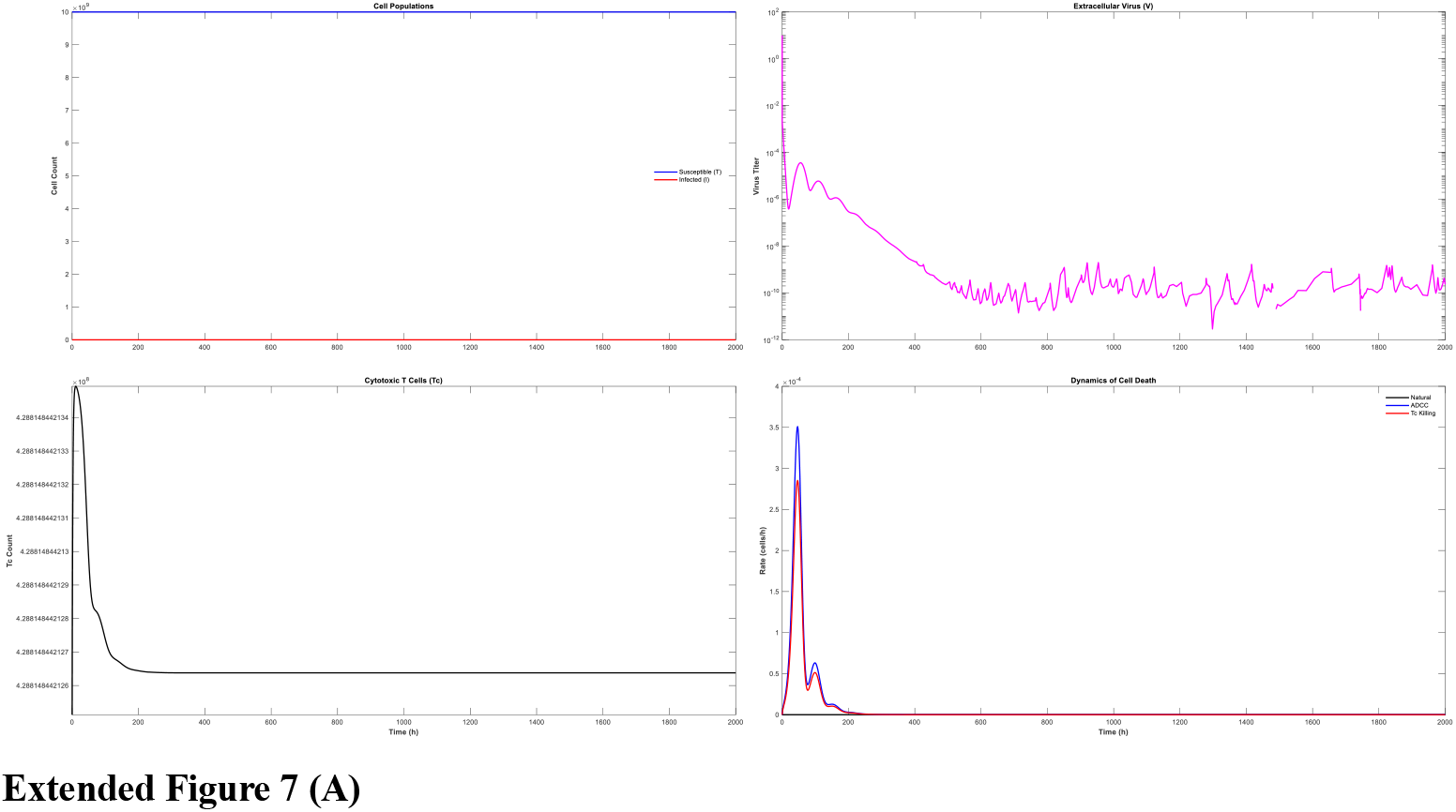
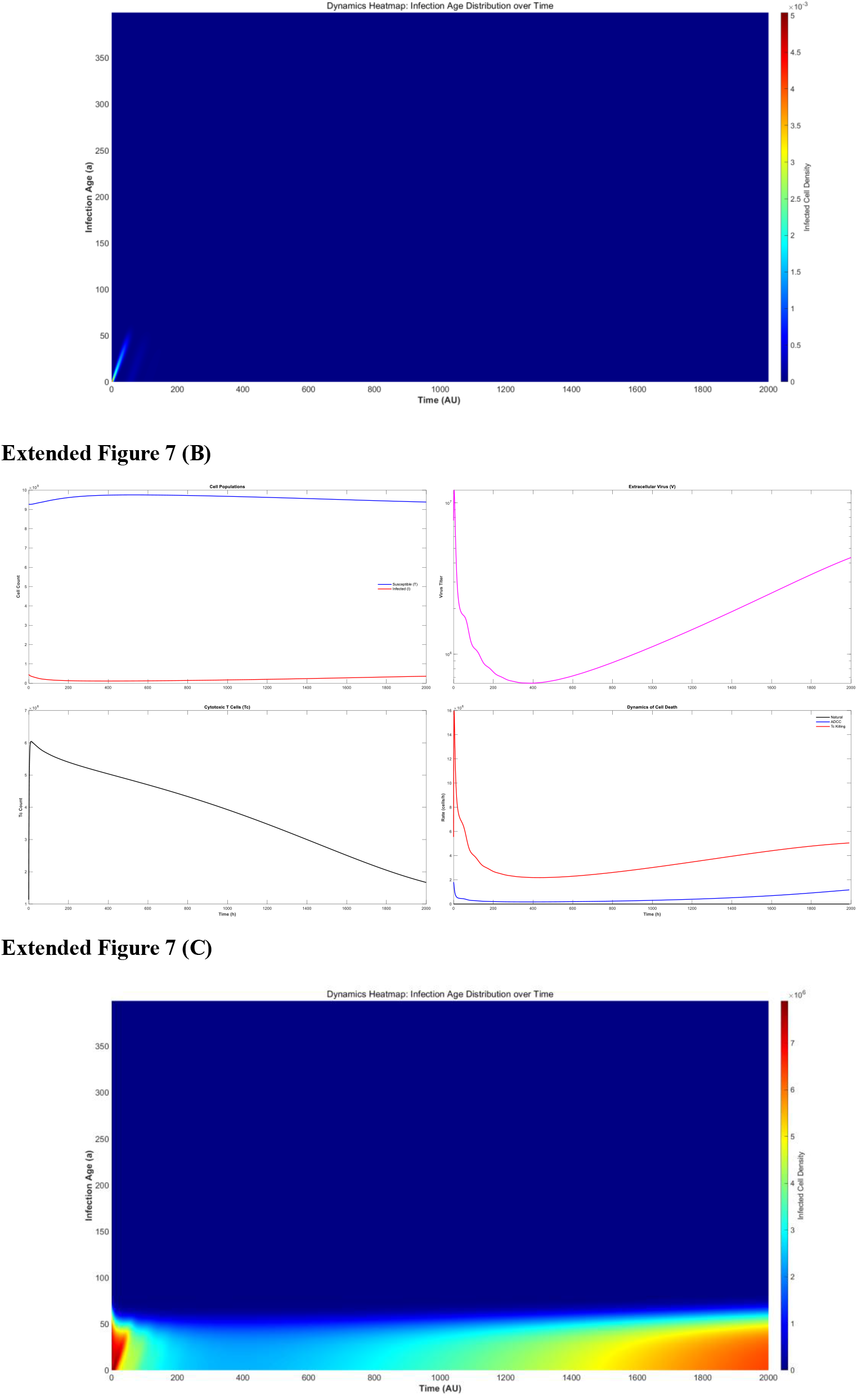
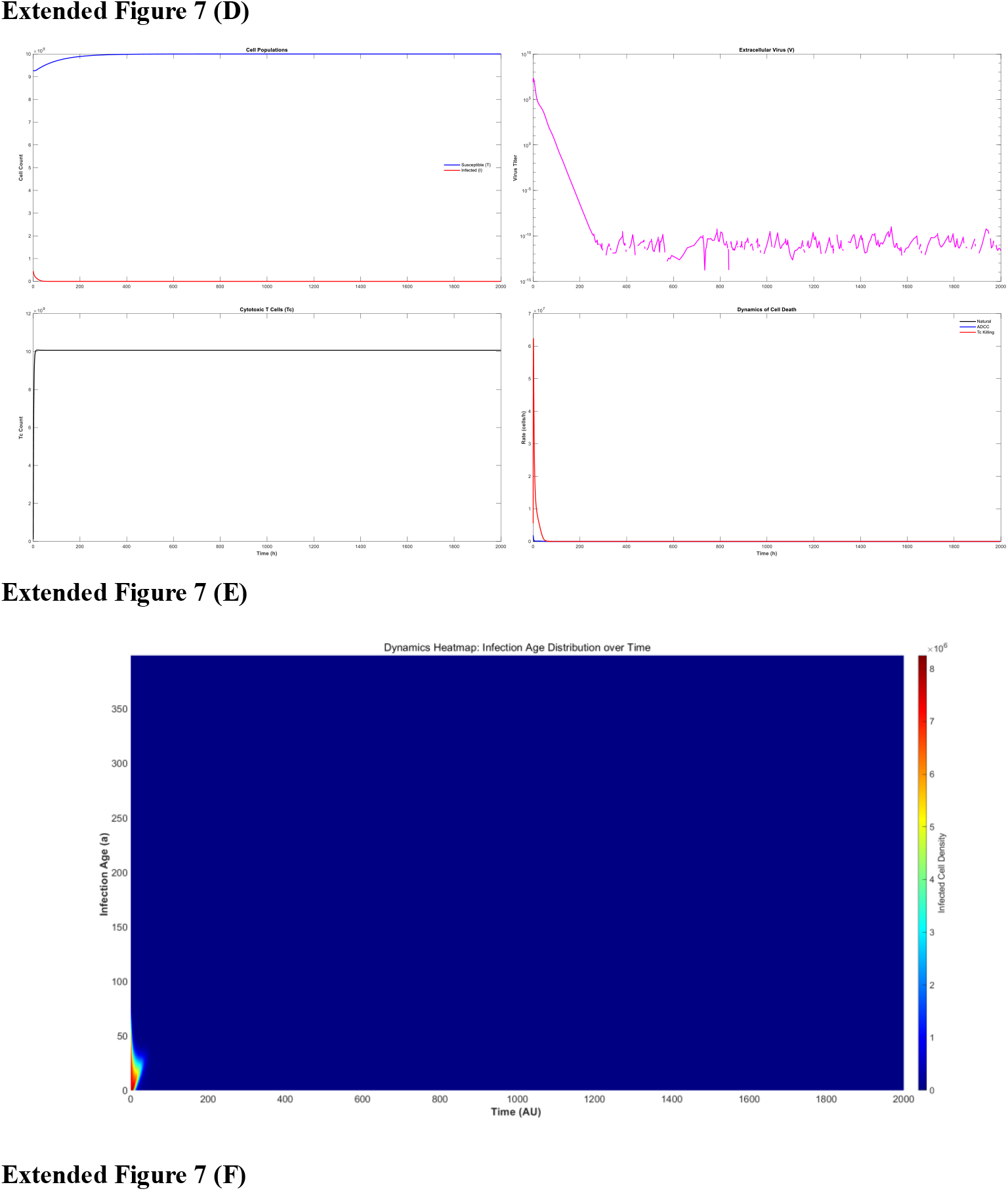
Dose-dependent thresholds for prophylactic versus therapeutic vaccination efficacy. **(A, B)** Low-dose immunization confers effective prophylaxis. Simulation of a viral challenge (*t*=1000) following priming with a low-dose viral antigen (5×10^12^). **A**, Temporal dynamics of cell populations, viral load, and immune effector mechanisms. The pre-existing immunity induced by the low-dose regimen effectively neutralizes the inoculum, preventing viral proliferation and limiting cytolysis (ADCC/CTL) to negligible levels. **B**, Two-dimensional heatmap of infected cell density *I*(*t,a*) confirms the complete blockade of infection establishment, demonstrating that a low antigenic threshold is sufficient for localized sterilization prior to viral expansion. **(C, D) Low-dose therapeutic intervention fails to reverse chronicity**. Administration of the same low-dose antigen (5×10^12^) in the context of an established chronic infection. **C**, Although the vaccine induces a transient suppression of viral load and infected cell numbers (panels 1-2), the boosted immune response (panels 3-4) is insufficient to overcome the established equilibrium. The system exhibits a “rebound” phenomenon, returning to a stable chronic plateau. **D**, the spatiotemporal projection illustrates the persistence of the infected cell reservoir, indicating that prophylactic-level doses are ineffective at clearing established viral reservoirs. **(E, F) High-dose therapeutic vaccination drives complete viral eradication**. Administration of a high-dose antigen bolus (1×10^14^, significantly exceeding the prophylactic dose) during chronic infection. **E**, the elevated antigenic load triggers a potent, high-amplitude spike in CTL proliferation and ADCC activity (bottom panels). This robust response successfully drives the viral load and infected cell population to extinction (*V*→0, I→0) without subsequent rebound. **F**, the heatmap visualizes the rapid and comprehensive elimination of infection cohorts across all ages. Validating the model prediction that the antigenic threshold required to break established immune tolerance and achieve a cure is magnitude-orders higher than that required for prevention.

## Notes

### Competing Interest Statement

The authors have declared no competing interest.

