## supplementary materials for "Integrated Modeling of BCR/TCR Repertoire Diversity Reveals the Mechanistic Basis of Immune Imprinting and Chronic Infection Control"

### 1 Mathematical Analysis and Dynamics of Model 3.1.1

#### 1.1. System Description and Positivity

The interaction between the virus (*V*), antibodies (*A*), and viral-antibody complexes (*C*) is governed by the following system of ordinary differential equations (ODEs):

$$\frac{dV}{dt} = k_0V - k_1VA + k_2C;$$

$$\frac{dA}{dt} = -k_1VA + k_2C + k_3C - k_4A + \pi;$$

$$\frac{dC}{dt} = k_1VA - k_2C - k_5C;$$

**Positivity of Solutions:**

To ensure biological feasibility, state variables must remain non-negative. Integrating the system from non-negative initial conditions:

- At V=0, $\frac{dV}{dt} = k_2C \geq 0$.
- At A=0, $\frac{dA}{dt} = k_2C + k_3C + \pi > 0$ (assuming $\pi$>0).
- At C=0, $\frac{dC}{dt} = k_1VA \geq 0$.

  Thus, the invariant region $\mathbb{R}^3_+$ is positively invariant, guaranteeing the biological validity of the model.

#### 1.2. Equilibrium Analysis

The system admits two distinct equilibria:

1. **Disease-Free Equilibrium (DFE), $E_0$:**

$$E_0 = (0, \frac{\pi}{k_4}, 0)$$

2. **Endemic Equilibrium (EE), $E^*$:**

$$V^* = \frac{k_0(k_2 + k_5)k_4 - \pi k_1k_5}{k_0k_1(k_3 - k_5)}, A^* = \frac{k_0(k_2 + k_5)}{k_1k_5}, C^* = \frac{k_0V^*}{k_5}$$

The existence of a biologically meaningful $E^*$ (where $V^*$>0) depends on the parameter threshold derived below.

#### 1.3. Stability Analysis

##### 1.3.1 Disease-Free Equilibrium Stability

The Jacobian matrix evaluated at $E_0$ is:

$$J(E_0) = \begin{pmatrix} k_0 - k_1\pi/k_4 & 0 & k_2 \\ -k_1\pi/k_4 & -k_4 & k_2 + k_3 \\ k_1\pi/k_4 & 0 & -k_2 - k_5 \end{pmatrix}$$

The eigenvalues are $\lambda_1 = k_0 - k_1\pi/k_4$, $\lambda_2 = -k_4$, and $\lambda_3 = -k_2 - k_5$.

Since $\lambda_2$ and $\lambda_3$ are always negative, the stability of $E_0$ is determined solely by $\lambda_1$.

- If $k_0 < k_1\pi/k_4$ , the DFE is asymptotically stable (viral clearance).
- If $k_0 > k_1\pi/k_4$ , the DFE is unstable.

**Biological Implication:** In most viral infections, the viral replication rate $k_0$ implies $k_0 > k_1\pi/k_4$ (given the typically small value of the association rate $k_1$). Mathematically, this prevents asymptotic convergence to V=0. However, biological systems are discrete rather than continuous. While mathematical variables approach zero asymptotically, biological components effectively vanish below a certain concentration. We address this in simulations by applying a discrete extinction threshold: if $V(t)<\epsilon$, then $V(t)\to 0$.

##### 1.3.2 Endemic Equilibrium Stability

The Jacobian matrix at the endemic equilibrium E∗$E*$ is:

$$J(E^*) = \begin{pmatrix} k_0 - k_1 A^* & -k_1 V^* & k_2 \\ -k_1 A^* & -k_1 V^* - k_4 & k_2 + k_3 \\ k_1 A^* & k_1 V^* & -k_2 - k_5 \end{pmatrix} = \begin{pmatrix} a & b & c \\ d & e & f \\ g & h & k \end{pmatrix}$$

The characteristic equation is $\lambda^3 - T\lambda^2 + M\lambda - D = 0$. Stability is analyzed via the Routh-Hurwitz criteria, where:

- **Trace (*T*):** $T=a+e+k$
- **Sum of principal minors (*M*):** $M=(ae-bd)+(ak-cg)+(ek-fh)$
- **Determinant (*D*):** $D=\det(J)=a(ek-fh)-b(dk-fg)+c(dh-eg)$

For asymptotic stability, specific conditions equivalent to the Routh-Hurwitz criteria (e.g., $T<0$, $D<0$, and $TM>D$) must be satisfied simultaneously.

#### 1.4. Numerical Simulations and Biological Interpretation

##### 1.4.1 Viral Clearance (Figure S1A)

Using the parameter set in **Table S1**, the condition $k_0 > k_1\pi/k_4$ holds, rendering the DFE mathematically unstable. Stability analysis of the endemic point ($V^*$, $A^*$, $C^*$) ≈ ($3.3\times10^4$,$10^7$,$6.7\times10^4$) yields eigenvalues with positive real parts ($Re(\lambda_{2,3})>0$), indicating the system oscillates or diverges without reaching a steady state.

However, incorporating the biological extinction threshold allows the simulation to demonstrate complete viral clearance (Figure S1A). This highlights the disconnect between continuous deterministic models and discrete biological realities: effective clearance occurs even when the DFE is mathematically unstable.

**Table S1: Parameters for Complete Clearance (Model 3.1.1)**

| Variable | Value | Parameter | Value |
|---|---|---|---|
| $V_0$ | 1 | $k_0$ (Replication) | 1 |
| $A_0$ | $10^3$ | $k_1$ (Association) | $10^{-7}$ |
| $C_0$ | 0 | $k_2$ (Dissociation) | $10^{-14}$ |
| | | $k_3$ (Feedback) | 2 |
| | | $k_4$ (Decay A) | $10^{-2}$ |
| | | $k_5$ (Decay C) | 0.5 |
| | | $\pi$ (Production) | 10 |

##### 1.4.2 Incomplete Clearance and Model Limitations (Figure S1B)

Under the parameters in **Table S2**, the system achieves a stable endemic equilibrium (Calculated $E^*$ eigenvalues have negative real parts: $\lambda_1\approx-0.54$, $\lambda_{2,3}\approx-0.0015\pm0.044i$). While the model can mathematically represent a persistent state (Figure S1B), it assumes well-mixed kinetics. In vivo chronic infections often rely on spatial compartmentalization (e.g., viral sanctuaries) not captured here. Consequently, we introduce Model 3.4.1 in the main text to better describe the mechanisms of chronic infection via cellular isolation.

**Table S2: Parameters for Incomplete Clearance**

| Variable | Value | Parameter | Value |
|---|---|---|---|
| $V_0$ | 1 | $k_0$ (Replication) | 1.2 |
| $A_0$ | $10^7$ | $k_1$ (Association) | $10^{-7}$ |
| $C_0$ | 0 | $k_2$ (Dissociation) | $10^{-14}$ |
| | | $k_3$ (Feedback) | 2 |
| | | $k_4$ (Decay A) | $10^{-2}$ |
| | | $k_5$ (Decay C) | 0.5 |
| | | $\pi$ (Production) | $10^5$ |

##### 1.4.3 Antibody Exhaustion and Immunosenescence (Figures S1C–E)

We defined "antibody exhaustion" as a kinetic regime where the viral replication rate exceeds the immune system's compensatory capacity. This is governed by three critical parameters:

1. **$k_0$ (Viral Load):** High replication rates.
2. **$k_3$ (Immune Feedback):** Represents CD4+ T cell activity (declines with aging/immunosenescence).
3. **$k_5$ (Clearance):** Represents antigen-antibody complex degradation (NK cell activity).

Combinatorial analysis (Table S3) demonstrates that high $k0$, low $k3$, or high $k5$ leads to a failure of antibody proliferation to match viral consumption. This manifests as a "germinal center proliferation defect," consistent with severe clinical outcomes. In this regime, the system is driven toward unstable dynamics where viral proliferation becomes unbounded (Figures S1C, S1D, and S1E).

**Table S3: Parameter Variations for Antibody Exhaustion Scenarios**

| Parameter | Setup 1 (High $k_5$) | Setup 2 (High $k_0$) | Setup 3 (Low $k_3$) |
|---|---|---|---|
| $k_0$ | 1 | **2** | 1 |
| $k_3$ | 2 | 2 | **1** |
| $k_5$ | **1** | 0.5 | 0.5 |
| $\pi$ | $10^4$ | $10^4$ | $10^4$ |
| *(Other parameters remain constant: $V_0=1$, $A_0=10^3$, $k_1=10^{-7}$, $k_2=10^{-14}$, $k_4=10^{-2}$)* | | | |

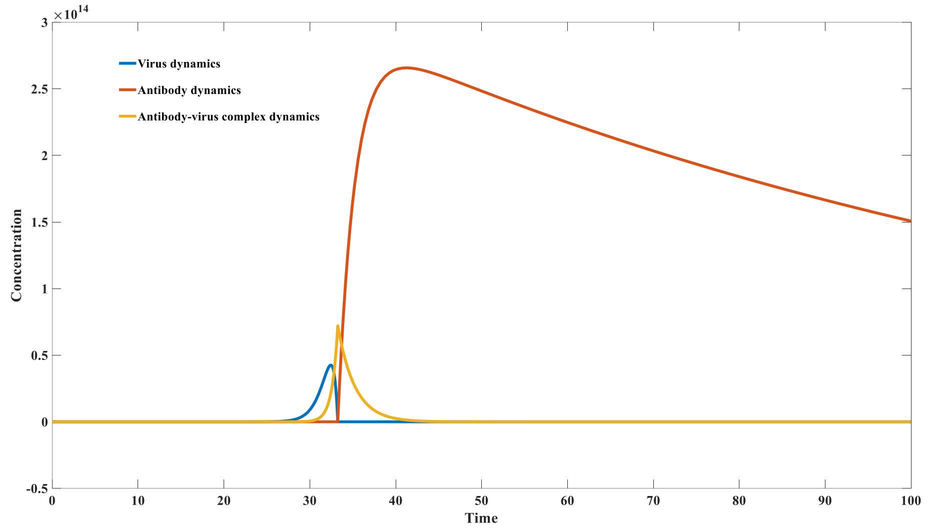


Figure S_1A: Dynamics of antibody - virus interaction （complete virus clearance）

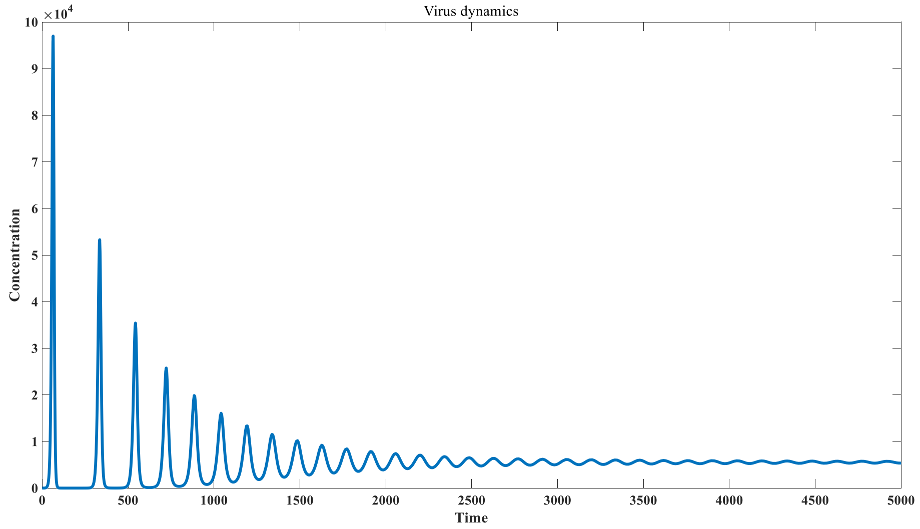


Figure S_1B: Dynamics of antibody - virus interaction (incomplete virus clearance)

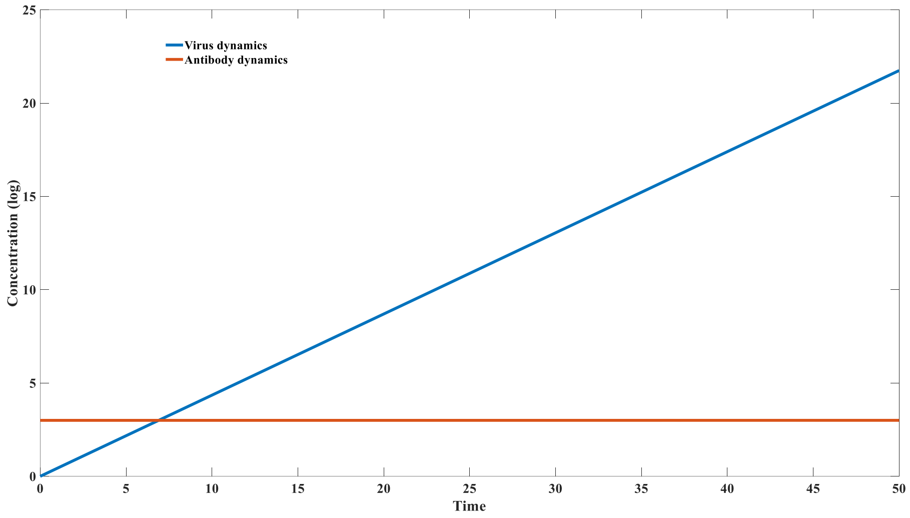


Figure S_1C: Defective antibody production due to fast clearance rate of antibody - virus complex ($k_5 = 1$)

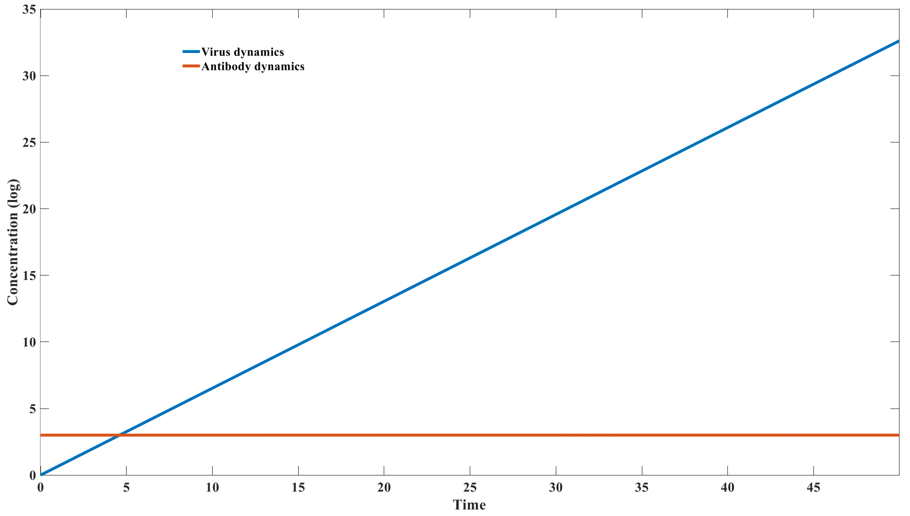


Figure S_1D: Defective antibody production due to fast virus replication rate ($k_0 = 1.5$)

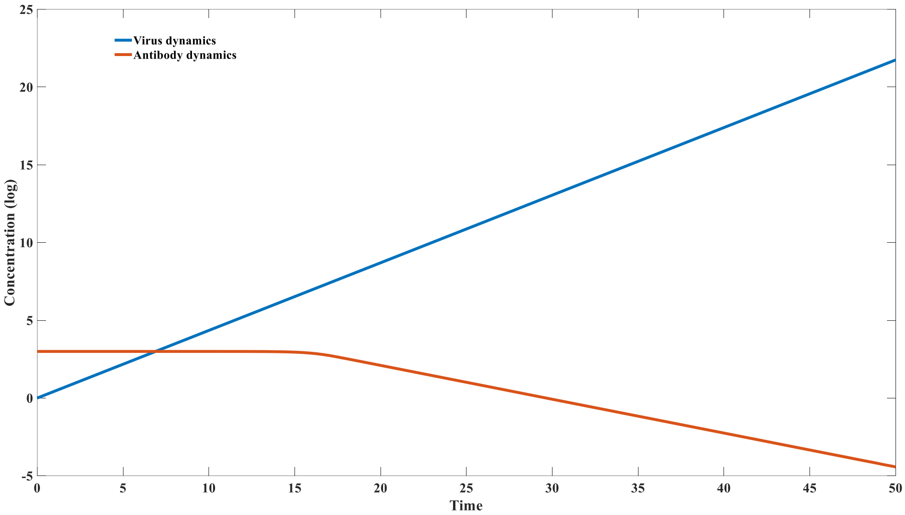


Figure S_1E: Defective antibody production due to slow antibody regeneration rate ($k_3 = 1$)

#### 2 Mathematical Analysis and Parameterization of Model 3.1.2

##### 2.1. Rationale and Model Reconstruction

A primary limitation of Model 3.1.1 is its inability to recapitulate long-term immune memory. In Model 3.1.1, following viral clearance, specific neutralizing antibody titers revert to the naïve, disease-free equilibrium without permanent upregulation. This contradicts *in vivo* observations where memory B cells persist, maintaining elevated IgG levels and providing sustained protection. To resolve this, Model 3.1.2 introduces environmental antigens ($E$) as a mechanism for homeostatic maintenance and feedback-driven antibody regeneration.

The governing ordinary differential equations (ODEs) for Model 3.1.2 are:

$$\frac{dV}{dt} = k_0V - k_1VA + k_2C;$$

$$\frac{dA}{dt} = -k_1VA + k_2C + k_3C - k_4A - p_1EA + p_2Q + p_3Q;$$

$$\frac{dC}{dt} = k_1VA - k_2C - k_5C;$$

$$\frac{dE}{dt} = \pi - p_1EA + p_2Q;$$

$$\frac{dQ}{dt} = p_1EA - p_2Q - p_4Q;$$

**Dynamical definition of Antibody Homeostasis:**

Unlike Model 3.1.1, which assumes a fixed basal influx of antibodies, Model 3.1.2 treats antibody supply as a dynamic process governed by interactions with environmental antigens.

- $-p_1EA$: Represents the binding of antibodies ($A$) to environmental antigens ($E$) to form the complex $Q$.
- $p_2Q$: Represents the dissociation of the complex back into free antibodies.
- $p_3Q$: The critical feedback term representing complex-stimulated antibody regeneration (mimicking memory B cell activation signals).

##### 2.2. Positivity and Invariance

To ensure biological feasibility, we verified the non-negativity of the system. Examining the boundary conditions of the non-negative orthant $\mathbb{R}_+^5$:

- At $V$=0, $\dot{V} = k_2 C \geq 0$.
- At $A$=0, $\dot{A} = k_2 C + k_3 C + p_2 Q + p_3 Q \geq 0$.
- At $C$=0, $\dot{C} = k_1 V A \geq 0$.
- At $E$=0, $\dot{E} = \pi + p_2 Q \geq 0$.
- At $Q$=0, $\dot{Q} = p_1 E A \geq 0$.

Thus, for any non-negative initial condition, the system trajectories remain consistently non-negative.

**2.3. Equilibrium and Stability Analysis**

In the absence of viral load ($V$=0 and $C$=0), the system acts as a reduced subsystem governing antibody homeostasis:

$$\frac{dA}{dt} = -k_4 A - p_1 E A + p_2 Q + p_3 Q;$$

$$\frac{dE}{dt} = \pi - p_1 E A + p_2 Q;$$

$$\frac{dQ}{dt} = p_1 E A - p_2 Q - p_4 Q;$$

Using the parameter set provided in **Table S4**, we calculated the initial conditions for the system stability analysis: ($A_0=10^7$; $E_0=10^8$; $Q_0=2\times10^5$).

**Jacobian Analysis:**

The Jacobian matrix $J$ of the full system evaluated at an equilibrium point ($V^*$,$A^*$,$C^*$,$E^*$,$Q^*$) is defined as:

$$J=\begin{pmatrix} k_0 - k_1 A^* & -k_1 V^* & k_2 & 0 & 0 \\ -k_1 A^* & -k_1 V^* - k_4 - p_1 E^* & k_2 + k_3 & -p_1 A^* & p_2 + p_3 \\ k_1 A^* & k_1 V^* & -k_2 - k_5 & 0 & 0 \\ 0 & -p_1 E^* & 0 & -p_1 A^* & p_2 \\ 0 & p_1 E^* & 0 & p_1 A^* & -p_2 - p_4 \end{pmatrix}$$

We identified two distinct equilibrium states:

1. **Disease-Free Equilibrium (DFE):**

   State vector: ($A^*=10^7$, $E^*=10^8$, $Q^*=2\times10^5$, $V^*=0$, $C^*=0$).

   Eigenvalues:

   $\lambda_{1..5}$= {0.9, −0.5197, −0.5, 0.0087, $4.1\times10^{-18}$}

   The presence of positive eigenvalues ($\lambda_1$,$\lambda_4$) indicates that the DFE is mathematically unstable (a saddle point) under continuous dynamics.

2. **Endemic Equilibrium (EE):**

   State vector: ($A^*=10^8$, $E^*=10^7$, $Q^*=2\times10^5$, $V^*=3\times10^5$, $C^*=6\times10^5$).

   Eigenvalues:

   $\lambda_{1..5}$={−0.5353, −0.5000, 0.0111±0.0911i, −0.0100}

**Constraint-Driven Clearance:**

Although linear stability analysis suggests the disease-free state is unstable (implying potential viral recrudescence or persistence), numerical simulations (Figure S2) demonstrate complete viral clearance. This discrepancy is resolved by imposing **discontinuous constraints** on the viral population $V$. In the biological context, when the viral load drops below a critical threshold (representing extinction), it is forced to zero, preventing the mathematical regeneration driven by the positive eigenvalue $\lambda_1$ . This hybrid dynamical approach allows the system to achieve a stable disease-free state indistinguishable from real-world clearance.

##### 2.4. Parameter Sets

**Table S4 | Parameter definitions and initial values for Model 3.1.2**

| Compartment | Initial Value ($t$=0) | Parameter | Value | Description |
|---|---|---|---|---|
| $V_0$ | 1 | $k_0$ | 1 | Viral replication rate |
| $A_0$ | $10^7$ | $k_1$ | $10^{-8}$ | Virus-Antibody association rate |
| $C_0$ | 0 | $k_2$ | $10^{-14}$ | Complex dissociation rate |
| $E_0$ | $10^8$ | $k_3$ | 2 | Complex clearance rate |
| $Q_0$ | $2\times10^5$ | $k_4$ | $10^{-2}$ | Antibody decay rate |
| | | $k_5$ | 0.5 | Complex decay rate |
| | | $\pi$ | $10^4$ | Antigen influx rate |
| | | $p_1$ | $10^{-10}$ | Env. Antigen-Ab association rate |
| | | $p_2$ | 0 | Env. Complex dissociation rate |
| | | $p_3$ | 1 | Feedback regeneration rate |
| | | $p_4$ | 0.5 | Env. Complex decay rate |

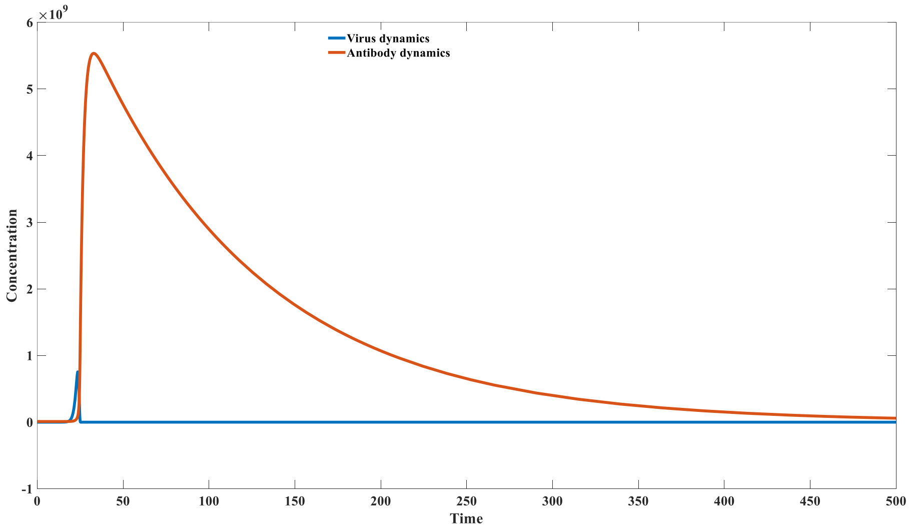


Figure S_2: Virus-antibody interaction dynamics using model 3.1.2

#### 3 Mathematical Analysis and Parameter Setting of Model 3.1.3 (Polyclonal Response)

##### 3.1. Rationale and Model Extension

While Model 3.1.2 elucidates the role of environmental antigens in maintaining immune memory, it treats the antibody pool as a homogeneous population. To fully recapitulate the mechanisms of clonal expansion, we extended the framework to Model 3.1.3 by incorporating antibody diversity. This model remains parsimonious but introduces a multivariable vector of antibody clones ($A_i$), allowing for the study of inter-clonal competition and repertoire reshaping.

A critical modeling assumption governs the interaction with environmental antigens (*E*):

- **Environmental Antigens (*E*) as a Constant Background:** As *E* represents a broad, polydisperse category of background antigens (e.g., self-antigens or commensal markers), their interaction kinetics ($p_1$ and $p_2$) are assumed to be uniform across different antibody clones (mean-field approximation). This ensures that in the disease-free state, the total antibody pool is maintained at a homeostatic level determined by the fixed supply $\pi$.
- **Pathogen Specificity:** In contrast, interactions with the virus (*V*) are clone-specific. The association ($k_{1,i}$) and dissociation ($k_{2,i}$) rates vary by antibody type, driving the selective pressure.

The expanded system of differential equations for *n* distinct antibody clones is expressed as:

$$\frac{dV}{dt} = k_0 V - \sum_{n}^{i=1} k_{1,i} V A_i + \sum_{n}^{i=1} k_{2,i} C_i \,;$$

$$\frac{dA_i}{dt} = -k_{1,i} V A_i + k_{2,i} C_i + k_3 C_i - k_4 A_i - p_1 E A_i + p_2 Q_i + p_3 Q_i;$$

$$\frac{dC_i}{dt} = k_{1,i} V A_i - k_{2,i} C_i - k_5 C_i;$$

$$\frac{dE}{dt} = \pi - \sum_{n}^{i=1} p_1 E A_i + \sum_{n}^{i=1} p_2 Q_i \,;$$

$$\frac{dQ_i}{dt} = p_1 E A_i - p_2 Q_i - p_4 Q_i;$$

where $i \in \{1,...,n\}$ represents the index of the antibody clone.

##### 3.2. Simulation and Parameter Sets

To visualize clonal selection, we simulated the dynamics of $n$=4 distinct antibody clones. The specific parameters are detailed in **Table S5**. The initial conditions assume an equipotent distribution, where all four antibodies constitute 25% of the total pool ($2.5\times10^6$ each).

**Table S5 | Parameter sets and initial conditions for Model 3.1.3**

| **Parameter** | **Description** | **Antibody 1 (High $k_{on}$)** | **Antibody 2 (Low Affinity)** | **Antibody 3 (Med $k_{on}$)** | **Antibody 4 (Low $k_{on}$)** |
|---|---|---|---|---|---|
| **Initial Values** | ($t$=0) | | | | |
| $V_0$ | Virus | 1 | 1 | 1 | 1 |
| $A_{0,i}$ | Antibody | $2.5\times10^6$ | $2.5\times10^6$ | $2.5\times10^6$ | $2.5\times10^6$ |
| $C_{0,i}$ | V-A Complex | 0 | 0 | 0 | 0 |
| $E_0$ | Env. Antigen | $2.5\times10^7$ | $2.5\times10^7$ | $2.5\times10^7$ | $2.5\times10^7$ |
| $Q_{0,i}$ | E-A Complex | $5\times10^4$ | $5\times10^4$ | $5\times10^4$ | $5\times10^4$ |
| **Kinetics** | | | | | |
| $k_0$ | Viral Rep. | 1 | 1 | 1 | 1 |
| $k_{1,i}$ | Association | $10^{-8}$ | $10^{-8}$ | $0.9\times10^{-8}$ | $0.8\times10^{-8}$ |
| $k_{2,i}$ | Dissociation | $10^{-14}$ | 1 | $0.9\times10^{-14}$ | $0.8\times10^{-14}$ |
| $k_3$ | Clearance | 2 | 2 | 2 | 2 |
| $k_4$ | Ab Decay | $10^{-2}$ | $10^{-2}$ | $10^{-2}$ | $10^{-2}$ |

| Parameter | Description | Antibody 1 (High $k_{on}$) | Antibody 2 (Low Affinity) | Antibody 3 (Med $k_{on}$) | Antibody 4 (Low $k_{on}$) |
|---|---|---|---|---|---|
| $k_5$ | Complex Decay | 0.5 | 0.5 | 0.5 | 0.5 |
| **Env. Interactions** | | | | | |
| π | Influx | $10^4$ | $10^4$ | $10^4$ | $10^4$ |
| $p_1$ | Association | $10^{-10}$ | $10^{-10}$ | $10^{-10}$ | $10^{-10}$ |
| $p_2$ | Dissociation | 0 | 0 | 0 | 0 |
| $p_3$ | Feedback | 1 | 1 | 1 | 1 |
| $p_4$ | Decay | 0.5 | 0.5 | 0.5 | 0.5 |

**3.3. Results and Biological Interpretation**

Simulation results (Figure S3) illustrate the mechanisms of selective clonal expansion. We emphasize two key observations:

1. **Affinity-Driven Selection:**
   Comparing Antibody 1 and Antibody 2, which share the same forward association rate ($k_1$ ), Antibody 1 exhibits a significantly lower dissociation rate ($k_2=10^{-14}$) compared to Antibody 2 ($k_2=1$). Consequently, Antibody 1 proliferates rapidly, while Antibody 2 fails to expand. This confirms that clones with stronger binding usage (high affinity) are selectively amplified by the feedback mechanism governed by the V-A complex formation ($k_3 C_i$).
2. **Kinetic Rate Dominance under Isothermodynamic Conditions:**
   Antibodies 1, 3, and 4 possess identical thermodynamic equilibrium constants ($Kd=k_2/k_1=10^{-6}$). However, they differ in their kinetic on-rates ($k_{1,i}$). The simulation reveals that despite having the same binding energy, the clone with the highest association rate (Antibody 1 > 3 > 4) exhibits the fastest proliferation. This suggests that kinetic on-rates ($k_{on}$), rather than equilibrium affinity alone, are the primary drivers of acute clonal expansion in this dynamic system.

**Imprinting of the Antibody Repertoire:**

The simulation demonstrates a permanent "scarring" or reshaping of the antibody repertoire post-infection. While the system initiates with a uniform distribution (25% each), the post-infection steady state is asymmetrical: Antibody 1 stabilizes at 38%, Antibody 3 at 31%, Antibody 4 at 24%, and the low-affinity Antibody 2 is marginalized to 8%. Although the total antibody concentration returns to a homeostatic set-point determined by $\pi$, the **relative proportions** are permanently altered. This diversity shift, driven by the varying virus-interaction histories, encodes the "memory" of the infection within the population structure of the antibody pool.

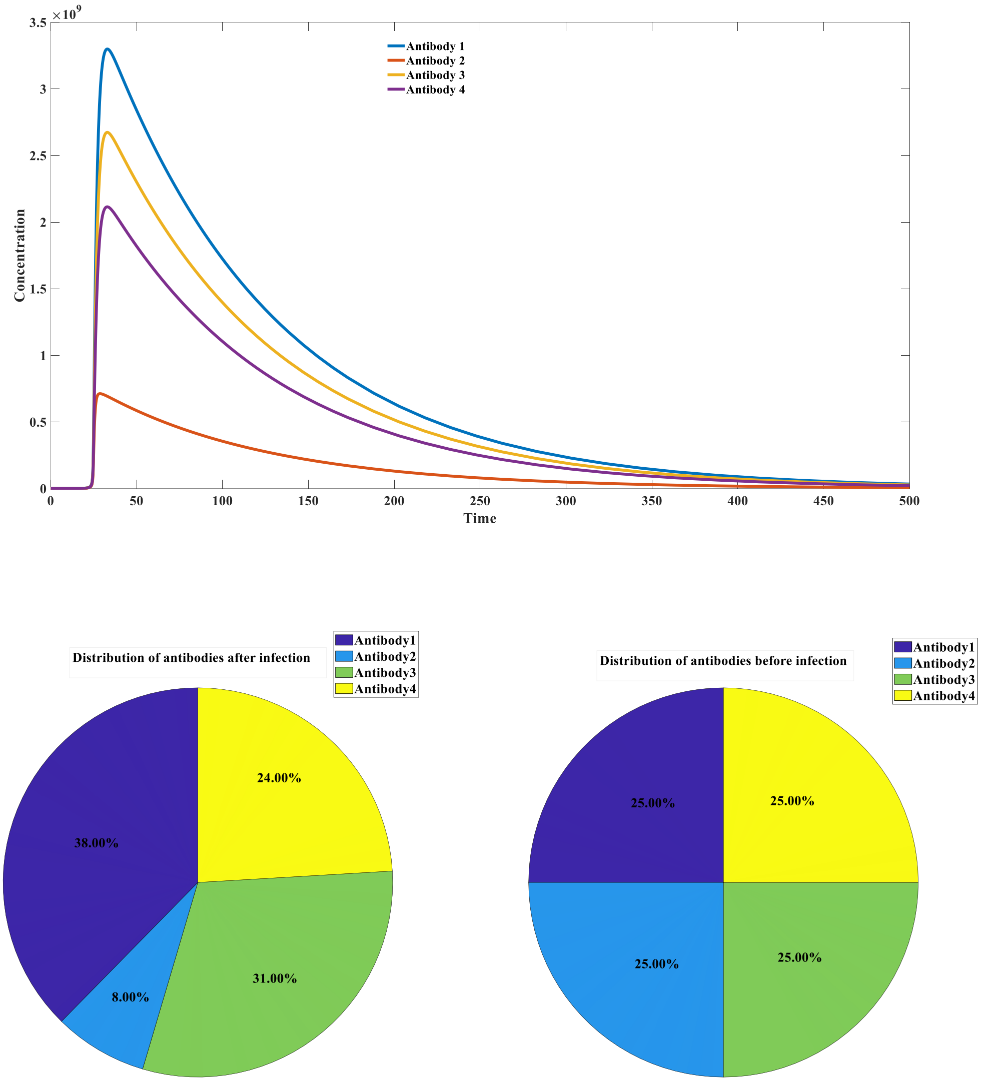


Figure S_3: Simulation of antibody dynamics using model 3.1.3

#### 4 Mathematical Modeling of Viral Rebound under Monoclonal Antibody Therapy (Model 3.1.4)

##### 4.1. Rationale and Model Structure

To investigate the mechanisms underlying viral rebound following passive immunization (monoclonal antibody therapy), we constructed Model 3.1.4. This model introduces an exogenous antibody species ($A_2$) alongside the host's endogenous antibody response ($A$).

The critical distinction in this model is the functional difference between endogenous and exogenous antibodies:

1. **Endogenous Antibodies ($A$):** Generated via the feedback loop $k_3C$, representing the host's active immune response (clonal expansion).
2. **Exogenous Antibodies ($A_2$):** Introduced via the dosing function $\pi(t)$. While $A_2$ can neutralize the virus ($V$) to form complexes ($C_2$), these exogenous complexes **do not** contribute to the regeneration of the host's antibody pool (i.e., the term $k_3C_2$ is absent

from the $\frac{dA_2}{dt}$ equations). Furthermore, exogenous antibodies are subject to metabolic clearance ($k_6$) independent of the host's immune homeostasis.

The governing equations are:

$$\frac{dV}{dt} = k_0V - k_1VA + k_2C - k_1VA_2 + k_2C_2;$$

$$\frac{dA}{dt} = -k_1VA + k_2C + k_3C;$$

$$\frac{dC}{dt} = k_1VA - k_2C - k_5C;$$

$$\frac{dA_2}{dt} = -k_1VA_2 + k_2C_2 - k_6A_2 + \pi(t);$$

$$\frac{dC_2}{dt} = k_1VA_2 - k_2C_2 - k_5C_2;$$

Here, $\pi(t)$ represents the administration of the monoclonal antibody drug.

##### 4.2. Parameter Sets and Initial Conditions

We simulated the viral dynamics under various dosing regimens. The standard parameters and initial conditions are detailed in **Table S6**. The decay rate of the exogenous antibody ($k_6$=0.05) is set significantly higher than natural homeostasis to reflect the pharmacokinetic half-life of the drug.

**Table S6 | Parameter definitions and initial values for Model 3.1.4**

| Compartment | Initial Value ($t$=0) | Parameter | Value | Description |
|---|---|---|---|---|
| **$V_0$** | 1 | $k_0$ | 0.5 | Viral replication rate |
| **$A_0$** | 10 | $k_1$ | $10^{-5}$ | Association rate (Endogenous & Exogenous) |
| **$C_0$** | 0 | $k_2$ | $10^{-14}$ | Dissociation rate (Endogenous & Exogenous) |
| **$A_{2,0}$** | 0 | $k_3$ | 1.5 | Endogenous Ab proliferation rate |
| **$C_{2,0}$** | 0 | $k_5$ | 0.5 | Complex clearance rate |
| | | $k_6$ | 0.05 | Exogenous mAb metabolic decay rate |
| | | $\pi_{dose}$ | $1.5\times10^6$ | Standard drug dosage (instantaneous) |

##### 4.3. Simulation Results: The "Kinetic Decoupling" Effect

##### A. Impact of Intervention Timing (Figure S4A)

We simulated the administration of a fixed dose ($1.5\times10^6$) of monoclonal antibodies at different time points relative to the infection onset (t=21 to t=30).

- **Early Intervention (t<28):** Administration effectively suppresses the viral peak. The residual virus is insufficient to trigger a rebound because the exogenous antibody maintains suppression until the endogenous system potentially matures, or the virus is cleared.
- **The Critical "Rebound" Window (t≈28–29):** A paradoxical viral rebound occurs. In this window, the mAb therapy suppresses the viral load rapidly, depriving the host's immune system of the antigen ($V$) required to stimulate the $k_3C$ feedback loop. Consequently, the

host remains immunologically naïve. Once the exogenous antibody $A_2$ metabolizes (decays via $k_6$) below a protective threshold, the residual virus resurges unchecked.

- **Late Intervention (t>29):** The therapy is administered after the endogenous response has already peaked. While it aids clearance, it does not induce rebound as the host immunity is already established.

**B. Impact of Dosage and the "Antigen Sink" (Figure S4B)**

We analyzed the dose-response relationship at a fixed intervention time (t=28). The results reveal a **non-monotonic response** to treatment dosage:

1. **Sub-therapeutic (<0.5×10$^6$):** The dose is insufficient to impact viral kinetics; viral proliferation continues unaffected.
2. **Pro-Rebound Range (1.0×10$^6$ –1.5×10$^6$):** This represents a "danger zone." The dosage is high enough to temporarily suppress viral load and sequester antigens, effectively "hiding" the virus from the host's adaptive immune system (preventing endogenous priming). However, the dose is too low to achieve sterilizing immunity. As the drug decays, the virus rebounds to high levels in the absence of host protection.
3. **Therapeutic/Sterilizing (>4.5×10$^6$):** The dosage exceeds the maximum threshold required for complete viral clearance. The virus is eliminated before the drug decays, preventing any potential for rebound.

This model demonstrates that viral rebound is a deterministic consequence of **"kinetic decoupling"**—a mismatch between the decay of passive immunity and the maturation of active immunity. Avoiding rebound requires either sufficiently early administration or dosages high enough to ensure complete clearance without reliance on the endogenous response.

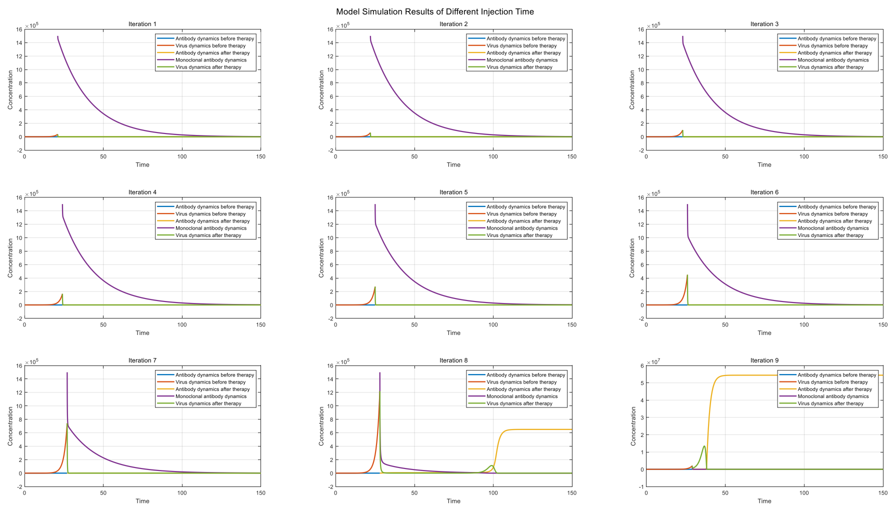


Figure S_4A: Simulation of host-virus interaction at different injection time of monoclonal antibody

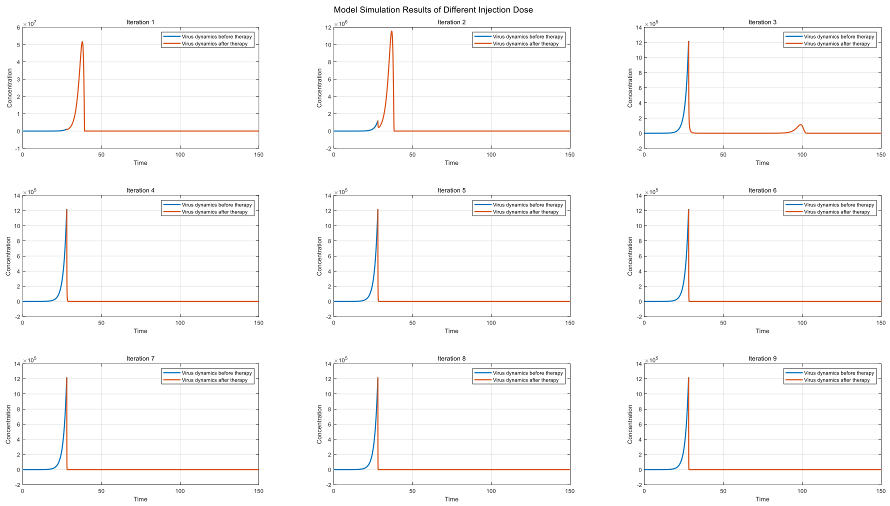


Figure S_4B: Simulation of host-virus interaction at different injection dose of monoclonal antibody

##### 4.4. Overview of the Standard Rebound Model (Perelson et al.)

A prevailing hypothesis, mathematically formalized by Perelson et al., postulates that viral rebound during monoclonal antibody (mAb) therapy is primarily driven by the *de novo* generation of resistant mutant strains during the infection course. The governing equations of this model are described as follows:

$$\frac{dT}{dt} = -\beta_1 V_1 T - \beta_2 V_2 T + \gamma T\left(1 - \frac{E_1 + E_2 + I_1 + I_2 + T}{T_M}\right);$$

$$\frac{dE_1}{dt} = -\beta_1 V_1 T - kE_1;$$

$$\frac{dE_2}{dt} = -\beta_2 V_2 T - kE_2;$$

$$\frac{dI_1}{dt} = kE_1 - \delta(t) I_1;$$

$$\frac{dI_2}{dt} = kE_2 - \delta(t) I_2;$$

$$\frac{dV_1}{dt} = \pi(1-\mu) I_1 - k_{on,1} V_1 A + k_{off,1} C_1 - cV_1;$$

$$\frac{dC_1}{dt} = k_{on,1} V_1 A - k_{off,1} C_1 - \gamma_1 C_1;$$

$$\frac{dV_2}{dt} = \pi I_2 + \pi\mu I_1 - k_{on,2} V_2 A + k_{off,2} C_2 - cV_2;$$

$$\frac{dC_1}{dt} = k_{on,2} V_2 A - k_{off,2} C_2 - \gamma_2 C_2;$$

The model utilizes discontinuous functions to approximate the dynamics of infected cell death $\delta(t)$ and antibody concentration $A(t)$:

$$\delta(t) = \begin{cases} \delta & t < t^* \\ \delta_M - (\delta_M - \delta) e^{-\sigma(t - t^*)} & t \geq t^* \end{cases}$$

$$A(t) = \begin{cases} 0 & t < t_{inf} \\ \frac{A_{max}}{\Delta t}\left(t - t_{inf}\right) & t_{inf} \le t < t_{inf} + \Delta t \\ A_{max} e^{-\alpha\left(t-\left(t_{inf}+\Delta t\right)\right)} & t \ge t_{inf} + \Delta t \end{cases}$$

**4.5. Critique and Limitations**

While the Perelson model provides a foundational framework, we identified four critical theoretical limitations that necessitate the reconstruction of the immune-viral dynamics in our proposed model:

1. **Inaccuracy of the Incidence Term:** The term $-\beta VT$ assumes simple mass-action kinetics. However, given the significant size disparity between virions and target cells ($V \ll T$), a single cell is subject to multiple viral entry events simultaneously (high Multiplicity of Infection, MOI). A quadratic approximation ($V \cdot T$) tends to overestimate the rate of susceptible cell depletion, failing to account for cellular saturation or localized stoichiometry.
2. **Neglect of Endogenous vs. Exogenous Distinction:** The model aggregates neutralizing activity into a single variable A(t) defined by a forcing function. It fails to fundamentally distinguish between **endogenous autoantibodies** (BCRs) and **exogenous mAbs**. Endogenous antibodies participate in a positive feedback loop (antigen binding → B-cell activation → regeneration), whereas exogenous antibodies are purely decaying agents. Ignoring this regenerative capability underestimates the host's potential for spontaneous control.
3. **Phenomenological Approximation of Immune Dynamics:** By using piecewise functions for A(t), the model bypasses the mechanistic derivation of antibody kinetics. This approach cannot capture the complex interplay between viral load and antibody stimulating, preventing the analysis of "kinetic decoupling" between viral replication and immune priming.
4. **Static Constraints on Affinity and Evolution:** The model lacks a realistic representation of affinity selection. In natural infection, high-affinity autoantibodies are selected by the virus. Conversely, for exogenous mAbs, the critical factor is not necessarily *de novo* mutation, but the pre-existence or co-infection of a variant strain against which the specific mAb exhibits reduced affinity ($k_{on}' \ll k_{on}$).

**4.6. Alternative Mechanism: Affinity-Dependent Rebound (Model 3.1.4 Extension)**

We propose that the observed viral rebound is more plausibly explained by the infection of a specific viral variant (Mutant Strain) against which the therapeutic mAb has compromised binding kinetics, rather than rapid *de novo* evolution.

To validate this, we simulated the dynamics of two competing strains: an **Original Strain (V)** and a **Mutant Strain ($V_{mut}$)**. We hypothesize that the mAb binds the Original Strain with high affinity ($kon \approx 10^{-5}$) but exhibits significantly reduced association with the Mutant Strain ($kon' \approx 10^{-6}$), as detailed in **Table S7**.

**Simulation Configuration:**

- **Intervention:** Monoclonal antibody is administered at t=28 (point injection).
- **Mechanism:** The mAb effectively neutralizes the Original Strain. However, due to the lower $k_{on}'$ for the Mutant Strain, a sub-lethal active concentration of mAb creates a selection window.
- **Outcome (Figure S4C):** As the mAb concentration decays metabolically ($k_6$), the

suppression on the Mutant Strain is lifted earlier than on the Original Strain, leading to a distinct rebound peak dominated by the resistant variant. This confirms that **insufficient dosage relative to the variant's binding affinity**, combined with the decay of exogenous antibodies ("kinetic decoupling"), is the primary driver of rebound.

**Table S7 | Parameter sets and initial values for the Multi-Strain Rebound Model**

| Kinetic Parameter | Symbol | Value (Original) | Value (Mutant) | Description |
|---|---|---|---|---|
| **Initial States** | | | | |
| Viral Load | $V_0$ | 1 | 1 | Co-infection starting values |
| Host Antibody | $A_0$ | 10 | 10 | Baseline immunity |
| Exogenous mAb | $A_{2,0}$ | 0 | 0 | Pre-treatment |
| **Therapeutic Kinetics** | | | | |
| **mAb Association** | $k_1'$ | $10^{-5}$ | $10^{-6}$ | **Reduced affinity for mutant** |
| mAb Dissociation | $k_2'$ | $10^{-14}$ | $10^{-14}$ | Complex stability |
| mAb Decay | $k_6$ | 0.05 | 0.05 | Metabolic clearance |
| Dosage | $\pi_{dose}$ | $2\times10^6$ | - | Administered at t=28 |
| **Host Kinetics** | | | | |
| Viral Replication | $k_0$ | 0.5 | 0.5 | |
| Host Ab Association | $k_1$ | $10^{-5}$ | $10^{-5}$ | Assumed cross-reactive |
| Host Ab Feedback | $k_3$ | 1.5 | 1.5 | Endogenous regeneration |

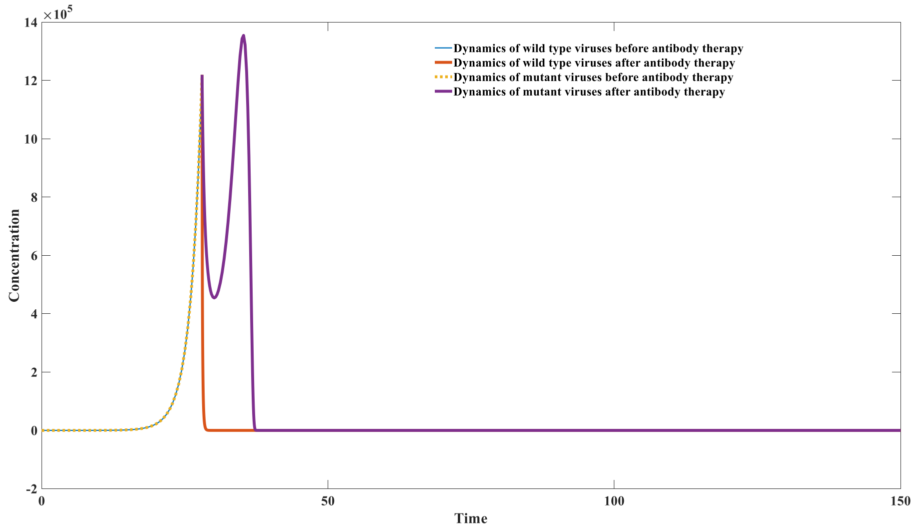


Figure S_4C: Simulation of monoclonal antibody therapy toward wild type virus and mutant strain

#### 5 Mathematical Modeling of Small Molecule Antiviral Therapy (Model 3.1.5)

##### 5.1. Model Structure and Mechanistic Rationale

To investigate the pharmacodynamics of small molecule inhibitors (e.g., cell entry inhibitors, protease inhibitors, or polymerase inhibitors) and their interplay with the host immune response, we developed Model 3.1.5.

Unlike monoclonal antibodies, small molecule drugs ($D$) are modeled to bind free virions ($V$) to form an inhibited complex ($C_2$). A key feature of this model is the consideration of **imperfect inhibition** and **ternary complex formation**:

1. **Residual Replication ($k_0 k_7 C_2$):** The drug-virus complex ($C_2$) may not be completely inert. The parameter $k_7$ represents the fraction of replication potential retained by the inhibited virus (or the rate of release of infectious material).
2. **Immune Synergy ($C_3$):** Host antibodies ($A$) can bind to the virus-drug complex ($C_2$) to form a ternary complex ($C_3$). Crucially, this ternary complex continues to stimulate the immune feedback loop (term $k_3 C_3$ in the $\frac{dA}{dt}$ equation), suggesting that drug-bound virions remain immunogenic.

The system of differential equations is defined as follows:

$$\frac{dV}{dt} = k_0 V - k_1 VA + k_2 C + k_0 k_7 C_2 - p_1 VD + p_2 C_2;$$

$$\frac{dA}{dt} = -k_1 VA + k_2 C + k_3 C - k_1 C_2 A + k_2 C_3 + k_3 C_3 - k_4 A + \pi;$$

$$\frac{dC}{dt} = k_1 VA - k_2 C - k_5 C;$$

$$\frac{dC_2}{dt} = p_1 VD - p_2 C_2 - k_1 C_2 A + k_2 C_3;$$

$$\frac{dC_3}{dt} = k_1 C_2 A - k_2 C_3 - k_5 C_3;$$

$$\frac{dD}{dt} = -p_1 VD + p_2 C_2 - k_6 D + \pi_1(t);$$

Where $\pi_1(t)$ is the dosing function for the small molecule inhibitor.

**5.2. Parameter Statistics and Initial Conditions**

We simulated the viral dynamics under various pharmacologic regimens. The kinetic parameters, including the drug association rate ($p_1$) and metabolic clearance rate ($k_6$), are detailed in **Table S8**.

**Table S8 | Parameter sets and initial values for Model 3.1.5**

| Compartment | Initial Value (t=0) | Parameter | Value | Description |
|---|---|---|---|---|
| $V_0$ | 1 | $k_0$ | 1 | Viral replication rate |
| $A_0$ | 1000 | $k_1$ | $10^{-7}$ | Ab association rate |
| $C_0$ | 0 | $k_2$ | $10^{-14}$ | Ab dissociation rate |
| $C_{2,0}$ | 0 | $k_3$ | 2 | Immune feedback stimulation |
| $C_{3,0}$ | 0 | $k_4$ | 0.01 | Antibody natural decay |
| $D_0$ | 0 | $k_5$ | 0.5 | Complex clearance |
| | | $k_6$ | 0.01 | Drug metabolic clearance |
| | | $k_7$ | 0.1 | **Residual replication factor** (10% leakage) |
| | | $\pi$ | 10 | Basal Ab production |
| | | $p_1$ | $10^{-7}$ | Drug-Virus association rate |
| | | $p_2$ | $10^{-14}$ | Drug-Virus dissociation rate |

##### 5.3. Simulation of Therapeutic Intervention

###### A. Temporal Sensitivity and the "Rebound Window" (Figure S5A)

We simulated the administration of a fixed high dose ($7\times10^8$) of the inhibitor at varying time points (t=16 to t=24). The results uncover a critical "window of vulnerability":

1. **Early Intervention (t<20):** Administration effectively suppresses viral replication early. The viral load is contained without rebound because the combination of drug pressure and basal immunity clears the pathogen.
2. **Critical Rebound Point (t≈20):** Administering the drug at this inflection point triggers a significant **viral rebound**. Mechanistically, the drug temporarily suppresses bacterial load, depriving the immune system of the antigen ($V$) needed to ramp up production ($k_3C$). Once the drug is metabolized ($k_6D$), the insufficient level of antibodies allows the residual virus to resurge.
3. **Late Intervention (t>20):** While late administration fails to lower the peak viral load (aggravating symptoms), it does not cause rebound. By this time, the endogenous immune response has already been primed by the high viral load, rendering the host robust against resurgence.

###### B. Dose-Dependent Dynamics (Figure S5B)

We further analyzed the impact of dosage magnitude (ranging from $1\times10^8$ to $12\times10^8$) administered at the critical time point (t=20). The results exhibit a non-linear response:

- **Sub-therapeutic (<$4\times10^8$):** The drug concentration is insufficient to overcome viral replication (k0$k0$); viral dynamics remain largely unchanged.
- **The "Danger Zone" ($6\times10^8$):** At this intermediate dosage, the viral load experiences a sharp, short-term decline. However, this transient suppression creates a false nadir. The drug concentration is high enough to pause the immune alarm but too low to achieve sterilization. As D$D$ decays, the virus rebounds aggressively to a high peak.
- **Sterilizing Dosage (>$10\times10^8$):** When the dose exceeds this threshold, the inhibition is robust enough to clear the virus independently of the immune lag, permanently suppressing replication and preventing rebound.

Model 3.1.5 demonstrates that small molecule inhibitors, similar to monoclonal antibodies, carry a risk of inducing viral rebound if administered at discontinuous overlapping intervals with the immune response (Kinetic Decoupling). Rebound is most likely when moderate doses are applied during the rapid expansion phase of the virus.

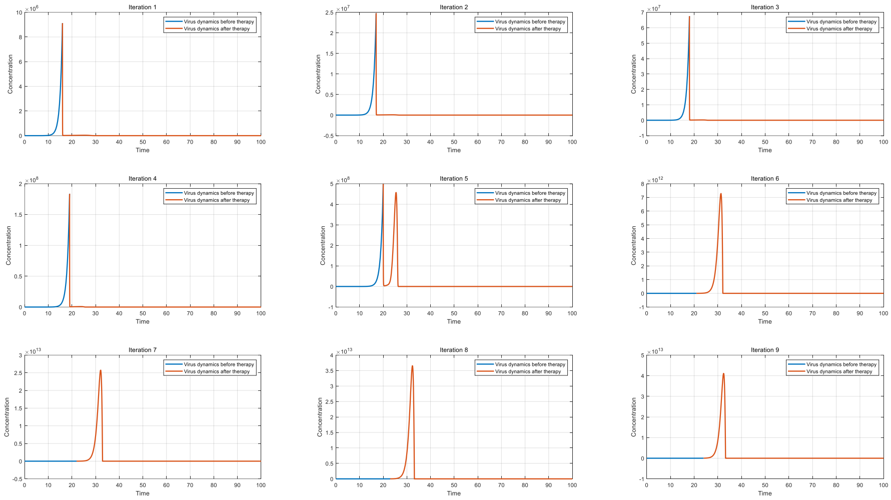


Figure S_5A: Viruses dynamics of different drug injection time

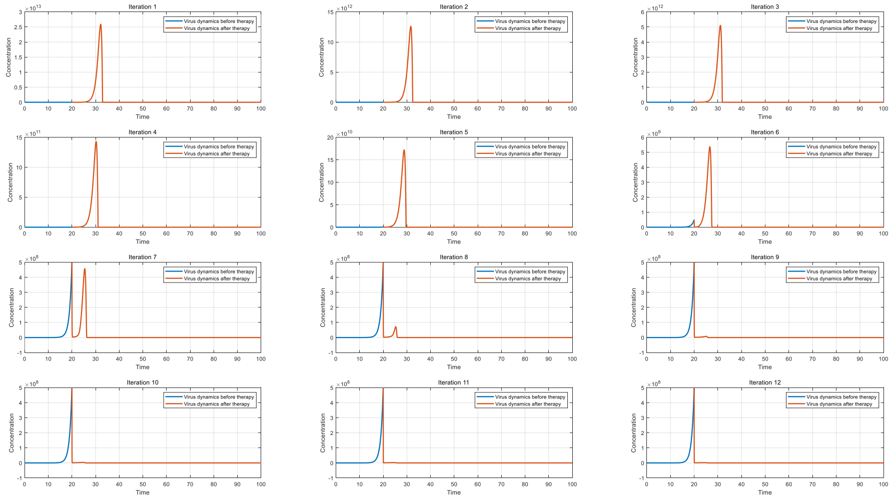


Figure S_5B: Viruses dynamics of different drug injection doses

#### 6 Parameter Settings and Simulation Results for Model 3.1.6

##### 6.1. Model Structure and Governing Equations

The dynamics of the system are described by Model 3.1.6, which governs the interactions between viral/vaccine antigens ($V$), antibodies ($A$), B cells ($B$), plasma cells/ASCs ($P$), and immune complexes ($C_1$, $C_2$). The system of ordinary differential equations (ODEs) is defined as follows:

$$\frac{dV}{dt} = k_0 V - k_1 VA + k_2 C_1 - k_1 VB + k_2 C_2;$$

$$\frac{dA}{dt} = -k_1 VA + k_2 C_1 - k_4 A + k_5 P;$$

$$\frac{dB}{dt} = -k_1VB + k_2C + k_3(1-\alpha)\frac{C_2}{(C_2+B)}C_2 - k_6B;$$

$$\frac{dP}{dt} = k_3\alpha\beta\frac{C_2}{(C_2+B)}C_2 - k_7P;$$

$$\frac{dC_1}{dt} = k_1VA - k_2C_1 - k_8C_1;$$

$$\frac{dC_2}{dt} = k_1VB - k_2C_2 - k_8C_2;$$

**Parameter definitions:**

- **$k_0$ (Antigen replication/decay):** For vaccine simulations, $k_0<0$ denotes non-replicating antigens that degrade over time. For viral infection simulations, $k_0>0$ represents viral replication.
- **α (Differentiation probability):** Represents the fraction of germinal center B cells differentiating into antibody-secreting cells (ASCs).

##### 6.2. Simulation Results

###### 6.2.1 Impact of Vaccination Interval and Dosage on Antibody Dynamics

We first simulated vaccination scenarios ($k_0=-0.02$) to evaluate the effects of discrete dosing strategies.

**Vaccination Interval (Fig. S6A):**
Using a fixed dose of $10^8$ for both primary and booster shots, intervals were varied from 50 to 600 time points.

- **Short Intervals (50 time points):** A transient decline in antibody levels was observed immediately post-boost, followed by recovery.
- **Intermediate Intervals (100–250 time points):** A significant reduction in antibody titers occurred after the second dose, attributed to antibody interference where circulating antibodies neutralize the booster antigen before it can stimulate memory B cells.
- **Long Intervals (>300 time points):** A resurgence in antibody concentration was observed. Notably, intervals exceeding 450 time points yielded peak antibody levels surpassing those of the primary response.
- **Conclusion:** Shorter intervals may accelerate antibody decay dynamics due to interference, suggesting optimal boosting requires sufficient antibody waning.

**Secondary Dosage (Fig. S6B):**
The second dose was administered at t=100, varying from $10^6$ to $10^{10}$ (incremented by $10^{0.5}$).

- **Low Dose:** Minimal impact on antibody dynamics.
- **Intermediate Dose:** Increased doses initially accelerated the decline of circulating antibodies via interference.

- **High Dose:** Threshold effects were overcome, allowing for a robust secondary immune response and net antibody boost.

**Table S9: Parameter sets and initial values for vaccination simulations (Model 3.1.6)**

| Compartment | Initial Value | Parameter | Value |
|---|---|---|---|
| V | $10^8$ | $k_0$ | −0.02 |
| A | 0 | $k_1$ | $10^{-7}$ |
| B | 500 | $k_2$ | $10^{-14}$ |
| P | 0 | $k_3$ | 2 |
| $C_1$ | 0 | $k_4$ | 0.02 |
| $C_2$ | 0 | $k_5$ | $10^5$ |
| | | $k_6$ | 0.005 |
| | | $k_7$ | 0.01 |
| | | $k_8$ | 0.5 |
| | | α | 0.5 |
| | | β | $10^{-3}$ |

##### 6.2.2 Influence of B-cell Differentiation (α) on Viral Control

We investigated the parameter α (0≤α≤1), which dictates the balance between B-cell proliferation and differentiation into ASCs.

- **Low *α*:** Impairs the conversion of non-plasma B cells to ASCs, limiting antibody production and facilitating viral replication.
- **High *α*:** Excessive differentiation depletes the pool of proliferating non-plasma B cells, ultimately capping the potential number of ASCs generated.

**Viral Dynamics (Fig. S6C):**
Simulations with α ranging from 0.05 to 0.45 (Table S10) reveal a non-monotonic relationship: as α increases, peak viral concentration decreases to a minimum before rising again.

**Optimal Parameter Space (Fig. S6D):**
The optimal α for viral suppression is a function of viral replication ($k_0$) and antibody regeneration ($k_3$):

1. **High Immunity ($k_3$↑)/Low Pathogenicity ($k_0$↓):** A higher conversion rate (high α) is optimal. Robust germinal centers facilitate rapid ASC conversion to clear infection.
2. **Weak Immunity/High Pathogenicity ($k_0$↑):** A lower conversion rate is beneficial to maintain the B-cell progenitor pool.

**Table S10: Parameter sets and initial values for α variation simulations**

| Compartment | Initial Value | Parameter | Value |
|---|---|---|---|
| V | 1 | $k_0$ | 0.5 |
| A | 0 | $k_1$ | $10^{-7}$ |
| B | 500 | $k_2$ | $10^{-14}$ |
| P | 0 | $k_3$ | 2 |
| $C_1$ | 0 | $k_4$ | 0.02 |
| $C_2$ | 0 | $k_5$ | $10^5$ |
| | | $k_6$ | 0.005 |
| | | $k_7$ | 0.01 |
| | | $k_8$ | 0.5 |
| | | $\beta$ | $10^{-3}$ |

##### 6.2.3 Antibody Exhaustion in Severe Infection

Consistent with Model 3.1.1, Model 3.1.6 predicts scenarios of antibody exhaustion in severe patients (Fig. S6E). This occurs when the rate of antibody depletion exceeds regeneration capacity, typically driven by high viral replication ($k_0$) and low antibody regeneration coefficients ($k_3$).

- **Mechanism:** Equilibrium analysis indicates that both disease-free and endemic equilibria become unstable.
- **Simulation:** Under conditions of strong viral replication ($k_0$=0.55) and moderate regeneration ($k_3$=2, $\alpha$=0.5), viral load increases indefinitely while antibody concentrations remain suppressed (Table S11). This signifies a loss of germinal center activity and functional immune collapse.

**Table S11: Parameter sets for Antibody Exhaustion scenario**

| Compartment | Initial Value | Parameter | Value |
|---|---|---|---|
| V | 1 | $k_0$ | 0.55 |
| A | 0 | $k_1$ | $10^{-7}$ |
| B | 500 | $k_2$ | $10^{-14}$ |
| P | 0 | $k_3$ | 2 |
| $C_1$ | 0 | $k_4$ | 0.02 |
| $C_2$ | 0 | $k_5$ | $10^5$ |
| | | $k_6$ | 0.005 |
| | | $k_7$ | 0.01 |
| | | $k_8$ | 0.5 |
| | | $\beta$ | $10^{-3}$ |

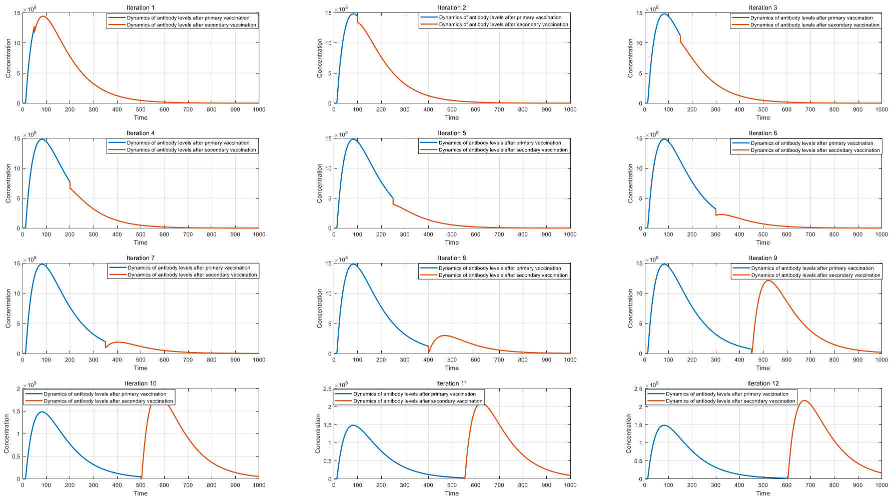


**Figure S6A:** Biphasic antibody dynamics resulting from varying vaccination intervals.

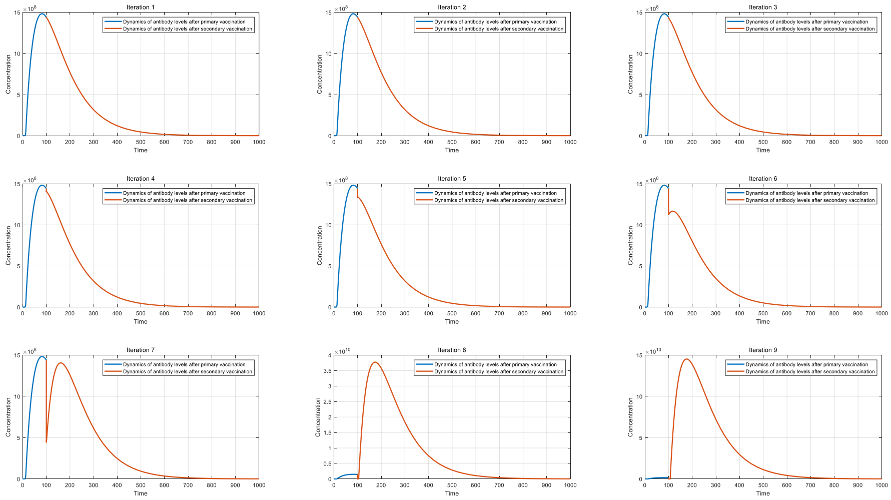


**Figure S6B:** Dose-dependent antibody kinetics and interference effects during secondary vaccination.

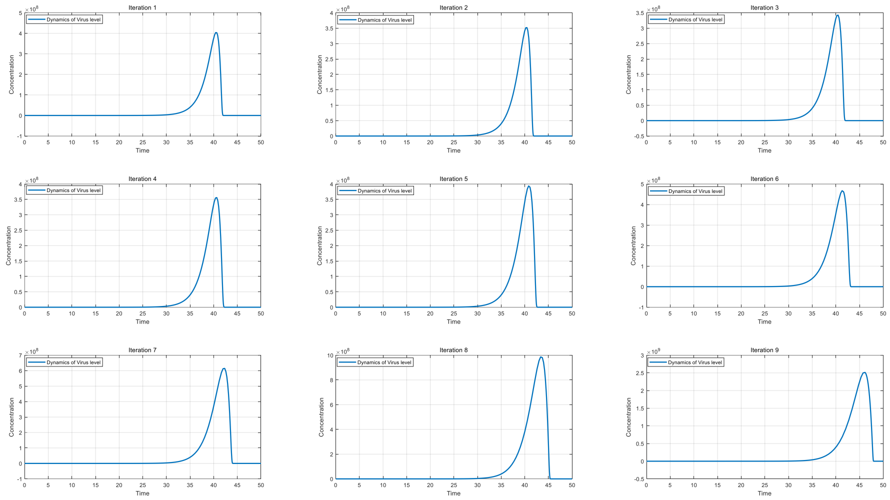


**Figure S6C:** Effect of differentiation parameter α on viral load trajectories.

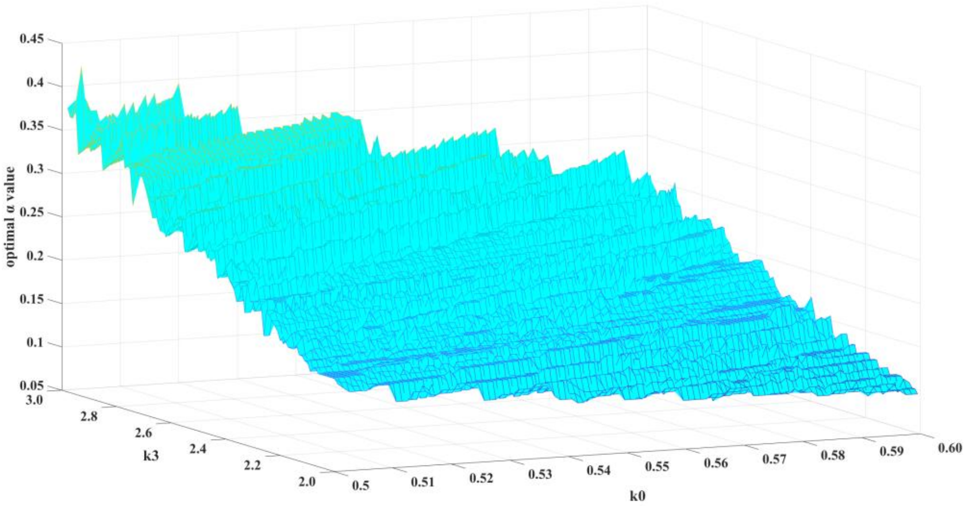


**Figure S6D:** Heatmap distribution of optimal α values across varying $k_0$ and $k_3$ combinations.

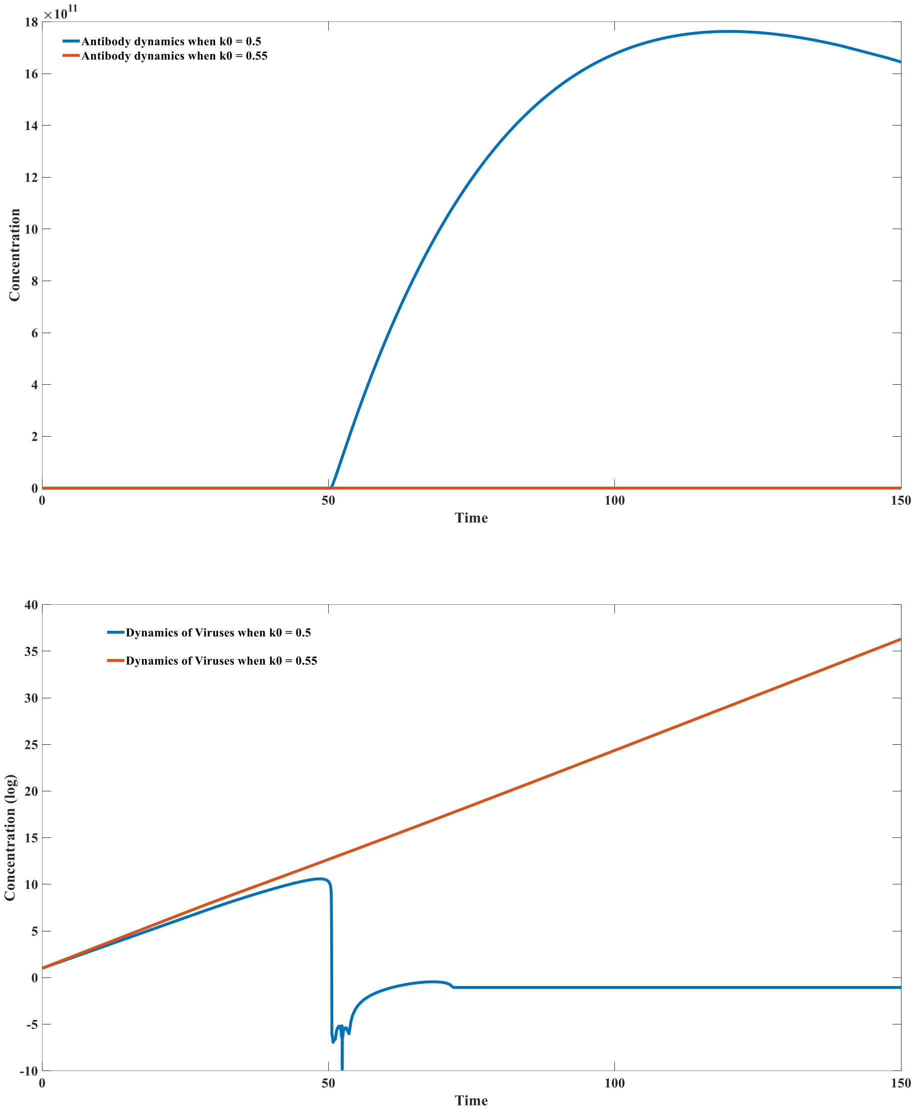


**Figure S6E:** Temporal dynamics of antibody exhaustion in response to high-replication viral infection.

#### 7 Parameter Settings and Simulation Results for Model 3.1.7

##### 7.1. Model Structure and Governing Equations

Model 3.1.7 extends the previous framework to include isotype switching (IgM to IgG), B-cell receptor (BCR) dynamics, antibody-secreting cell (ASC) differentiation, and interactions with both viral (V) and environmental (E) antigens. The dynamics for the $i$-th antibody clone are governed by the following system of ordinary differential equations:

**B-cell Receptor (BCR) Dynamics:**

$$\frac{dB_i^{IgM}}{dt} = \underbrace{\pi f_i}_{\text{Influx}} - p_1 E B_i^{IgM} + p_2 C_{2,i} + \underbrace{k_3 C_{2,i}}_{\text{Env-driven Prolif.}} - k_{1,i} V B_i^{IgM} + k_{2,i} C_{1,i}$$

$$+ \underbrace{k_4(1-\gamma)(1-\theta)\frac{C_{1,i}}{C_{1,i} + B_i^{IgM}} C_{1,i}}_{\text{Virus-driven Prolif.\& Retention}} - k_5 B_i^{IgM};$$

$$\frac{dB_i^{IgG}}{dt} = -p_1 E B_i^{IgG} + p_2 C_{4,i} + \underbrace{k_3 C_{4,i}}_{\text{Env-driven Prolif.}} - k_{1,i} V B_i^{IgG} + k'_{2i} C_{3i}$$

$$+ \underbrace{k_4 \gamma(1-\theta)\frac{C_{1,i}}{C_{1i} + B_i^{IgG}} C_{1,i}}_{\text{Class Switching (IgM}\rightarrow\text{IgG)}} + \underbrace{k_4(1-\theta)\frac{C_{3,i}}{C_{3,i} + B_i^{IgG}} C_{3,i}}_{\text{Self-Proliferation}} - k_6 B_i^{IgG};$$

**Secreted Antibody Dynamics:**

$$\frac{\mathrm{d}A_i^{IgM}}{\mathrm{dt}} = \mathrm{k}_7 P_i^{IgM} - \mathrm{p}_1 \mathrm{E} A_i^{IgM} + \mathrm{p}_2 \mathrm{C}_{6,\mathrm{i}} - \mathrm{k}_{1,\mathrm{i}} \mathrm{V} A_i^{IgM} + \mathrm{k}_{2,\mathrm{i}} \mathrm{C}_{5,\mathrm{i}} - \mathrm{k}_8 A_i^{IgM};$$

$$\frac{\mathrm{d}A_i^{IgG}}{\mathrm{dt}} = \mathrm{k}_9 P_i^{IgG} - \mathrm{p}_1 \mathrm{E} A_i^{IgG} + \mathrm{p}_2 \mathrm{C}_{8,\mathrm{i}} - \mathrm{k'}_{1,\mathrm{i}} \mathrm{V} A_i^{IgG} + \mathrm{k'}_{2,\mathrm{i}} \mathrm{C}_{7,\mathrm{i}} - \mathrm{k}_{10} A_i^{IgG};$$

**Antibody-Secreting Cell (ASC) Dynamics:**

$$\frac{dP_i^{IgM}}{dt} = \underbrace{k_{12} C_{2,i}}_{\text{Basal Diff.}} + \underbrace{k_4 \epsilon (1-\gamma)\theta \frac{C_{1,i}}{C_{1,i} + B_i^{IgM}} C_{1,i}}_{\text{Virus-driven Differentiation}} - k_{13} P_i^{IgM};$$

$$\frac{\mathrm{d}P_i^{IgG}}{\mathrm{dt}} = \mathrm{k}_{12} \mathrm{C}_{4,\mathrm{i}} + \mathrm{k}_4 \epsilon \theta \frac{\mathrm{C}_{3,\mathrm{i}}}{\mathrm{C}_{3,\mathrm{i}} + B_i^{IgG}} \mathrm{C}_{3,\mathrm{i}} - \mathrm{k}_{14} P_i^{IgG};$$

**Immune Complexes Kinetics:**

The model tracks eight distinct species of immune complexes ($C_{1,i}$...$C_{8,i}$), subject to a uniform clearance rate $k_{11}$.

*Note: For brevity, representative equations for viral (V) and environmental (E) complexes are shown below.*

$$\frac{dC_{1,i}}{dt} = k_{1,i} V B_i^{IgM} - k_{2,i} C_{1,i} - k_{11} C_{1,i};$$

$$\frac{dC_{3,i}}{dt} = k'_{1,i} V B_i^{IgG} - k'_{2,i} C_{3,i} - k_{11} C_{3,i};$$

$$\frac{dC_{5,i}}{dt} = k_{1,i} V A_i^{IgM} - k_{2,i} C_{5,i} - k_{11} C_{5,i};$$

$$\frac{dC_{7,i}}{dt} = k'_{1,i} V A_i^{IgG} - k'_{2,i} C_{7,i} - k_{11} C_{7,i};$$

Complexes involving environmental antigens ($C_2$, $C_4$, $C_6$, $C_8$) follow an analogous structure determined by parameters $p_1$ (association) and $p_2$ (dissociation).

$$\frac{dC_{2,i}}{dt} = \mathrm{p}_1 E B_i^{IgM} - \mathrm{p}_2 C_{2,i} - k_{11} C_{2,i};$$

$$\frac{dC_{4,i}}{dt} = \mathrm{p}_1 E B_i^{IgG} - \mathrm{p}_2 C_{4,i} - k_{11} C_{4,i};$$

$$\frac{dC_{6,i}}{dt} = \mathrm{p}_1 E A_i^{IgM} - \mathrm{p}_2 C_{6,i} - k_{11} C_{6,i};$$

$$\frac{dC_{8,i}}{dt} = \mathrm{p}_1 E A_i^{IgG} - \mathrm{p}_2 C_{8,i} - k_{11} C_{8,i};$$

**Antigen and Global Dynamics**

$$\frac{dV}{dt} = k_0 V - \sum_n^{i=1} k_{1,i} V B_i^{IgM} + \sum_n^{i=1} k_{2,i} C_{1,i} - \sum_n^{i=1} k'_{1,i} V B_i^{IgG} + \sum_n^{i=1} k'_{2,i} C_{3,i} - \sum_n^{i=1} k_{1,i} V A_i^{IgM} + \sum_n^{i=1} k_{2,i} C_{5,i} - \sum_n^{i=1} k'_{1,i} V A_i^{IgG} + \sum_n^{i=1} k'_{2,i} C_{7,i};$$

$$\frac{dE}{dt} = \pi_1 - \sum_n^{i=1} p_1 E B_i^{IgM} + \sum_n^{i=1} p_2 C_{2,i} - \sum_n^{i=1} p_1 E B_i^{IgG} + \sum_n^{i=1} p_2 C_{4,i} - \sum_n^{i=1} p_1 E A_i^{IgM} + \sum_n^{i=1} p_2 C_{6,i} - \sum_n^{i=1} p_1 E A_i^{IgG} + \sum_n^{i=1} p_2 C_{8,i};$$

#### 7.2. Parameterization Strategy

##### 7.2.1 General Parameters

Parameters essentially independent of antibody isotype are listed in Table S12.

**Table S12: Parameter sets and initial values in Model 3.1.7**

| Name | Meaning | Value |
|---|---|---|
| $k_0$ | Virus replication constant | 1.2 |
| $k_3$ | Antibody regeneration constant (Environmental Ag complex) | 1 |
| $k_4$ | Antibody regeneration constant (Virus complex) | 2 |
| $k_5$ | Decay constant of IgM-BCR B cell | 0.01 |
| $k_6$ | Decay constant of IgG-BCR B cell (Memory B cell) | 0.005 |
| $k_7$ | IgM production rate of IgM-ASC | $4.4\times10^9$ |
| $k_8$ | Decay constant of IgM | 0.05 |
| $k_9$ | IgG production rate of IgG-ASC | $1.2\times10^{10}$ |
| $k_{10}$ | Decay constant of IgG | 0.025 |
| $k_{11}$ | Decay constant of antibody-antigen complex | 0.5 |
| $k_{12}$ | Regeneration constant of ASC from Environmental Ag complex | $5\times10^{-7}$ |
| $k_{13}$ | Decay constant of IgM-secreting plasma B cell | 0.1 |
| $k_{14}$ | Decay constant of IgG-secreting plasma B cell (Memory plasma) | 0.05 |
| $\gamma$ | Transformation ratio of IgG-BCR from IgM-BCR (Class switching) | 0.02 |
| $\theta$ | Maximal transformation ratio of ASC from non-plasma B cell | 0.1 |
| $\epsilon$ | ASC/BCR ratio | $10^5$ |
| $p_1$ | Forward binding constant (Ab + Environmental Ag) | $10^{-20}$ |
| $p_2$ | Dissociation constant (Ab-Environmental Ag complex) | 0.5 |
| $\pi$ | Overall supplement of non-plasma IgM-BCR B cell | $0.5\times10^{13}$ |
| $\pi_1$ | Supplement of environmental antigens | $2.2\times10^{17}+10^{13}$ |

##### 2.2 Disease-Free Equilibrium (DFE) Calibration

Given the high dimensionality of the parameter space, we adopted a calibration strategy based on the disease-free equilibrium (DFE) state. In the absence of viral infection (V=0), the viral-specific complexes ($C_1$, $C_3$, $C_5$, $C_7$) vanish, reducing the system to the following steady-state equations:

$$0 = \underbrace{\pi f_i}_{\text{Influx}} - p_1 E B_i^{IgM} + p_2 C_{2,i} + \underbrace{k_3 C_{2,i}}_{\text{Env-driven Prolif.}} - k_5 B_i^{IgM}$$

$$0 = -p_1 E B_i^{IgG} + p_2 C_{4,i} + \underbrace{k_3 C_{4,i}}_{\text{Env-driven Prolif.}} - k_6 B_i^{IgG}$$

$$0 = \mathrm{k}_7 P_i^{IgM} - \mathrm{p}_1 \mathrm{E} A_i^{IgM} + \mathrm{p}_2 \mathrm{C}_{6,\mathrm{i}} - \mathrm{k}_8 A_i^{IgM}$$
$$0 = \mathrm{k}_9 P_i^{IgG} - \mathrm{p}_1 \mathrm{E} A_i^{IgG} + \mathrm{p}_2 \mathrm{C}_{8,\mathrm{i}} - \mathrm{k}_{10} A_i^{IgG}$$
$$0 = \underbrace{k_{12} C_{2,i}}_{\text{Basal Diff.}} - k_{13} P_i^{IgM}$$
$$0 = \mathrm{k}_{12} \mathrm{C}_{4,\mathrm{i}} - \mathrm{k}_{14} P_i^{IgG}$$
$$0 = \mathrm{p}_1 E B_i^{IgM} - \mathrm{p}_2 C_{2,i} - k_{11} C_{2,i}$$
$$0 = \mathrm{p}_1 E B_i^{IgG} - \mathrm{p}_2 C_{4,i} - k_{11} C_{4,i}$$
$$0 = \mathrm{p}_1 E A_i^{IgM} - \mathrm{p}_2 C_{6,i} - k_{11} C_{6,i}$$
$$0 = \mathrm{p}_1 E A_i^{IgG} - \mathrm{p}_2 C_{8,i} - k_{11} C_{8,i}$$

$$0 = \pi_1 - \sum_{n}^{i=1} p_1 E B_i^{IgM} + \sum_{n}^{i=1} p_2 C_{2,i} - \sum_{n}^{i=1} p_1 E B_i^{IgG} + \sum_{n}^{i=1} p_2 C_{4,i} - \sum_{n}^{i=1} p_1 E A_i^{IgM} + \sum_{n}^{i=1} p_2 C_{6,i}$$
$$- \sum_{n}^{i=1} p_1 E A_i^{IgG} + \sum_{n}^{i=1} p_2 C_{8,i}$$

This reduced system consists of 11 equations governing 11 equilibrium state variables, involving 15 parameters ($k_3$, $k_5$, …, $k_{14}$, $p_1$, $p_2$, $\pi$, $\pi_1$).

To solve this system, we estimated initial values and subsets of parameters based on existing literature and biological plausibility. We fixed specific priors (constants derived from previous models) and solved the algebraic system to determine the remaining parameter values. The results of this calibration are presented in Table S13.

**Table S13: Known parameters and initial values for Model 3.1.7 (DFE State)**

| Parameter/State | Value |
|---|---|
| IgM-BCR | $10^{15}$ |
| IgG-BCR | $10^{15}$ |
| IgM | $4\times10^{18}$ |
| IgG | $4\times10^{19}$ |
| $C_{2,i}$ | $10^{13}$ |
| $C_{4,i}$ | $10^{13}$ |

| Parameter/State | Value |
|---|---|
| $C_{6,i}$ | $4\times10^{16}$ |
| $C_{8,i}$ | $4\times10^{17}$ |
| IgM-ASC | $5\times10^{7}$ |
| IgG-ASC | $10^{8}$ |
| E | $10^{18}$ |
| $k_3$ | 1 |
| $k_5$ | 0.01 |
| $k_6$ | 0.005 |
| $k_8$ | 0.05 |
| $k_{10}$ | 0.025 |
| $k_{11}$ | 0.5 |
| $k_{13}$ | 0.1 |
| $k_{14}$ | 0.005 |
| $p_1$ | $10^{-20}$ |
| $p_2$ | 0.5 |

##### 7.3. Antibody Clonal Diversity and Kinetic Parameters

To capture the heterogeneity of the immune response, we modeled a polyclonal repertoire consisting of 100 distinct antibody clones for each isotype (IgM and IgG) and their corresponding cellular compartments (BCR, ASC). Each antibody clone $i$ is uniquely defined by a pair of integer indices $(m,n)$, where $1\leq m,n\leq 10$ such that the clone index is given by $i=10(m-1)+n$.

**Kinetic Rate Constants**

The kinetic properties of each clone are determined by the indices $m$ (governing association) and $n$ (governing dissociation).

- **Dissociation Rate ($k_{off}$):** The dissociation rate constant $k_{2,i}$ is identical for both isotypes (assuming the same epitope-paratope bond stability) and is defined as:
  $k_{2,i}=k_{2,i}'=10^{(n-1)}$
- **Association Rate ($k_{on}$):** The association rate constants vary by isotype to account for valency differences.
  - **IgG:** Modeled as a monomeric interaction.
    $k_{1,i}=10^{(m-23)}$
  - **IgM:** To account for the high avidity of IgM, which exists as a pentamer in serum and multimeric complexes on B cells, we applied a scaling factor. The effective association rate is set to five times that of IgG:
    $k_{1,i}=5\times10^{(m-23)}$

**Initial Clonal Distribution**

The initial abundance of each clone follows a log-normal distribution, derived from experimental reports on affinity landscapes. The probability mass function $F_i$ for the $i$-th clone is calculated as the product of the probabilities for the kinetic coordinates $m$ and $n$:

$$F_i = P(m-1 \leq X \leq m) \times P(n-1 \leq Y \leq n)$$

Here, variables X (association) and Y (dissociation) follow a normal distribution $N(\mu=5, \sigma=0.82)$.

**Initialization**

The initial populations for all compartments—$B_i^{IgM}$, $B_i^{IgG}$, $A_i^{IgM}$, $A_i^{IgG}$, $P_i^{IgM}$ and $P_i^{IgG}$—were determined by distributing the total initial population sizes (e.g., 100 cells/molecules total per category) across the 100 clones according to the probability weighting $F_i$.

##### 7.4. Simulation Results and Biological Interpretation

###### 7.4.1 Clonal Dynamics and Stability (Fig. S7A)

Simulation of the antibody repertoire following viral infection reveals distinct kinetic profiles for IgM and IgG isotypes (Fig. S7A):

- **IgM Dynamics:** Driven by the multimeric nature of IgM and IgM-BCRs, the system exhibits a rapid initial proliferation of IgM. The enhanced avidity of multimers directs the immune response toward immediate containment. However, IgM levels decline rapidly post-infection and fail to sustain a stable peak in the high-affinity region over extended periods.
- **IgG Dynamics:** In contrast, IgG proliferation is slower but results in the sustained dominance of high-affinity clones.

**Stability Analysis:** Linear stability analysis of the post-infection state confirms that the new disease-free equilibrium is stable (all eigenvalues are negative). This stability provides the dynamical framework for immunological memory, ensuring that high-affinity IgG clones persist long-term.

###### 7.4.2 Evolutionary Implications of Isotype Structure

The simulation results suggest that the structural differences between isotypes—multimeric IgM versus monomeric IgG—represent an evolutionary optimization between response speed and metabolic efficiency:

1. **Rapid Activation (IgM):** The multimeric structure of IgM-BCRs compensates for low initial affinity through high avidity, allowing for rapid B-cell activation and immediate pathogen suppression before affinity maturation occurs.
2. **Efficient Memory (IgG):** IgG predominates as a monomer. Since specific binding affinity increases during maturation, the host can afford to switch to a monomeric form. This is metabolically advantageous; for an equivalent synthesis mass, monomeric IgG yields a higher molar concentration than pentameric IgM, maximizing viral neutralization capacity per unit of metabolic investment during the memory phase.

##### 7.4.3 *In Silico* Serology (Fig. S7B)

To validate the model against standard clinical diagnostics, we simulated an Enzyme-Linked Immunosorbent Assay (ELISA).

- **Methodology:** The full Model 3.1.7 was reduced to a cell-free system (removing BCR and ASC compartments) with no viral replication ($k_0$=0).
- **Antigen Dosage:** The simulation utilized a fixed antigen dosage of $10^{16}$. This parameter is critical; insufficient antigen dosage causes non-specific binding of moderate-affinity antibodies to dominate the signal, while excessive dosage creates a "hook effect" where un-neutralized antigen obscures results.
- **Results:** As illustrated in **Figure S7B**, the simulation reproduces canonical serological patterns: IgM levels surge significantly during primary infection, whereas IgG dominates the response during secondary infection. This confirms that the model accurately captures the isotype switching and memory kinetics observed in biological systems.

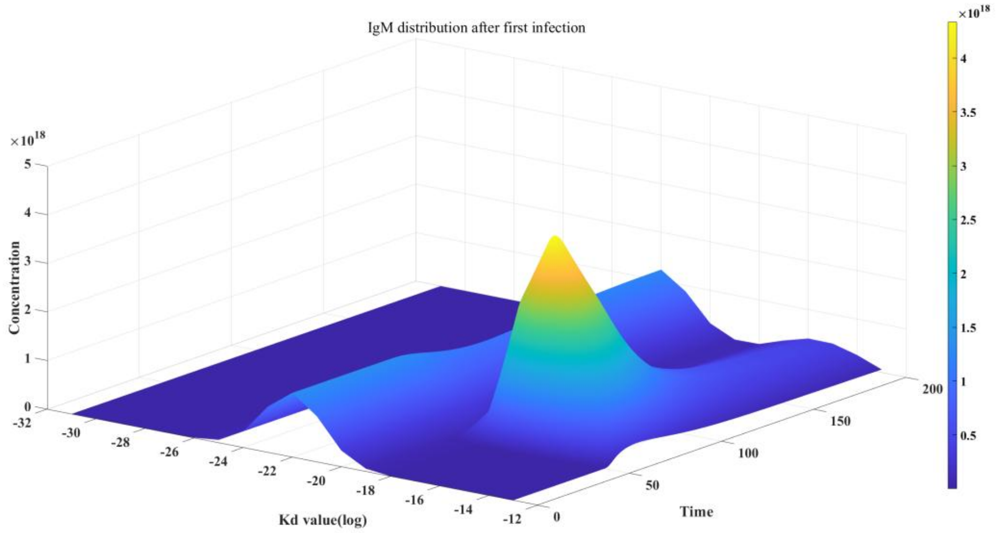

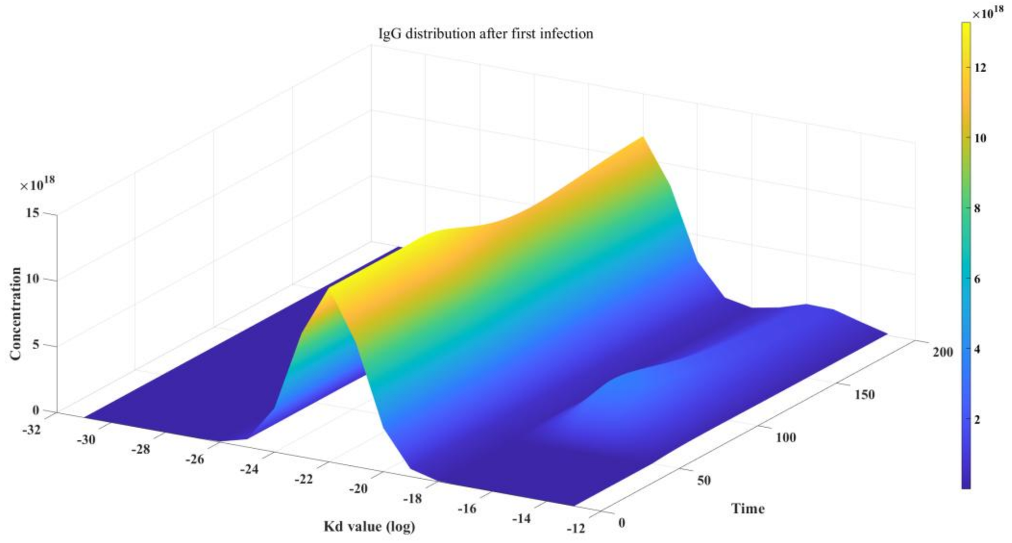


**Figure S7A: Temporal distribution of antibody affinity maturation.** The heatmap visualizes the evolution of the antibody repertoire ($m \times n$ landscape) post-infection. IgM exhibits a rapid, broad-spectrum response that fades quickly, whereas IgG matures into a focused, high-affinity population that persists over time.

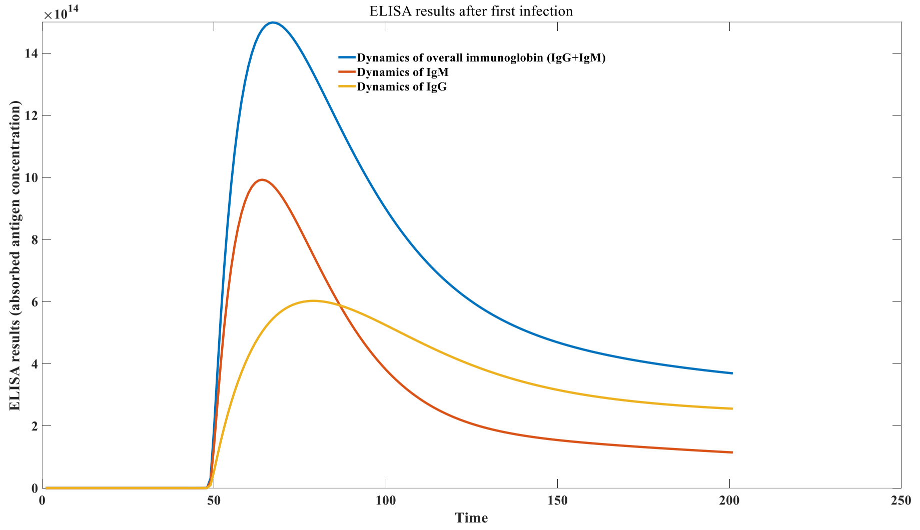

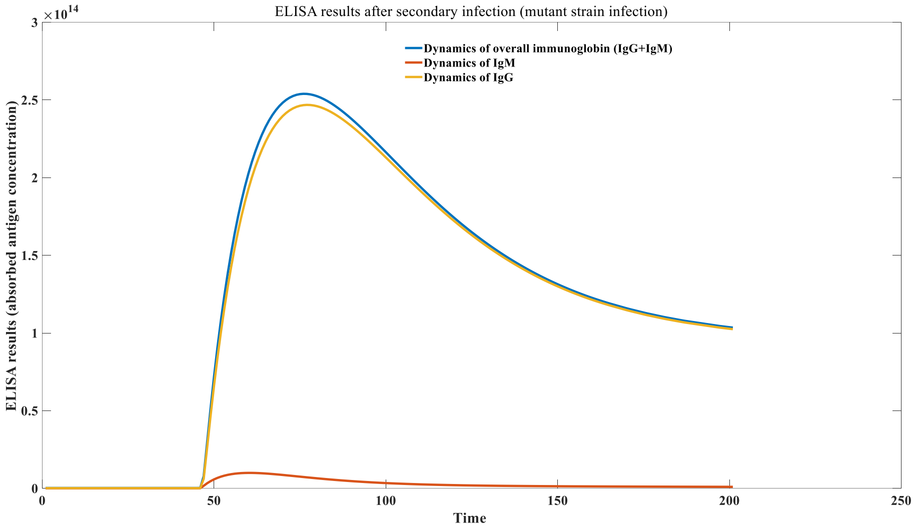


**Figure S7B:** Simulated ELISA kinetics. Comparison of IgM and IgG titers during primary and secondary infections, demonstrating the classic isotype switch and the robust memory recall response of IgG.

#### 7.5 Extended Model for Immune Memory and Discontinuous Antibody Decay (Model 3.1.8)

##### 7.5.1. Rationale and Model Expansion

To investigate the mechanisms underlying the persistence of immune memory and the phenomenon of discontinuous antibody decay, we expanded Model 3.1.7 to distinguish the developmental origins of memory B cells. In this extended framework (Model 3.1.8), the IgG-BCR compartment is stratified into two distinct populations:

1. $B_{i,1}^{IgG}$ Memory B cells maintained through the self-renewal/proliferation of existing IgG-BCR clones.
2. $B_{i,2}^{IgG}$: Memory B cells newly generated via isotype switching from the IgM-BCR compartment.

This distinction allows for the simulation of competition between established memory clones and newly generated clones arising from heterologous infections.

##### 7.5.2. Governing Equations

**B-cell Receptor (BCR) Dynamics:**

The dynamics for IgM-BCR remain similar to previous models, while the IgG-BCR equation is split. The term γ governs the class-switching flow from IgM to the $B_{i,2}^{IgG}$ compartment.

$$\frac{dB_i^{IgM}}{dt} = \underbrace{\pi f_i}_{\text{Influx}} - p_1 E B_i^{IgM} + p_2 C_{2,i} + \underbrace{k_3 C_{2,i}}_{\text{Env-driven Prolif.}} - k_{1,i} V B_i^{IgM} + k_{2,i} C_{1,i}$$

$$+ \underbrace{k_4(1-\gamma)(1-\theta)\frac{C_{1,i}}{C_{1,i} + B_i^{IgM}} C_{1,i}}_{\text{Virus-driven Prolif.\& Retention}} - k_5 B_i^{IgM};$$

$$\frac{dB_{i,1}^{IgG}}{dt} = -p_1 E B_{i,1}^{IgG} + p_2 C_{4,i} + \underbrace{k_3 C_{4,i}}_{\text{Env-driven Prolif.}} - k_{1,i} V B_{i,1}^{IgG} + k'_{2,i} C_{3i}$$

$$+ \underbrace{k_4(1-\theta)\frac{C_{3,i}}{C_{3,i} + B_{i,1}^{IgG} + B_{i,2}^{IgG}} C_{3,i}}_{\text{Self-Proliferation}} - k_6 B_{i,1}^{IgG};$$

$$\frac{dB_{i,2}^{IgG}}{dt} = -p_1 E B_{i,2}^{IgG} + p_2 C_{4,i}' + \underbrace{k_3 C_{4,i}'}_{\text{Env-driven Prolif.}} - k_{1,i} V B_{i,2}^{IgG} + k'_{2,i} C_{3i}'$$

$$+ \underbrace{k_4 \gamma (1-\theta)\frac{C_{1,i}}{C_{1i} + B_i^{IgM}} C_{1,i}}_{\text{Class Switching (IgM}\rightarrow\text{IgG)}} + \underbrace{k_4(1-\theta)\frac{C_{3,i}'}{C_{3,i}' + B_{i,1}^{IgG} + B_{i,2}^{IgG}} C_{3,i}}_{\text{Self-Proliferation}} - k_6 B_{i,2}^{IgG};$$

Note that $C_{3i}'$ and $C_{4i}'$ represent complexes formed by the newly switched $B_{i,2}^{IgG}$ population.

##### 7.5.3. Simulation Results: The Mechanism of Discontinuous Decay

**Origins of Memory B Cells (Figure S7C)**

We simulated the viral infection course to track the quantitative evolution of $B_{i,1}^{IgG}$ and $B_{i,2}^{IgG}$. The results, presented in Figure S7C, demonstrate that a significant proportion of the post-infection IgG-BCR pool consists of type 2 cells ($B_{i,2}^{IgG}$)—high-affinity clones derived directly from the conversion of IgM-BCRs during the active immune response.

**Theory of Discontinuous Antibody Decay**

These results support a mechanistic theory for the stepwise decline of specific antibodies observed clinically:

1. **Homeostasis via Environmental Antigen:** The total size of the antibody/BCR pool is constrained by the availability of environmental antigens (E), which provide necessary survival signals.
2. **Dilution by Heterologous Challenge:**
   - Ideally, if a host encounters no further pathogens, the high-affinity IgG produced against the initial virus would persist indefinitely.
   - However, in reality, hosts are sequentially challenged by heterologous viruses.
   - Upon infection with a *new* virus, the IgM-BCR pool (which has reverted to a generic, naive distribution post-infection) activates and undergoes class switching.
   - This generates a new influx of $B_{i,2}^{IgG}$ specific to the *new* virus.

3. **Stepwise Decline:** Because the IgM-BCR pool resets to a naive state between infections, the new $B_{i,2}^{IgG}$ cells generated during a heterologous infection have low affinity for the *original* virus. These new cells enter the memory compartment and compete for limited survival signals (E). This influx dilutes the population of memory cells specific to the original virus.
4. **Conclusion:** Consequently, antibody titers against a specific pathogen do not decay continuously solely due to half-life. Instead, they exhibit a "discontinuous stepwise decay," dropping significantly with every subsequent distinct viral infection. This downward trend is only reversed (rejuvenated) if the host encounters the original virus again or a closely related variant.

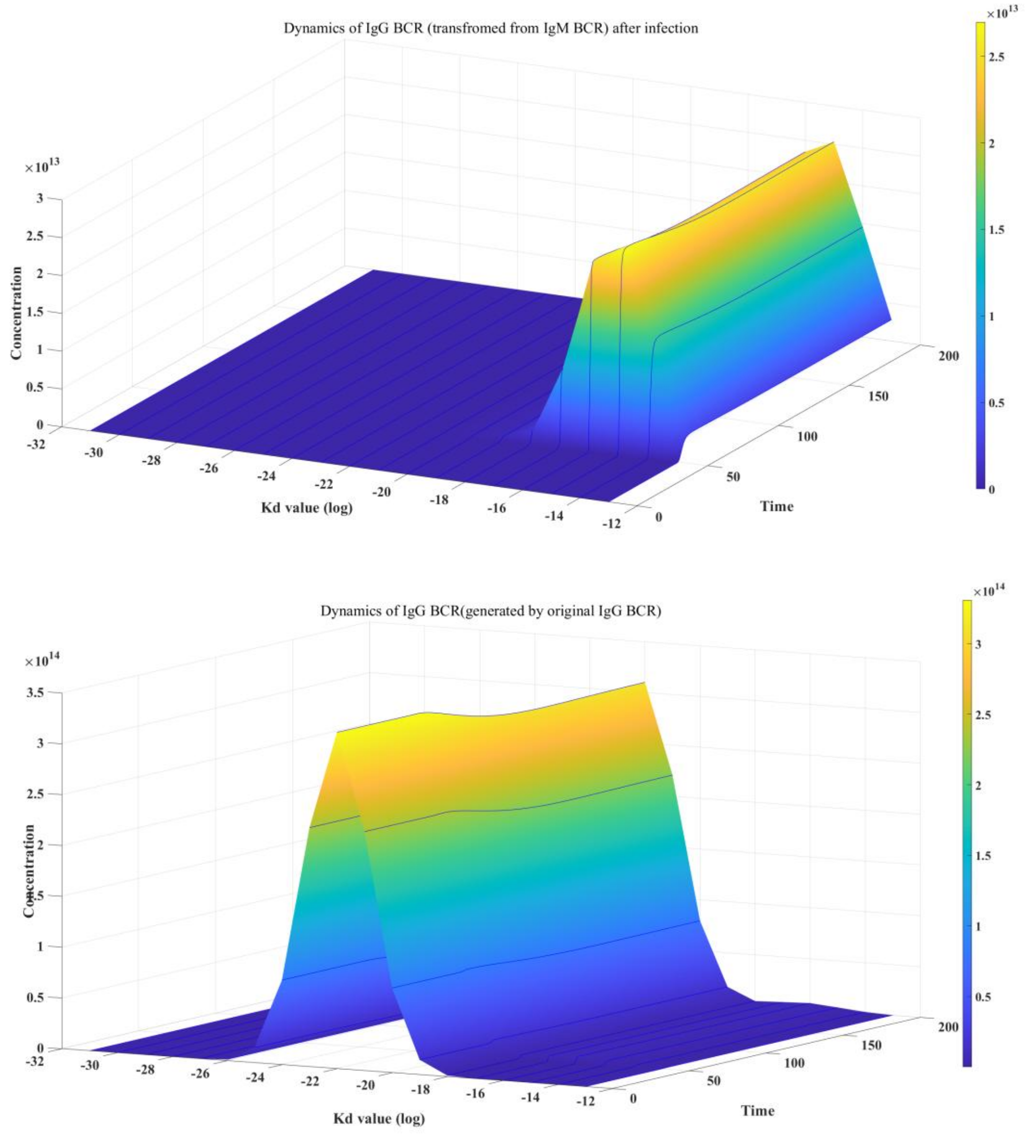


Figure S7_C: Dynamics of IgG with different originality

#### 8. Numerical Simulation of Immune Imprinting Principles

To investigate the mechanisms of immune imprinting (often referred to as "original antigenic sin"), it is necessary to model the dynamics of secondary infections with antigenically distinct variants.

##### 8.1 Modeling Antigenic Drift

The immune landscape differs significantly between stable DNA viruses and rapidly mutating RNA viruses. For DNA viruses, while antibody titers exhibit a stepwise discontinuous decay (as established in Model 3.1.8), the decline is relatively slow, often providing lifelong protection. Conversely, for RNA viruses, high mutation rates lead to a mismatch in affinity distributions. An antibody repertoire that forms a high-affinity peak against the wild-type (WT) strain will perceive a mutant strain differently; the effective affinity distribution against the mutant lies between the mature "imprinted" distribution and a naive normal distribution.

To quantify this, we introduced a mutation coefficient, α. The effective probability distribution of the antibody binding parameters against a mutant strain, $F_{New}$, is modeled as a linear combination of the distribution against the wild-type ($F$) and the naive reference distribution ($f$):

$$F_{New} = (1 - 10^{-\alpha})f + 10^{-\alpha}F$$

Where:

- $f(x;\mu,\sigma)$ is the initial naive normal distribution ($N(\mu,\sigma)$).
- $F$ is the distribution of antibody binding parameters generated against the wild-type antigen.
- $\alpha$ is the mutation coefficient. When $\alpha$=0, $F_{New} = F$ (identity to WT).
  As $\alpha \to \infty$, $F_{New} \to f$ (complete escape/novelty).

##### 8.2 Viral Dynamics in Secondary Infection (Fig. S8A)

We simulated the relationship between viral proliferation and antigenic drift ($\alpha$) during a secondary challenge, assuming the host had reached a steady antibody state from a primary infection.

**Results:** As illustrated in **Figure S8A**, a threshold effect is observed. For low mutation coefficients (α≤3), the existing antibody pool prevents viral proliferation, resulting in a sterilized secondary challenge. However, as α$\alpha$ increases, the peak viral load rises progressively. This confirms that the primary driver of symptomatic secondary infection with variants is viral mutation (antigenic distance), rather than the waning of specific antibody titers against the ancestral strain.

##### 8.3 Quantification of Immune Imprinting

To rigorously assess immune imprinting, we defined two cross-reactivity metrics based on the population of high-affinity antibodies generated after secondary infection. Here, we analyze antibodies with indices *i* corresponding to high affinity (specifically indices 51, 61–62, 71–73, 81–84, and 91–95).

We define the metrics as follows:

1. **Imprinting Level (Cross Reaction$_1$):** The proportion of the secondary response derived from the recall of cross-reactive memory clones rather than *de novo* naive recruitment.
2. **Affinity Cross-Reactivity (Cross Reaction$_2$ ):** The ratio of the current specific response against the mutant to the original specific response against the wild-type.

**Equations:**

$$Cross\ reaction_1 = \frac{\sum_{i=1}^{n}\left(Ab(i) \times \frac{Ab(i)_{initial} - Base}{Ab(i)_{initial}}\right.}{\sum_{i=1}^{n} Ab(i)}$$

$$Cross\ reaction_2 = \frac{\sum_{i=1}^{n}\left(Ab(i) \times \frac{Ab(i)_{initial} - Base}{Ab(i)_{initial}}\right.}{\sum_{i=1}^{n} Ab_{wt}(i)}$$

**Auxiliary Variables:**

- **Effective Initial Memory:** $Ab(i)_{initial} = Ab_{wt}(i) \times 10^{-\alpha}$
- **Naive Baseline:** $Base = (1 - 10^{-\alpha})fN$
- $Ab(i)$: Concentration of high-affinity antibodies to the **mutant** after second infection.
- $Ab_{wt}(i)$: Concentration of high-affinity antibodies to the **wild-type** after first infection.

**8.4 Dynamics of Imprinting (Fig. S8B & S8C)**

**Natural Infection Scenarios (Figure S8B)**

Our simulations reveal a complex, non-monotonic relationship between the mutation coefficient $\alpha\alpha$ and the level of immune imprinting during natural reinfection:

1. **Sterilizing Immunity (α≤3):** Viral proliferation is suppressed. No significant reshaping of the repertoire occurs. The imprinting level is static (approximating $\frac{10^{-\alpha}FN}{(1-10^{-\alpha})fN+10^{-\alpha}FN}$ ).
2. **Recall Bias (α≈4):** Secondary infection occurs. The virus differs enough to replicate but is similar enough to trigger a massive recall of original memory B cells ("back-boosting"). This drives a sharp **increase** in the observed imprinting level.
3. **Immune Escape (α>4):** As mutation increases further, the ancestral memory clones bind too weakly to the mutant to compete effectively. The system is forced to recruit *de novo* naive B cells targeting the mutant's unique epitopes. Consequently, the relative contribution of imprinted memory **declines**.

This biphasic peak explains why partial immune escape often triggers the strongest "original antigenic sin" phenotypes.

**Vaccination Scenarios (Figure S8C)**

In contrast, simulation of booster vaccination with a mutant strain (Antigen dosage $10^{14}$) shows a different profile. Both $Cross\ Reaction_1$ and $Cross\ Reaction_2$ exhibit a monotonic downward trend as $\alpha$ increases. This suggests that high-dose antigen exposure (typical of vaccination) may more effectively overcome the affinity threshold required to recruit naive cells, thereby reducing the relative dominance of imprinted memory as the antigenic distance increases. Notably, even at $\alpha \to \infty$ (implying distinct viral origins), a residual imprinting level (approximately 10%) persists, reflecting inherent limits in the diversity of the germline repertoire.

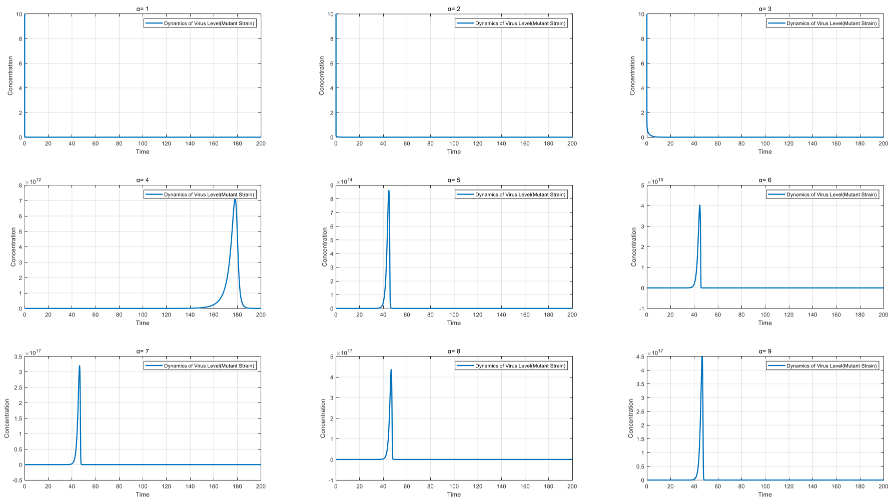


**Figure S8A: Viral dynamics under varying mutation coefficients.** Viral load trajectories during secondary infection. Breakthrough infection occurs only when the mutation coefficient α$\alpha$ exceeds a critical threshold ($\alpha$>3).

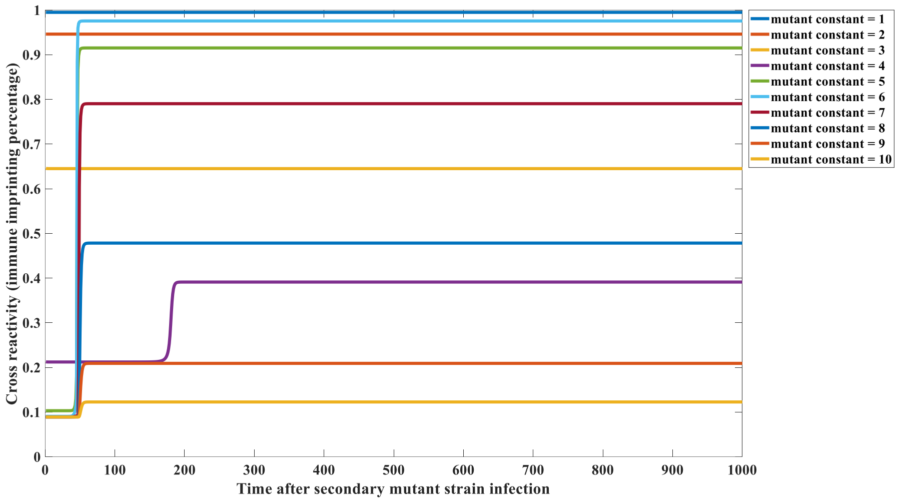


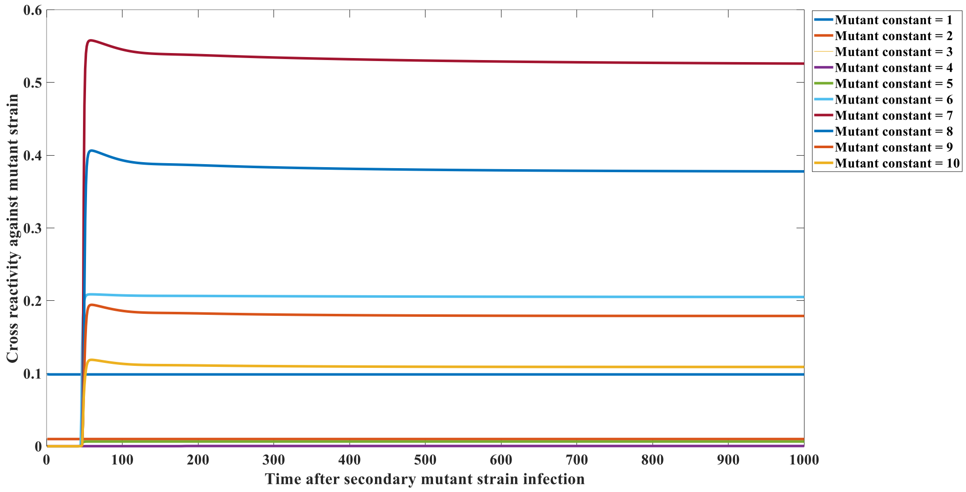


**Figure S8B: Immune imprinting in natural infection.** The relationship between antigenic distance ($\alpha$) and cross-reactivity indices reveals a "U-shaped" or intermediate peak behavior, where imprinting is maximized at intermediate mutation levels that permit infection but retain structural homology.

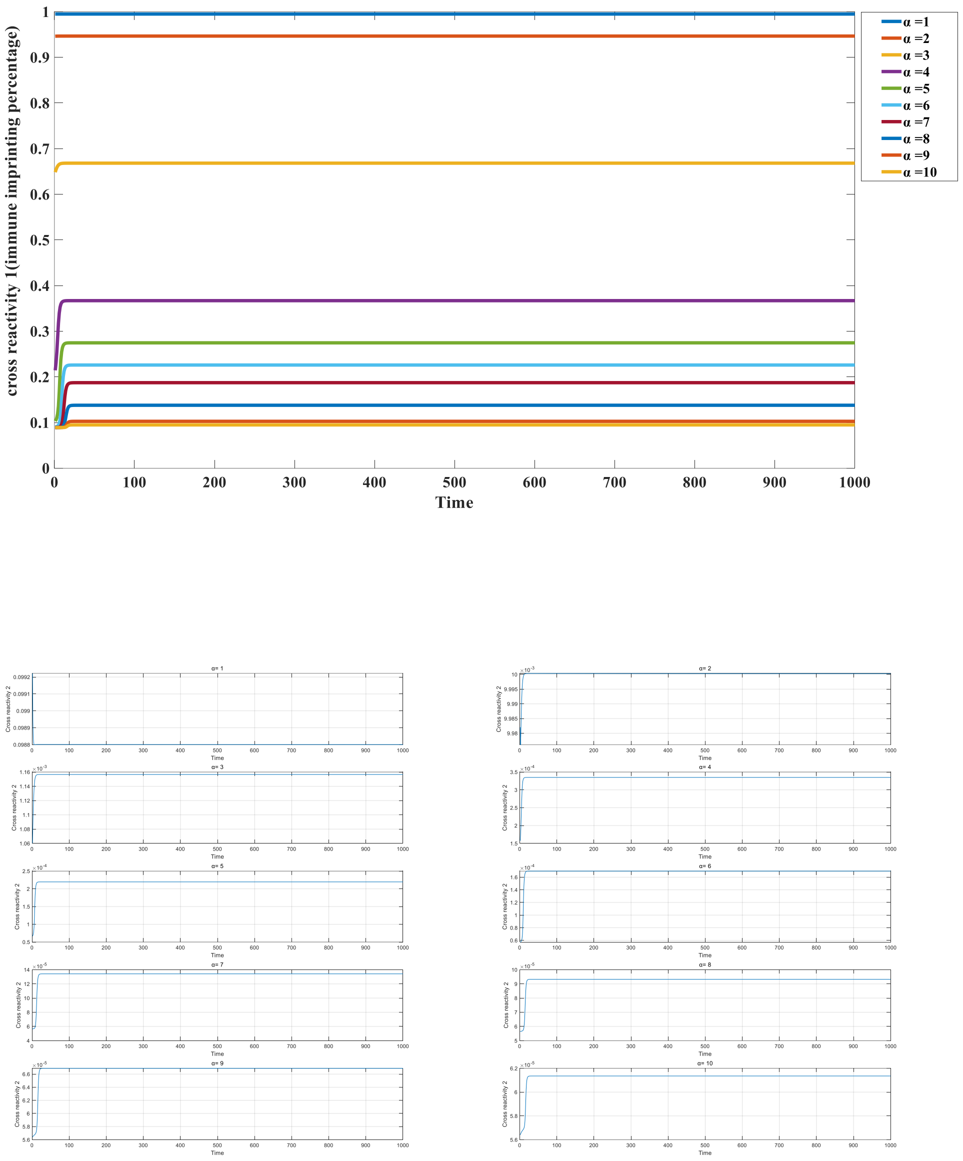


**Figure S8C: Immune imprinting following mutant-strain vaccination.** Unlike natural infection, vaccination leads to a monotonic decrease in imprinting signatures as antigenic distance increases.

#### 9 Dynamics and Parameterization of Model 3.2.1

##### 9.1 Model Equations and Biological Rationale

Model 3.2.1 describes the interplay between humoral and cellular immunity driven by two distinct antigen categories.

**Type I antigens** (e.g., SARS-CoV-2) are characterized by their ability to be directly recognized by Pattern Recognition Receptors (PRRs), such as Toll-like receptors (TLRs) and C-type lectin receptors (CLRs), on the surface of dendritic cells (DCs).

**Type II antigens** (e.g., self-antigens and LCMV) lack intrinsic PRR recognition motifs. Consequently, these antigens require opsonization—binding to antibodies to form antigen-antibody complexes (C)—before they can be recognized and internalized via Fc receptors.

The differential equations governing the dynamics of free virus/antigen (V), specific antibodies (A), antigen-antibody complexes (C), and cytotoxic T cells (Tc) are defined as follows:

**Common Dynamics (Both Types):**

$$\frac{dV}{dt} = k_0 V - k_1 VA + k_2 C;$$

$$\frac{dA}{dt} = -k_1 VA + k_2 C + k_3 C + \pi - k_4 A;$$

$$\frac{dC}{dt} = k_1 VA - k_2 C - k_5 C;$$

**Distinct Cellular Immunity Dynamics:**

For Type I antigens**,** Tc expansion is driven by both free antigen and immune complexes:

$$\frac{dTc}{dt} = k_6(C + V) + \pi_1 - k_7 Tc;$$

For Type II antigens**,** Tc expansion is strictly dependent on immune complexes:

$$\frac{dTc}{dt} = k_6 C + \pi_1 - k_7 Tc$$

Here, the varying dependence of Tc proliferation on V and C reflects the distinct recognition mechanisms described above. Therefore, the cellular immune response (proliferation capacity of Tc cells) exhibits a differential relationship with humoral immunity depending on the antigen type.

##### 9.2 Parameter Values

The parameters utilized in the simulations are detailed in **Supplementary Table S14**. Note that k3, representing antibody production stimulation, is modulated to simulate the effects of Rituximab treatment.

**Supplementary Table S14 | Parameters and initial values for Model 3.2.1**

| Variable/Parameter | Description | Value/Initial Condition |
|---|---|---|
| $V_0$ | Initial Antigen | $10^{10}$ |
| $A_0$ | Initial Antibody | $10^5$ |
| $C_0$ | Initial Complex | 0 |
| $Tc_0$ | Initial Cytotoxic T Cells | $10^3$ |
| $k_0$ | Antigen replication/decay rate | −0.05 |

| Variable/Parameter | Description | Value/Initial Condition |
|---|---|---|
| $k_1$ | Association rate constant | $10^{-7}$ |
| $k_2$ | Dissociation rate constant | $10^{-14}$ |
| $k_3$ | Antibody stimulation rate | 5 (Control); 3 (Rituximab) |
| $k_4$ | Antibody decay rate | 0.01 |
| $k_5$ | Complex clearance rate | 2 |
| $k_6$ | T cell stimulation rate | $10^{-5}$ |
| $k_7$ | T cell decay rate | 0.01 |
| $\pi$ | Basal Ab production | $10^3$ |
| $\pi_1$ | Basal T cell production | 10 |

##### 9.3 Simulation Results regarding Rituximab Treatment

Simulation outcomes for Type I and Type II antigens are illustrated in **Supplementary Fig. S9A** and **S9B**, respectively.

**Type I Antigen Dynamics (Fig. S9A):**
In the context of Type I antigens, subjects with suppressed humoral immunity (Rituximab treatment group) exhibit significantly reduced specific antibody proliferation compared to controls. Conversely, CD8+ T cell proliferation is markedly enhanced in this group. Mechanistically, this inverse relationship arises because the natural decay rate of the free antigen is significantly lower than the clearance rate of antigen-antibody complexes. In the control group, the abundance of available Fc receptors (e.g., on NK cells) facilitates rapid degradation of immune complexes. While these complexes stimulate DC maturation and Tc activation, their short half-life limits the duration of the stimulus. By contrast, when antibody production is restricted, free antigen persists longer, providing sustained stimulation via PRR pathways and resulting in a robust Tc response.

**Type II Antigen Dynamics (Fig. S9B):**
The dynamics for Type II antigens differ fundamentally. Because DCs require antigen-antibody complexes for recognition and subsequent T cell priming, cellular immunity activation is positively correlated with the magnitude of the humoral response. Consequently, in scenarios where humoral immunity is suppressed (Rituximab treatment), the formation of immune complexes is limited, leading to the concurrent suppression of cellular immune functions.

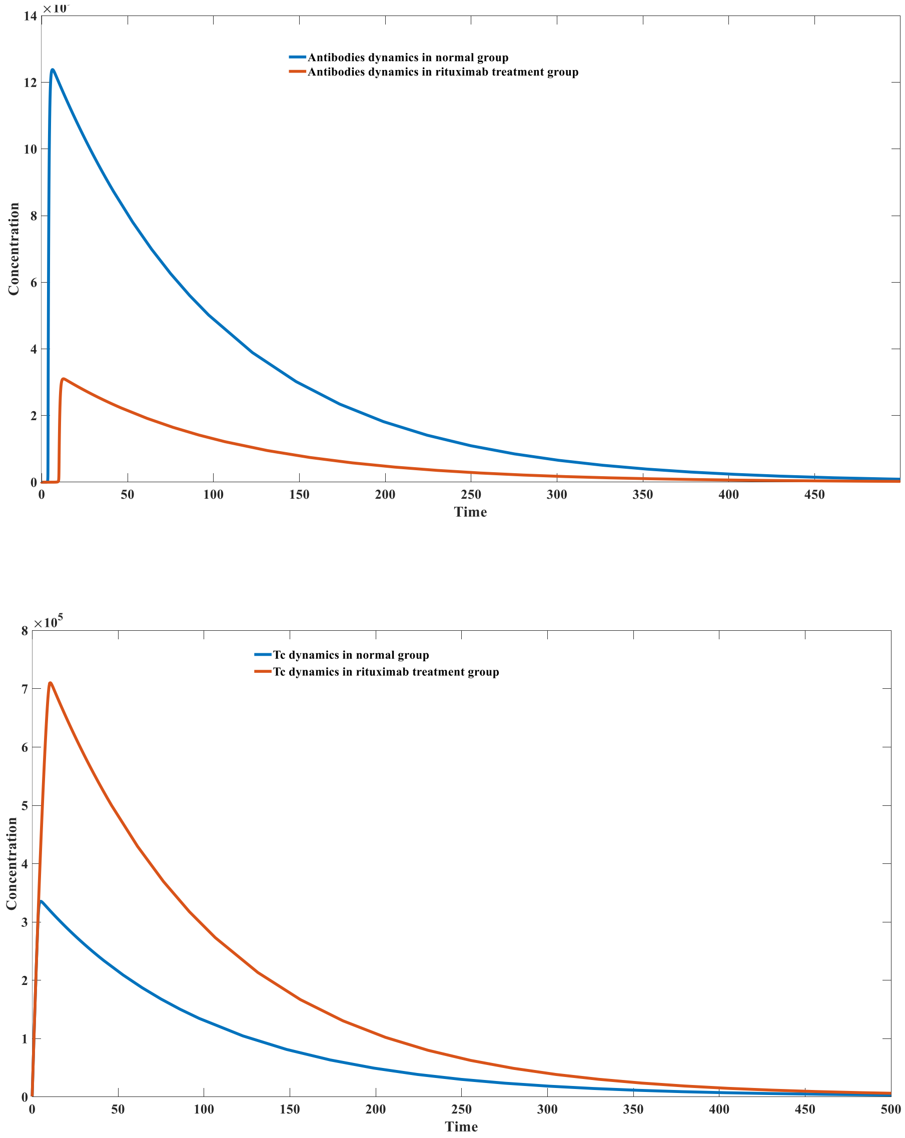


**Fig. S9A | Divergent trends in humoral and cellular immune responses following Type I antigen vaccination under Rituximab treatment.**

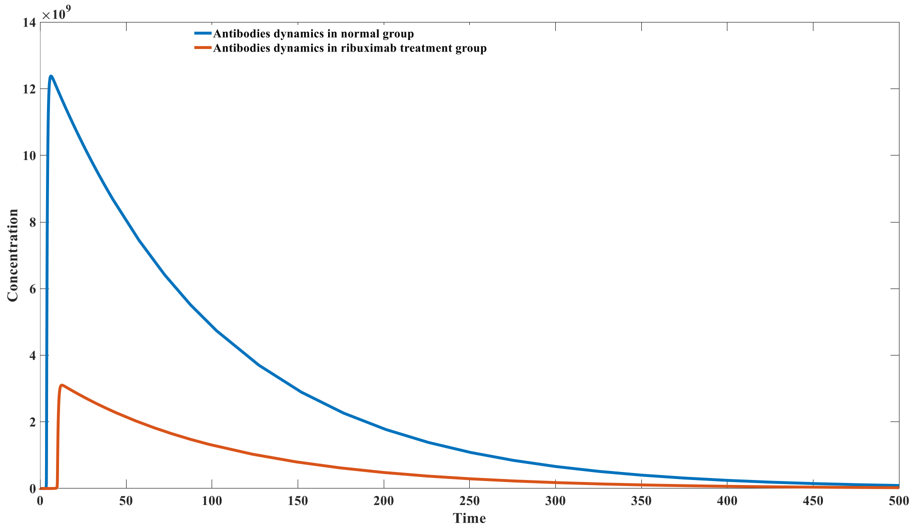


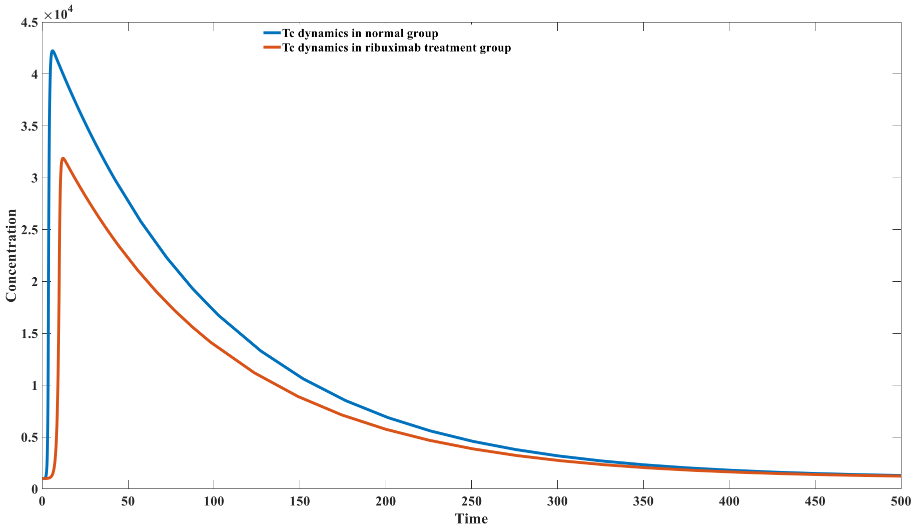


**Fig. S9B | Concurrent reduction of humoral and cellular immune responses following Type II antigen vaccination under Rituximab treatment.**

#### 10 Dynamics and Parameterization of Model 3.2.2

##### 10.1 Model Equations and Rationale

Model 3.2.2 defines the kinetic relationship between CD8+ T cell cytotoxicity and the intracellular antigen concentration of infected cells. This formulation functions as the core cellular immunity component within the broader framework of Model 3.4.1.

The model describes the interaction between cytotoxic T cells ($Tc$) and $n$ populations of infected target cells ($N_i$), resulting in the formation of immunological effector complexes ($C_i$). The governing differential equations are as follows:

$$\frac{dTc}{dt} = \sum_{n}^{i=1} -k_1 N_i Tc + k_{2,i} C_i \,;$$

$$\frac{dN_i}{dt} = -k_1 N_i Tc + k_{2,i} C_i;$$

$$\frac{dC_i}{dt} = k_1 N_i Tc - k_{2,i} C_i;$$

The dissociation or reaction rate constant $k_{2,i}$ is not static; rather, it is modulated by the antigen density per cell ($V_i/N_i$). This relationship follows a logarithmic saturation function defined by:

$$log\ (k_{2i}) = \mu - \tau \frac{V_i/N_i}{V_i/N_i + Km};$$

This equation captures the functional dependence of T cell efficacy on the antigen load presented by the target cells.

##### 10.2 Parameter Values

The specific parameters utilized in Model 3.2.2 are listed in **Supplementary Table S15**.

**Supplementary Table S15 | Parameters and values for Model 3.2.2**

| Parameter | Description | Value |
|---|---|---|
| $k_1$ | Association rate constant | 0.01 |
| μ | Baseline logarithmic rate factor | 12 |
| τ | Rate modulation scaling factor | 16 |
| $K_m$ | Michaelis constant for antigen saturation | $10^3$ |

#### 11 Mathematical Analysis and Parameterization of Model 3.3.1

##### 11.1 Model Formulation

Model 3.3.1 describes the multiscale dynamics of the adaptive immune response, integrating B cell receptor (BCR) and T cell receptor (TCR) repertoire diversity. The system tracks the kinetics of IgM and IgG producing B cells/antibodies, naive/effector ($T^1$) and memory ($T^2$) T cells, antigen-antibody complexes ($C$), antigen-presenting complexes ($D$), and immunological synapses/clusters ($F$).

The governing differential equations for the $i$-th B cell/antibody clone and $j$-th T cell clone are defined as follows:

###### 11.1.1. B-Cell Clonal Dynamics and Isotype Switching

The expansion of IgM ($B_i^{IgM}$) and IgG ($B_i^{IgG}$) B-cell clones is driven by signals received from immunological synapses formed with CD4+ T cells.

$$\frac{dB_i^{IgM}}{dt} = \pi f_i^B - p_1 E B_i^{IgM}/\beta + p_2 C_{2,i} - k_{1,i} V B_i^{IgM}/\beta + k_{2,i} C_{1,i} + \beta\gamma \sum_{j=1}^{m} \left[\alpha k_3\left(\mathcal{F}_{1,ij} + \mathcal{F}_{2,ij}\right) + k_4\left(\mathcal{F}_{3,ij} + \mathcal{F}_{4,ij}\right)\right] - d_1 B_i^{IgM};$$

$$\frac{dB_i^{IgG}}{dt} = -p_1 E B_i^{IgG}/\beta + p_2 C_{4,i} - k'_{1,i} V B_i^{IgG}/\beta + k'_{2,i} C_{3,i} + \beta\gamma \sum_{j=1}^{m} \left[(1-\alpha) k_3\left(\mathcal{F}_{1,ij} + \mathcal{F}_{2,ij}\right) + k_3\left(\mathcal{F}_{5,ij} + \mathcal{F}_{6,ij}\right) + k_4\left(\mathcal{F}_{7,ij} + \mathcal{F}_{8,ij}\right)\right] - d_2 B_i^{IgG};$$

Here, $\mathcal{F}_{k,ij}$ represents the synapse complex formed between B cell clone i and T cell clone j. The term β scales the BCR density per cell, and γ quantifies the stoichiometric amplification of B-cell proliferation induced by T-cell help. α denotes the probability of IgM-to-IgG class switching upon T-cell priming.

##### 11.1.2. CD4+ T Cell Recruitment and Memory

T cells are recruited into synapses by pMHC-II complexes (D) presented on B cells.

$$\frac{dT_j^{(1)}}{dt} = \pi_1 f_j^T - T_j^{(1)} \sum_{i=1}^{n} \sum_{k\in\Psi} \sigma_{1,i} D_{k,i} + \sum_{i=1}^{n} \sum_{k\in\Psi} \sigma_{2,i} \mathcal{F}_{k,ij} - T_j^{(1)} \sum_{i=1}^{n} \sum_{k\in\Psi} \omega_1 D_{k,i} + \sum_{i=1}^{n} \sum_{k\in\Psi} \omega_2 \mathcal{F}_{k,ij} + P(T) - d_3 T_j^{(1)};$$

$$\frac{dT_j^{(2)}}{dt} = -T_j^{(2)} \sum_{i=1}^{n} \sum_{k\in\Psi} \sigma_{1,i} D_{k,i} + \sum_{i=1}^{n} \sum_{k\in\Psi} \sigma_{2,i} \mathcal{F}_{k,ij} - T_j^{(2)} \sum_{i=1}^{n} \sum_{k\in\Psi} \omega_1 D_{k,i} + \sum_{i=1}^{n} \sum_{k\in\Psi} \omega_2 \mathcal{F}_{k,ij} + P_{mem}(T) - d_4 T_j^{(2)};$$

where P(T) represents the T-cell proliferation feedback driven by successful synapse formation.

$$P(T) = \alpha k_3 \sum_{i=1}^{n} \mathcal{F}_{1,ij} + \alpha k_3 \sum_{i=1}^{n} \mathcal{F}_{5,ij} + k_4 \sum_{i=1}^{n} \mathcal{F}_{3,ij} + k_4 \sum_{i=1}^{n} \mathcal{F}_{7,ij}$$

$$P_{mem}(T) = (1-\alpha) \mathrm{k}_3 \sum_{i=1}^{n} (\mathcal{F}_1)_{\mathrm{ij}} + (1-\alpha) \mathrm{k}_3 \sum_{i=1}^{n} (\mathcal{F}_5)_{\mathrm{ij}} + \mathrm{k}_3 \sum_{i=1}^{n} (\mathcal{F}_2)_{\mathrm{ij}} + \mathrm{k}_3 \sum_{i=1}^{n} (\mathcal{F}_6)_{\mathrm{ij}} + \mathrm{k}_4 \sum_{i=1}^{n} (\mathcal{F}_4)_{\mathrm{ij}} + \mathrm{k}_4 \sum_{i=1}^{n} (\mathcal{F}_8)_{\mathrm{ij}}$$

##### 11.1.3. Antigen Processing and Presentation Pipeline

The model tracks the lifecycle of the antigen from surface binding (C) to endosomal degradation and surface presentation as pMHC-II (D).

**Step 1: Surface Binding (Representative for Virus-IgM):**

$$\frac{dC_{1,i}}{dt} = \frac{k_{1,i}V}{\beta} B_i^{IgM} - k_{2,i}C_{1,i} - (k_5 + d_5)C_{1,i};$$

$$\frac{dC_{2,i}}{dt} = \frac{p_1 E}{\beta} B_i^{IgM} - p_2 C_{2,i} - (k_5 + d_5)C_{2,i};$$

$$\frac{dC_{3,i}}{dt} = \frac{k_{1,i}V}{\beta} B_i^{IgG} - k_{2,i}C_{3,i} - (k_5 + d_5)C_{3,i};$$

$$\frac{dC_{4,i}}{dt} = \frac{p_1 E}{\beta} B_i^{IgG} - p_2 C_{4,i} - (k_5 + d_5)C_{4,i};$$

Here, $k_5$ represents the rate of internalization and processing into pMHC-II, while $d_5$ denotes non-productive degradation.

**Step 2: pMHC-II Presentation:**

$$\frac{dD_{1,i}}{dt} = k_5 C_{1,i} - \sum_{j=1}^{m} (D_{1,i}T_j^{(1)}\sigma_{1,i} - \sigma_{2,i}\mathcal{F}_{1,ij} + D_{1,i}T_j^{(2)}\sigma_{1,i} - \sigma_{2,i}\mathcal{F}_{2,ij}) - d_6 D_{1,i};$$

$$\frac{dD_{2,i}}{dt} = k_5 C_{2,i} - \sum_{j=1}^{m} (D_{2,i}T_j^{(1)}\sigma_{1,i} - \sigma_{2,i}\mathcal{F}_{3,ij} + D_{2,i}T_j^{(2)}\sigma_{1,i} - \sigma_{2,i}\mathcal{F}_{4,ij}) - d_6 D_{2,i};$$

$$\frac{dD_{3,i}}{dt} = k_5 C_{3,i} - \sum_{j=1}^{m} (D_{3,i}T_j^{(1)}\sigma_{1,i} - \sigma_{2,i}\mathcal{F}_{5,ij} + D_{3,i}T_j^{(2)}\sigma_{1,i} - \sigma_{2,i}\mathcal{F}_{6,ij}) - d_6 D_{3,i};$$

$$\frac{dD_{4,i}}{dt} = k_5 C_{4,i} - \sum_{j=1}^{m} (D_{4,i}T_j^{(1)}\sigma_{1,i} - \sigma_{2,i}\mathcal{F}_{7,ij} + D_{4,i}T_j^{(2)}\sigma_{1,i} - \sigma_{2,i}\mathcal{F}_{8,ij}) - d_6 D_{4,i};$$

$D_{1,i}$ represents the viral peptide-MHC complex derived from IgM-mediated uptake. $D_{2,i}, D_{3,i}, D_{4,i}$ correspond to complexes derived from environmental antigen (IgM), viral antigen (IgG), and environmental antigen (IgG), respectively.

**Step 3: B-T Synapse Formation ($\mathcal{F}$):**

The formation of the cognate synapse $\mathcal{F}_{k,ij}$ is governed by:

$$\frac{d\mathcal{F}_{1,ij}}{dt} = \sigma_{1,i}D_{1,i}T_j^{(1)} - \sigma_{2,i}\mathcal{F}_{1,ij} - d_7\mathcal{F}_{1,ij};$$

This generic form applies to all 8 permutations of interaction (Naive/Memory T cells × 4 types of Antigen sources).

##### 11.1.4. Global Antigen Dynamics

$$\frac{dV}{dt} = k_0 V - \sum_{i=1}^{n}\left(\frac{k_{1,i}VB_i^{IgM}}{\beta} - k_{2,i}C_{1,i}\right) - \sum_{i=1}^{n}\left(\frac{k'_{1,i}VB_i^{IgG}}{\beta} - k'_{2,i}C_{3,i}\right);$$

$$\frac{dE}{dt} = \pi_2 - \sum_{i=1}^{n}\left(\frac{p_1 EB_i^{IgM}}{\beta} - p_2 C_{2,i}\right) - \sum_{i=1}^{n}\left(\frac{p_1 EB_i^{IgG}}{\beta} - p_2 C_{4,i}\right);$$

**Parameter Definitions:**

- $\sigma_{1,i}, \sigma_{2,i}$: Association and dissociation rates for the TCR–pMHC-II interaction.
- $\omega_1, \omega_2$: Kinetic constants for interactions involving environmental antigens (assumed to have lower/baseline affinity).
- $f_i^B, f_j^T$: Initial precursor frequencies for B and T cells, distributed normally based on affinity to reflecting the naïve repertoire.
- $d_6, d_7$: Turnover rates for pMHC-II complexes and immunological synapses, respectively.

**11.2 Parameter Selection and Equilibrium Analysis**

General parameters and initial conditions are provided in **Supplementary Table S16**. Parameter selection adhered to the stability protocols established in Model 3.1.7, identifying values that maintain a homeostatic, disease-free equilibrium. Equilibrium concentrations were estimated through extensive literature mining and theoretical constraints.

Key homeostatic assumptions include:

1. A 1:1 stoichiometric ratio between total T cell and B cell populations at equilibrium.
2. A 1:1 ratio between memory and effector cell subsets.
3. A scaling factor $\beta=10^5$ to account for the multiplicity of BCRs on a single B cell relative to the T cell population.

In **Table S16**, blue-highlighted parameters indicate fixed experimental or literature-derived values, while red-highlighted parameters were derived to satisfy equilibrium constraints. Mathematical analysis and numerical simulations confirm that with an initial virus load $V_0$ =0, the system maintains a stable steady state without spontaneous fluctuations.

**Supplementary Table S16 | Parameters and initial values for Model 3.3.1**

| Compartment | Initial Value | Parameter | Value |
|---|---|---|---|
| $V_0$ | 10 | $k_0$ | 0.8 |
| $E_0$ | $10^{15}$ | $k_3$ | 2 |
| $\sum_{i=1}^{n} B_i^{IgM}$ | $10^{15}$ | $k_4$ | 1 |
| $\sum_{i=1}^{n} B_i^{IgG}$ | $10^{15}$ | $k_5$ | 0.5 |
| $T^1, T^2$ | $10^{10}$ | $p_1$ | 10−16 |
| $C_1, C_3$ | 0 | $p_2$ | ≈1.77 (10/3.6 −1) |
| $C_2, C_4$ | $3.6\times10^{13}$ | $d_1, d_4$ | 0.008 |

| Compartment | Initial Value | Parameter | Value |
|---|---|---|---|
| $D_1$,$D_3$ | 0 | $d_2$ | 0.004 |
| $D_2$,$D_4$ | $10^{13}$ | $d_3$ | 0.016 |
| $F_{1,2,5,6}$ | 00 | $d_5$ | 0.5 |
| $F_{3,4,7,8}$ | $10^8$ | $d_6$,$d_7$ | 0.6 |
| | | $\omega_1$ | $10^{-4}$ |
| | | $\omega_2$ | $10^6$–0.6 |
| | | $\alpha$ | 0.95 |
| | | $\beta$ | $10^5$ |
| | | $\gamma$ | 2 |
| | | $\pi$ | $1\times10^{12}$ |
| | | $\pi_1$ | $0.8\times10^8$ |
| | | $\pi_2$ | $7.2\times10^{13}$ |

##### 11.3 Repertoire Diversity and Affinity settings

To computationally manage the high dimensionality of adaptive immune repertoires, we simplified the combinatorial complexity. Instead of simulating a full 100×100 matrix of BCR and TCR interactions, two distinct schemes were employed:

1. **Low BCR / High TCR Diversity:** 2 BCR types × 100 TCR types.
2. **High BCR / Low TCR Diversity:** 100 BCR types × 2 TCR types.

**Case 1 Configuration:**

- **BCR Affinity:** The two BCR types were assigned distinct affinity profiles for IgM and IgG.
    - IgM affinities: $k_{1,1}=5\times10^{-12}$, $k_{2,1}=10^{-3}$; $k_{2,1}=10^{-3}$, $k_{2,2}=10$.
    - IgG affinities: $k_{1,1}'=10^{-12}$, $k_{2,1}'=10^{-3}$; $k_{1,2}'=10^{-16}$, $k_{2,2}'=10$.
    - The high-affinity BCRs constitute a fraction of $10^{-5}$ of the total, with low-affinity BCRs comprising the remainder.
- **TCR Diversity:** 100 TCR phenotypes were defined, indexed by $i=10(m-1)+n$ (where $m,n\in[1,10]$).
    - The positive affinity coefficient to MHC-II ($\sigma_{1i}$) is defined as $10^{(m-13)}$.
    - The dissociation coefficient ($\sigma_{2i}$) is defined as $10^{(n-4)}$.
    - The distribution of initial effector and memory CD4+ T cells follows a probability function $f_i^T$ derived from a normal distribution of log-affinity values: $log(K) \sim N(5,0.82)$.

Simulation results for this configuration at equilibrium and under viral infection are presented in **Supplementary Fig. S10A** and **S10B**, respectively.

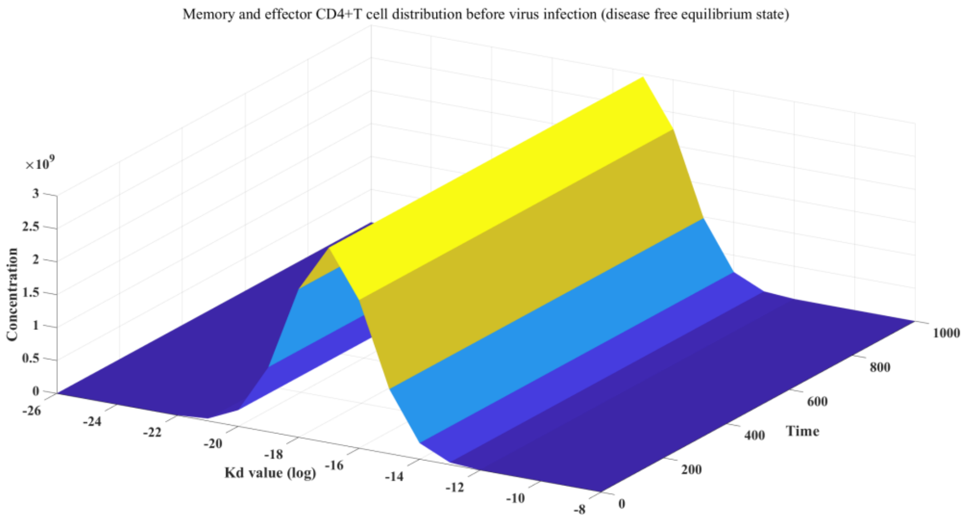


Figure S10_A: Disease free equilibrium state in B-CD4+T interaction model before infection (BCR diversity = 2; TCR diversity = 100).

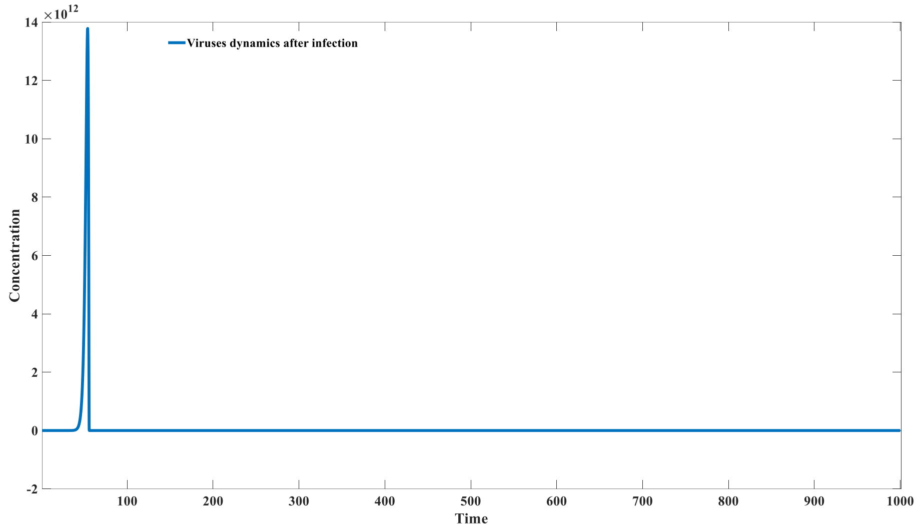


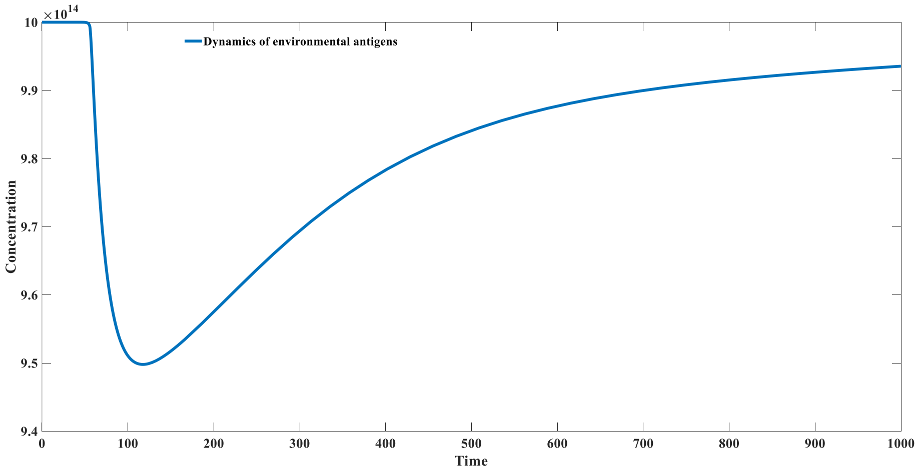

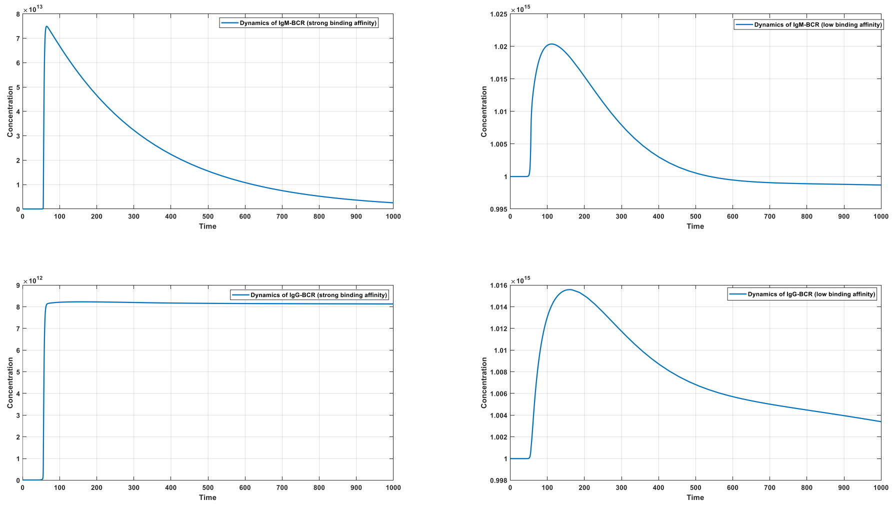
Dynamics of IgM-BCR (strong binding affinity)
Dynamics of IgM-BCR (low binding affinity)
Dynamics of IgG-BCR (strong binding affinity)
Dynamics of IgG-BCR (low binding affinity)
Concentration
Time

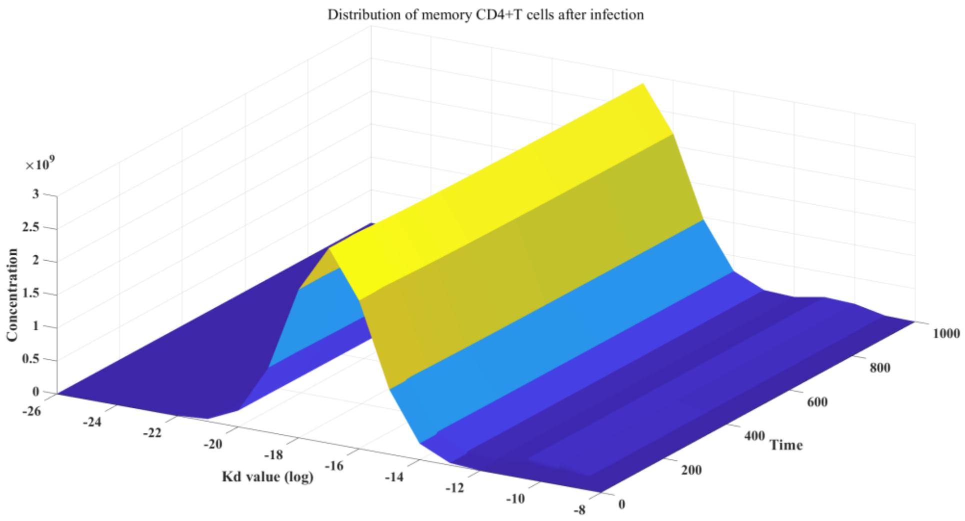
Distribution of memory CD4+T cells after infection
Concentration
Kd value (log)
Time

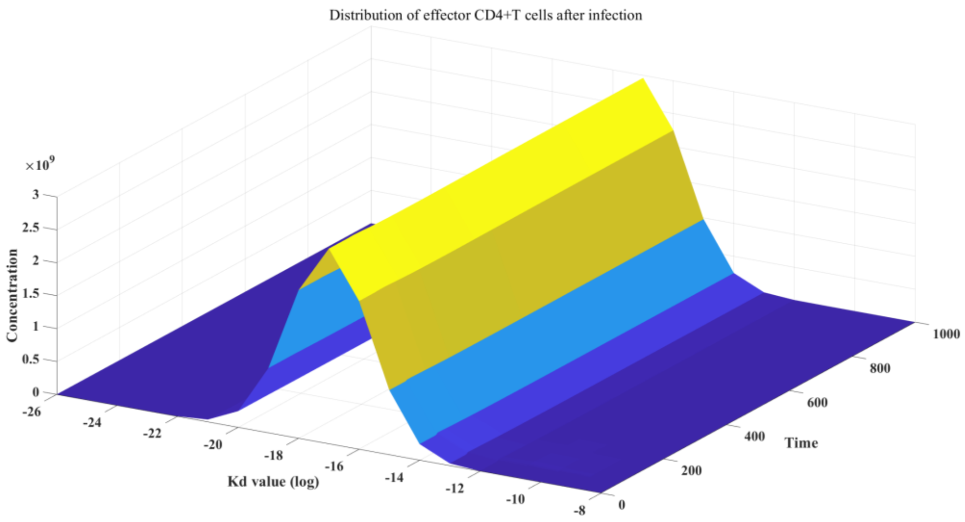
Distribution of effector CD4+T cells after infection
Concentration
Kd value (log)
Time

Figure S10_B: Dynamics of host-virus interaction in B-CD4+T interaction model after infection (BCR diversity = 2; TCR diversity = 100).

**Case 2 Configuration:**

In the second configuration, we modeled a restricted T cell repertoire coupled with a diverse B cell repertoire (2 TCR clonotypes × 100 BCR clonotypes).

**T Cell Receptor (TCR) Parameters:**
The two TCR populations were defined with the following MHC-II affinity coefficients:

- **High-affinity TCR:** $\sigma_{1,1}=5\times10{-}12$, $\sigma_{2,1}=10{-}3$. This phenotype represents $10^{-5}$ of the total population.
- **Low-affinity TCR:** $\sigma_{1,2}=5\times10{-}16$, $\sigma_{2,2}=10$. This phenotype comprises the remaining proportion $(1-10^{-5})$.

**B Cell Receptor (BCR) Parameters:**
We defined 100 distinct BCR clonotypes, indexed by $i=10(m-1)+n$ (where $m,n\in[1,10]$). This diversity applies to both naive (IgM) and memory (IgG) subsets, though with differing kinetic properties.

- **IgM-BCRs:**
  - Association rate constant: $k_{1,i}=5\times10^{(m-22)}$.
  - Dissociation rate constant: $k_{2,i}=10^{(n-4)}$.
  - Initial distribution: The probability $f(IgM)_i$ follows a log-normal distribution of affinity coefficients: $log(K){\sim}N(5,0.82)$.
- **IgG-BCRs:**
  - Association rate constant: $k_{1,i}'=10^{(m-22)}$.
  - Dissociation rate constant: $k_{2,i}'=10^{(n-4)}$.
  - Initial distribution: $f(IgG)_i$ follows the same probability distribution defined for IgM.

Simulation results illustrating the host-virus interaction dynamics under this specific diversity configuration are presented in **Supplementary Fig. S10C**.

**11.4 Investigation of Memory-Mediated Protection**

To evaluate the relative contributions of cellular and humoral memory in preventing secondary infections, we established a two-phase simulation protocol.

1. **Primary Phase:** A primary infection was simulated over $t\in[0,1000]$.
2. **Secondary Phase:** The system was re-initialized to simulate a secondary challenge. To isolate specific memory effects, one arm of the adaptive immune system was retained at its post-primary equilibrium state (values at $t$=1000), while the other was reset to naive baseline levels.

**CD4+ T Cell Memory (Fig. S10D):**
In this scenario, CD4+ T cell compartments (both memory and effector subsets) were initialized at their post-infection levels, while B cell compartments were reset to naive states. As shown in **Fig. S10D**, the presence of established CD4+ T cell memory reduced the peak viral load compared to a primary infection but failed to prevent the infection entirely.

**B Cell Memory (Fig. S10E):**
Here, B cell compartments (IgM and IgG populations) were retained at post-infection levels, while T cells were reset. The results, displayed in Fig. S10E.

#### 12 Cohort-Based Model of Temporal Viral Infection Dynamics

##### 12.1 Model Equations and Formulation

Model 3.4.1 employs a cohort-based approach to simulate the temporal evolution of host-virus interactions. The concentration updates over time step $\Delta t$ utilize the baseline parameters established in Model 3.1.7. However, specific modifications were introduced to account for the non-cellular extracellular environment and the cooperative effects of T cell help on B cell clonal expansion.

1. **Extracellular Context:** In this specific integration phase, the intrinsic viral replication term is set to zero ($k_0$=0 in the context of extracellular interaction), focusing on interaction and decay dynamics.
2. **T Cell help ($\delta$):** To model the non-linear enhancement of B cell proliferation driven by CD4+ T cells, we introduced an exponent $\delta$=1.25.

##### 12.2 Parameter Values

The parameters characterizing Model 3.4.1 are provided in **Supplementary Table S17**.

**Supplementary Table S17 | Parameters and initial values for Model 3.4.1**

| Parameter | Value | Description |
|---|---|---|
| $k_m$ | $2\times10^{6}$ | Michaelis constant |
| $k_4$ | $10^{-3}$ (Baseline) | Antigen-BCR complex regeneration/proliferation rate |
| $k_5$ | 5 | IgM-BCR decay rate |
| $k_6$ | $10^{7}$ | IgG-BCR decay rate |
| $k_7$ | $10^{-3}$ | IgM secretion rate |
| $gen_c$ | $5\times10^{-5}$ | Generation coefficient |
| $threshold_1$ | $10^{14}$ | Threshold constant 1 |
| $threshold_2$ | $10^{9}$ | Threshold constant 2 |

##### 12.3 Simulation Results

**Impact of Individual Immune Strength ($k_4$):**
The coefficient $k_4$, representing the regeneration rate of antigen-BCR complexes, serves as a proxy for individual immune responsiveness (clonal expansion potential). We compared scenarios of different immune responses on infection persistence.

As shown in **Supplementary Fig. S11**, a lower immune proliferation rate fails to clear the pathogen, resulting in persistent viral loads and stable infected cell populations. Conversely, a robust response leads to rapid viral elimination and resolution of infected cells, as illustrated in **Supplementary Fig. S11**.

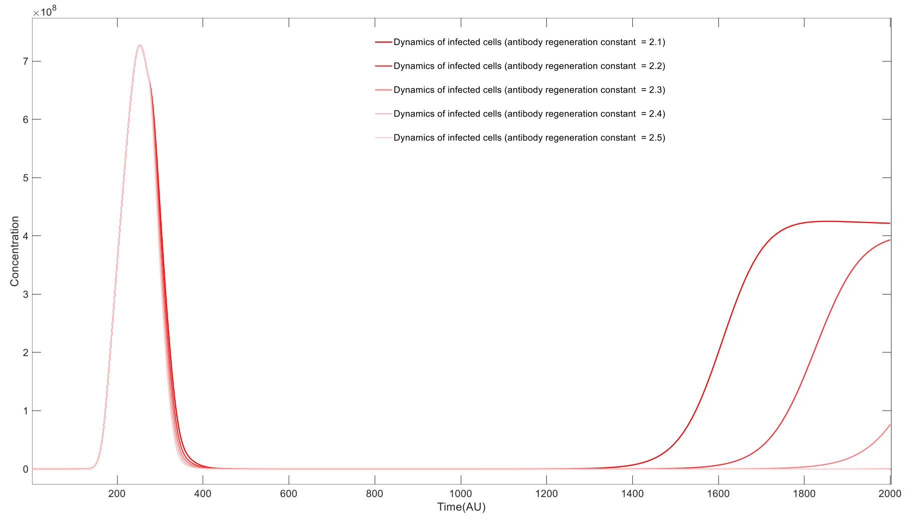


**Figure S11:** Impact of Individual Immune Strength on chronic infection

**Influence of Viral Replication Rate ($k_0$):**
We investigated the relationship between viral replication kinetics and chronicity. Counter-intuitively, a lower viral replication coefficient increases the propensity for chronic infection compared to a higher rate (refer to **Fig. S12**). This phenomenon suggests that slow-replicating viruses may fall below the activation threshold required to trigger a sterilizing immune response, establishing a low-level equilibrium.

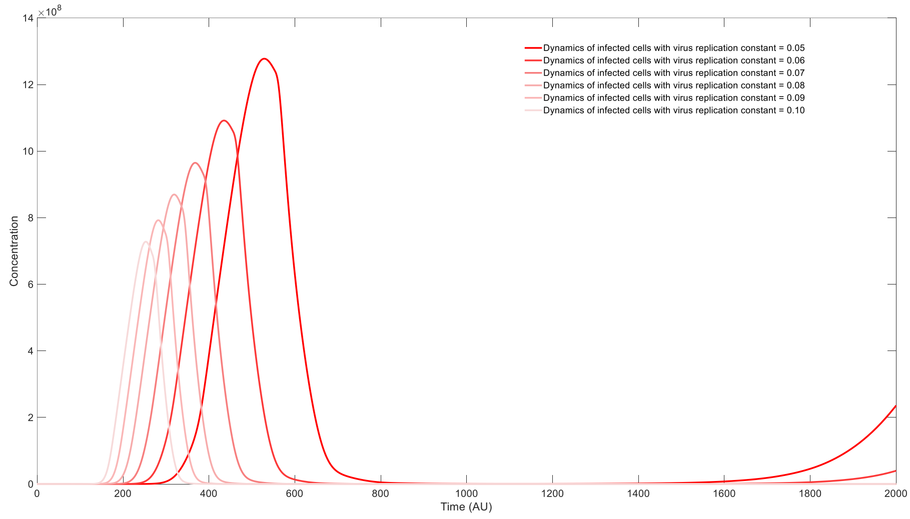


**Figure S12:** Impact of Viral Replication Rate on chronic infection

**Immune Imprinting and Secondary Infections:**

We explored the dynamics of secondary infections caused by mutant strains with varying degrees of antigenic distance (mutation coefficient $\alpha$).

- **Goldilocks Zone for Chronicity:** As indicated in **Supplementary Fig. S13**, the likelihood of chronic secondary infection initially increases with $\alpha$. At low $\alpha$, pre-existing antibodies effectively clear the virus. At intermediate $\alpha$ values, high titers of cross-reactive but non-neutralizing antibodies from the primary infection suppress the early expansion of the viral load (*sterilizing immunity failure*) but also prevent the rapid triggering of *specific* B cell expansion (*proliferation suppression*). This creates an equilibrium trap leading to chronic infection. At very high $\alpha$, the virus escapes recognition entirely, triggering a de novo acute response.

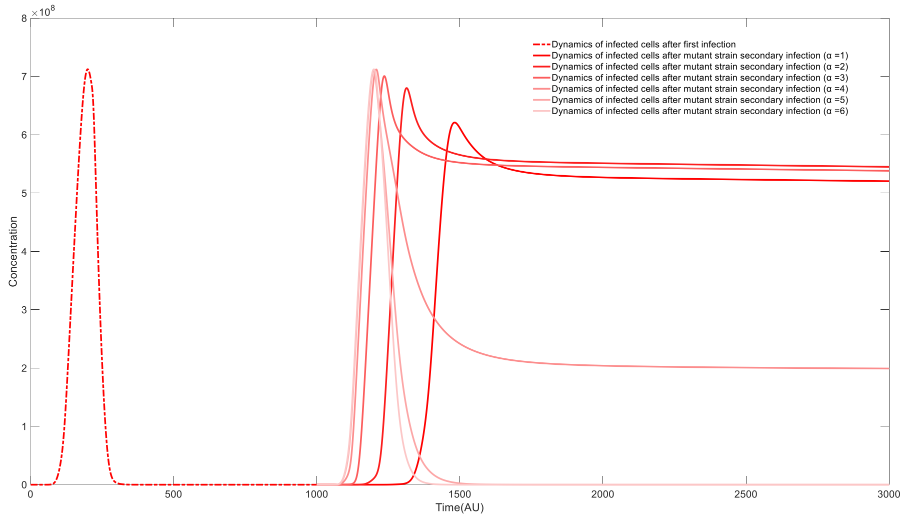


**Figure S13:** Impact of Immune Imprinting on chronic infection

**Therapeutic Interventions for Chronic Infection:**

Three distinct therapeutic modalities were modeled (simulated at $t$=200 during established chronic phase):

1. **Monoclonal Antibody (mAb) Therapy:** Administration of high-affinity mAbs ($Kd \approx 10^{-13}$) at doses of $2\times10^{16}$ to $2^{10}\times10^{16}$ (**Supplementary Fig. S14**) produced a rapid but transient reduction in viral load. Chronic infection re-established following antibody clearance using linear decay kinetics.
2. **Antigen-Specific Immunotherapy (Self-Antigen):** Administration of self-antigen (at doses of $2\times10^{18}$ to $2^{8}\times10^{18}$) temporarily alleviated chronic burden but failed to induce long-term resolution **(Supplementary Fig. S15)**.
3. **Therapeutic Vaccination:** Administration of therapeutic vaccine at doses of $2\times10^{11}$ to $2^{10}\times10^{11}$ (**Supplementary Fig. S16**) produced varied effects on overcome chronic bottleneck. Therapeutic vaccine challenge delivering viral antigens at high doses successfully disrupted the chronic equilibrium, leading to complete viral clearance.

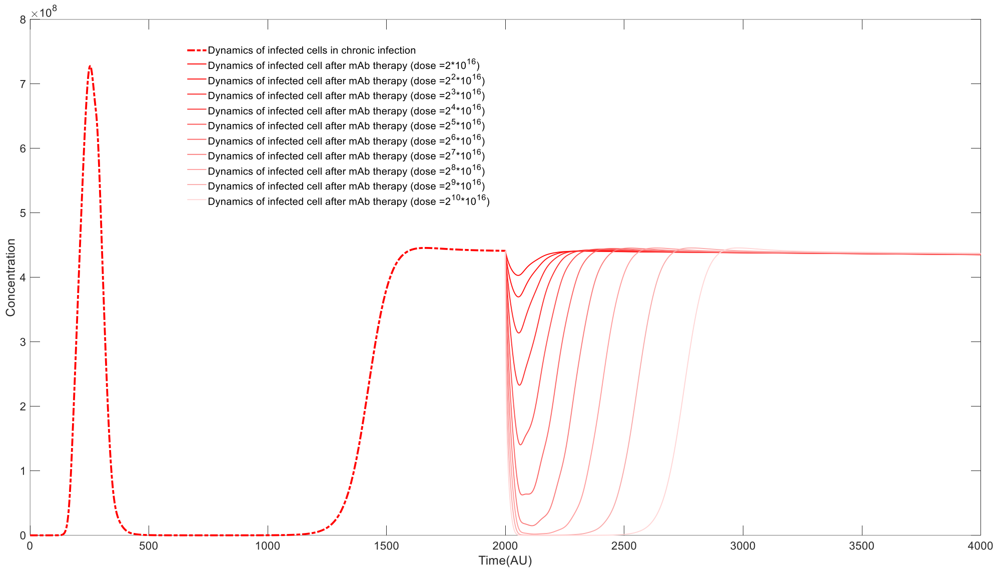


**Figure S14:** Monoclonal Antibody (mAb) Therapy on chronic infection

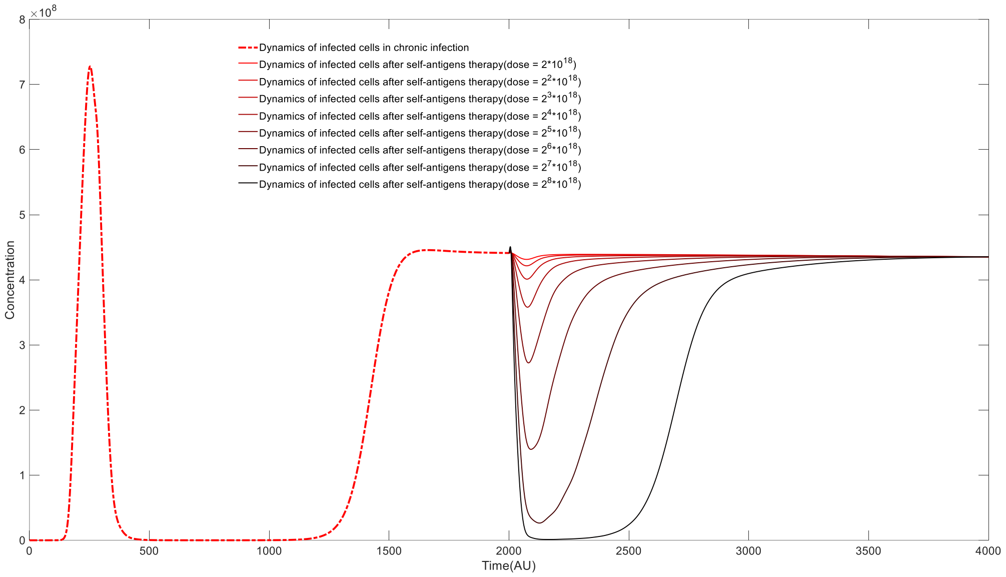


**Figure S15:** Antigen-Specific Immunotherapy (Self-Antigen) on chronic infection

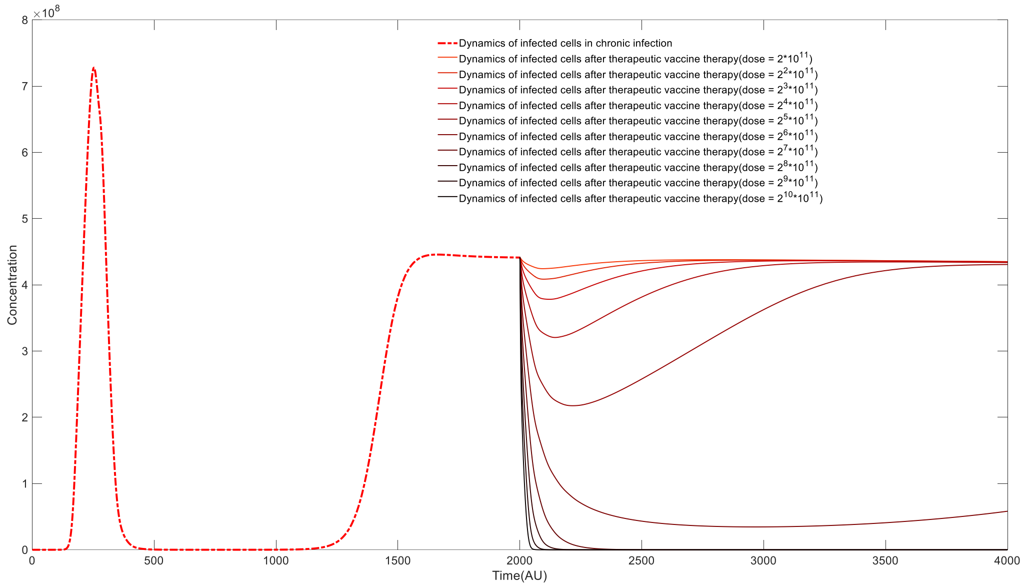


**Figure S16:** Impact of Therapeutic Vaccination on chronic infection

#### 13 Mathematical Analysis and Simulation of Tumor-Immune Dynamics (Model 3.5.1)

##### 13.1 Model Formulation

Model 3.5.1 captures the dynamic interactions between tumor cells, the adaptive immune system (antibodies and cytotoxic T cells), and tumor-derived neoantigens. The governing differential equations are defined as follows:

$$\frac{dT_u}{dt} = \pi_1 + k_1 T_u - d_1 T_u - \beta \frac{A}{A + k_m} T_u - \alpha T_c T_u;$$

$$\frac{d(Neo)}{dt} = -k_2(Neo)A + k_3 C + \rho_1\left(d_1 T_u + \beta \frac{A}{A + k_m} T_u + \alpha T_c T_u\right);$$

$$\frac{dC}{dt} = k_2(Neo)A - k_3 C - k_4 C;$$

$$\frac{dA}{dt} = -k_2(Neo)A + k_3 C + k_5 C + \pi_2 - d_2 A;$$

$$\frac{dT_c}{dt} = -\rho_2 \alpha T_c T_u + k_6 C + \pi_3 - d_3 T_c;$$

**Variables:**

- *Tu*: Tumor cell population
- *Neo*: Extracellular neoantigen concentration
- *C*: Antigen-Antibody immune complexes
- *A*: Free antibody concentration
- *Tc*: Cytotoxic CD8+ T cell population

The model incorporates tumor growth ($k_1$), steady-state influx ($\pi_1$), and three clearance mechanisms: natural death ($d_1$), antibody-dependent cellular cytotoxicity (saturated term $\beta\frac{A}{A+k_m}$), and T cell-mediated lysis ($\alpha T_c T_u$). Neoantigens are released upon tumor cell death ($\rho_1$ term) and neutralized by antibodies to form complexes ($C$).

##### 13.2 Parameter Values and Equilibrium Analysis

Parameters and initial conditions are detailed in **Supplementary Table S18**. Parameter selection was performed via reverse deduction from biologically plausible steady-state values observed in clinical literature.

**Supplementary Table S18 | Parameters and initial values for Model 3.5.1**

| Variable | Initial Value ($t$=0) | Parameter | Value |
|---|---|---|---|
| Tu | 0 | $k_1$ | 1 |
| A | $10^5$ | $k_2$ | $10^{-7}$ |
| C | 0 | $k_3$ | 0 |
| Neo | 0 | $k_{4,5}$ | 1,2 |
| Tc | $10^3$ | $k_6$ | $2\times10^{-4}$ |
| | | $\pi_{1,2}$ | 100 |
| | | $\pi_3$ | 0.1 |
| | | $d_1$ | 0.05 |
| | | $d_{2,3}$ | 0.001 |
| | | $\rho_1$ | $10^4$ |
| | | $\rho_2$ | 0.1 |
| | | $\alpha$ | $10^{-4}$ |
| | | $\beta$ | 10 |
| | | $k_m$ | $10^7$ |

**Stability Analysis:**

The system possesses a non-trivial endemic equilibrium point $E^*$:

($Tc^*\approx2.01\times10^5$, $Tu^*\approx3.44$, $Neo^*\approx1.00\times10^4$, $C^*\approx1.03\times10^6$, $A^*\approx1.03\times10^9$)

The Jacobian matrix $J$ evaluated at $E^*$ is given by:

$$J = \begin{pmatrix} k_1 - d_1 - \beta\frac{A}{A+k_m} - \alpha T_c & 0 & 0 & \frac{-\beta k_m T_u}{(A+k_m)^2} & -\alpha T_u \\ \rho_1(d_1 + \beta\frac{A}{A+k_m} + \alpha T_c) & -k_2 A & k_3 & -k_2(Neo) + \frac{\rho_1 \beta k_m T_u}{(A+k_m)^2} & \rho_1 \alpha T_u \\ 0 & k_2 A & -k_3 - k_4 & k_2(Neo) & 0 \\ 0 & -k_2 A & k_3 + k_5 & -k_2(Neo) - d_2 & 0 \\ -\rho_2 \alpha T_c & 0 & 0 & k_6 & -\rho_2 \alpha T_u - d_3 \end{pmatrix}$$

The eigenvalues of this matrix are $\lambda_1$=−103.452, $\lambda_2$=−8.954, $\lambda_3$=−0.998, $\lambda_4$=−0.0026, and $\lambda_5$=−0.0020. Since Re($\lambda_i$) < 0 for all *i*, the equilibrium state is locally asymptotically stable.

##### 13.3 Simulation Results: Tumor Progression

Simulations were initialized from a disease-free state (assuming $T_u(0)$ =0 and sudden oncogenic transformation).

**Baseline Dynamics ($k_1$=1.0, Fig. S17):**
The system exhibits an initial proliferation phase of tumor cells followed by stabilization. Although the system reaches a mathematically stable equilibrium, the transient peak represents the biological manifestation of clinical cancer.

**Varying Tumor Growth Rates (Fig. S18):**
Increasing the tumor proliferation coefficient to $k_1$=1.1 results in a significantly higher peak tumor burden during the early phase. The system approximates a new stable equilibrium ($Tc*\approx 2.02\times 10^5$, $A*\approx 1.04\times 10^9$), with eigenvalues maintaining negative real parts, indicating that the system retains stability despite the higher pathological load.

**Impact of Immune Competence ($k_5$, Fig. S19 – S20):**
The parameter $k_5$ reflects the strength of antibody-mediated feedback (immune strength).

- Decreasing $k_5$ to 1.9 (Supplementary Fig. S19) leads to a marked increase in peak tumor burden.
- A further reduction to $k_5$=1.8 (Supplementary Fig. S20) results in dramatic tumor proliferation.
  While the mathematical equilibrium remains stable, the massive transient expansion of tumor cells corresponds to symptomatic, clinical progression of cancer driven by immune senescence or compromise.

##### 13.4 Therapeutic Scenarios

**Immune Checkpoint Blockade (Fig. S21):**
The parameter $\rho_2$ governs the exhaustion or inhibition of T cells upon interaction with tumor cells. Reducing $\rho_2$ simulates the effect of PD-1/PD-L1 inhibitors. As shown in **Supplementary Fig. S21**, lowering $\rho_2$ significantly suppresses the peak tumor burden.

**Monoclonal Antibody Therapy (Fig. S22):**
To model exogenous antibody therapy, we extended model 3.5.1 by introducing therapeutic antibodies $A'$:

$$\frac{dT_u}{dt} = \pi_1 + k_1T_u - d_1T_u - \beta\frac{A+A'}{A+A'+k_m}T_u - \alpha T_cT_u;$$
$$\frac{d(Neo)}{dt} = -k_2(Neo)A + k_3C + \rho_1(d_1T_u + \beta\frac{A+A'}{A+A'+k_m}T_u + \alpha T_cT_u);$$

$$\frac{dC}{dt} = k_2(Neo)A - k_3C - k_4C;$$

$$\frac{dA}{dt} = -k_2(Neo)A + k_3C + k_5C + \pi_2 - d_2A;$$

$$\frac{dT_c}{dt} = -\rho_2\alpha T_cT_u + k_6C + \pi_3 - d_3T_c;$$

$$\frac{dA'}{dt} = -k_2(Neo)A' + k_3C' - d_4A';$$

(See **Supplementary Table S19** for parameters).

**Supplementary Table S19 | Parameters and initial values for monoclonal antibody therapy**

| **Variable** | **Initial Value (*t*=0)** | **Parameter** | **Value** |
|---|---|---|---|
| Tu | 0 | $k_1$ | 1.1 |
| A | $10^5$ | $k_2$ | $10^{-7}$ |
| C | 0 | $k_3$ | 0 |
| Neo | 0 | $k_4$ | 1 |
| Tc | $10^3$ | $k_5$ | 2 |
| A′ | 0 | $k_6$ | $2*10^{-4}$ |
| C′ | 0 | $\pi_1$ | 100 |
| | | $\pi_2$ | 100 |
| | | $\pi_3$ | 0.1 |
| | | $d_1$ | 0.05 |
| | | $d_2$ | 0.001 |
| | | $d_3$ | 0.001 |
| | | $d_4$ | 0.05 |
| | | $\rho_1$ | $10^4$ |
| | | $\rho_2$ | 0.1 |
| | | $\alpha$ | $10^{-4}$ |
| | | $\beta$ | 10 |
| | | $k_m$ | $10^7$ |

Administration of antibodies at t=10 revealed a dose-dependent suppression of tumor cell proliferation (Supplementary Fig. S22), confirming the efficacy of targeting neoantigens via exogenous immunoglobulins.

**Neoantigen Vaccination (*Neo* Addition):**

The timing of neoantigen administration determines the therapeutic outcome:

- **Early Intervention (*t*=5, Fig. S23A):** Administration during the incipient phase effectively primes the immune system, controlling tumor growth.

- **Late Intervention ($t$=10, Fig. S23B):** Administration during the active growth phase exacerbates tumor proliferation. This counter-intuitive result arises because the influx of soluble neoantigens acts as a "sink," depleting available antibodies ($A$) through complex formation ($-k_2(Neo)A$), thereby reducing the antibody-dependent cytotoxicity directed at the tumor.

**MHC-I vs. MHC-II Modulated Vaccines:**
We differentiated vaccines based on their downstream effector mechanisms:

1. **MHC-I Agonists (Fig. S24):** Modeled by enhancing the T-cell proliferation feedback ($k_6$ ). Increasing $k_6$ effectively effectively curtails early tumor expansion by boosting the cytotoxic T cell response.
2. **MHC-II Agonists (Fig. S25):** Modeled by enhancing antibody proliferation signals ($k_5$). Increasing $k_5$ leads to superior control of the initial tumor outbreak by augmenting humoral immunity without the depletion risks associated with direct neoantigen flooding.

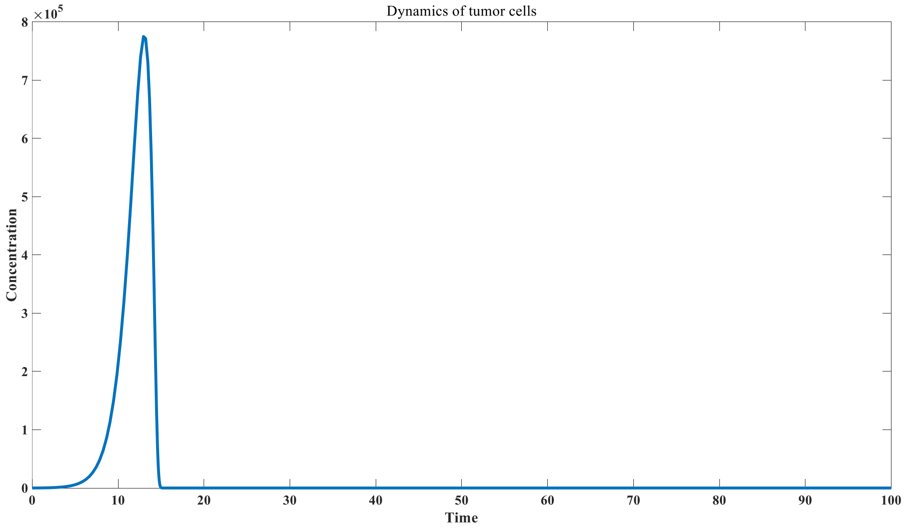


Figure S17: Dynamics of tumor cells when $k_1$= 1.0

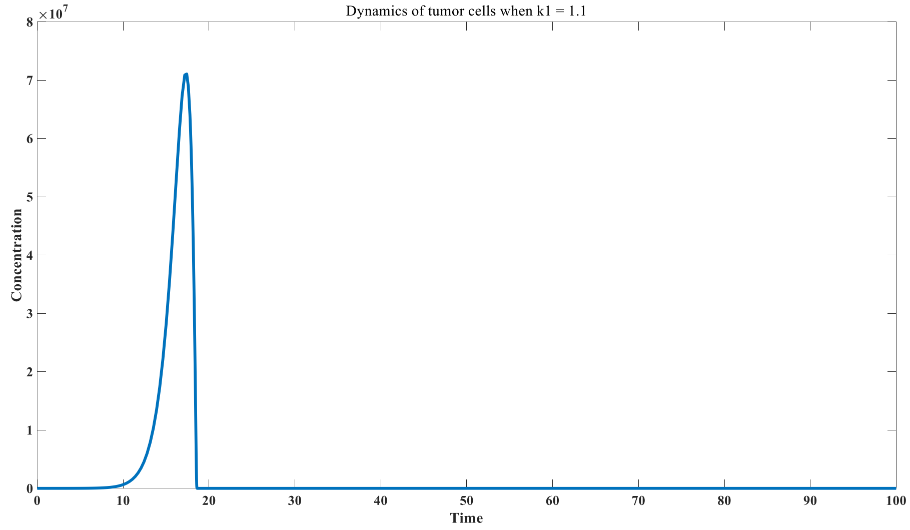


Figure S18: Dynamics of tumor cells when $k_1$= 1.1

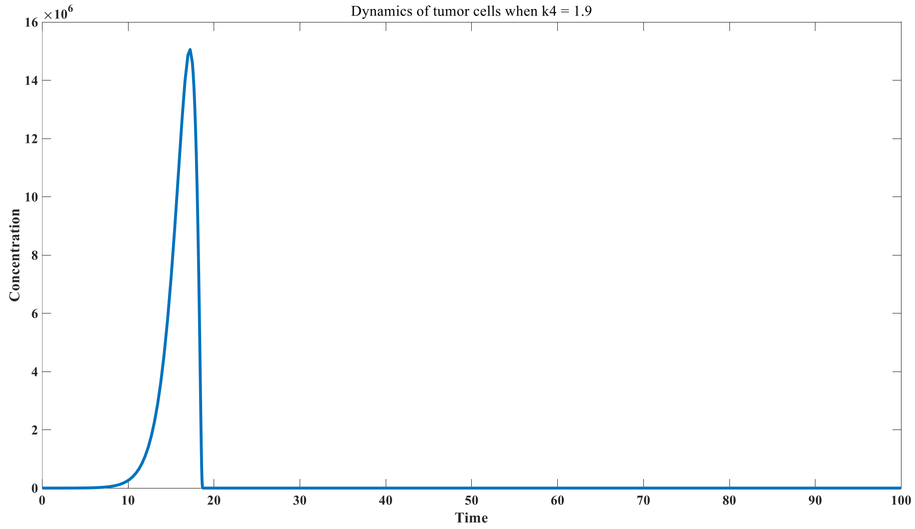


Figure S19: Dynamics of tumor cells when $k_5$= 1.9

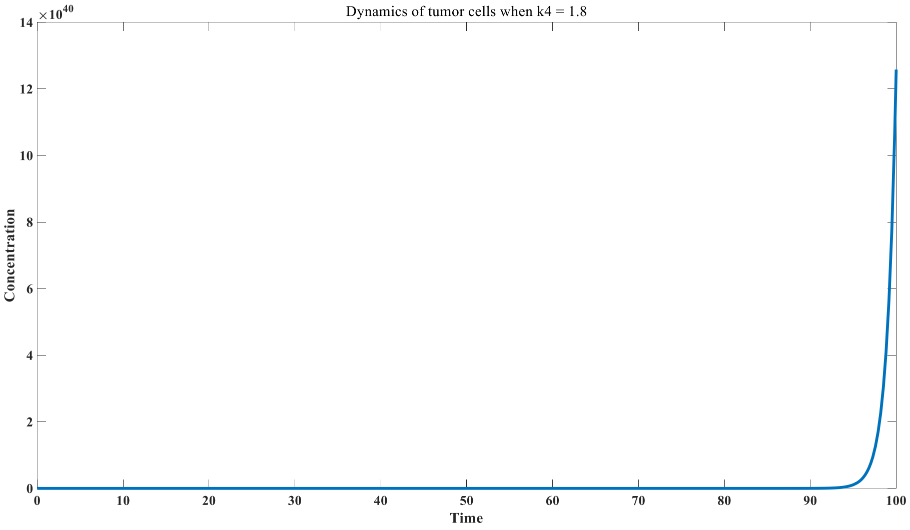


Figure S20: Dynamics of tumor cells when $k_5$= 1.8

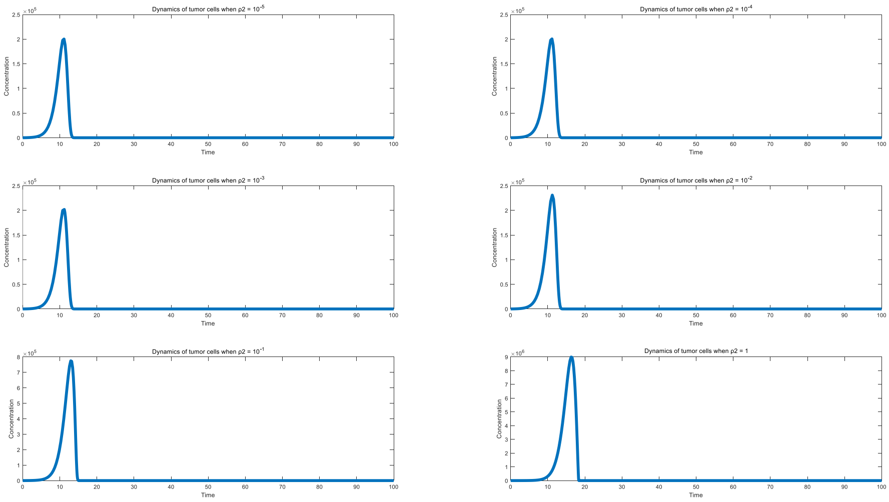


Figure S21: Mechanism of PD-1 inhibitor on tumor control

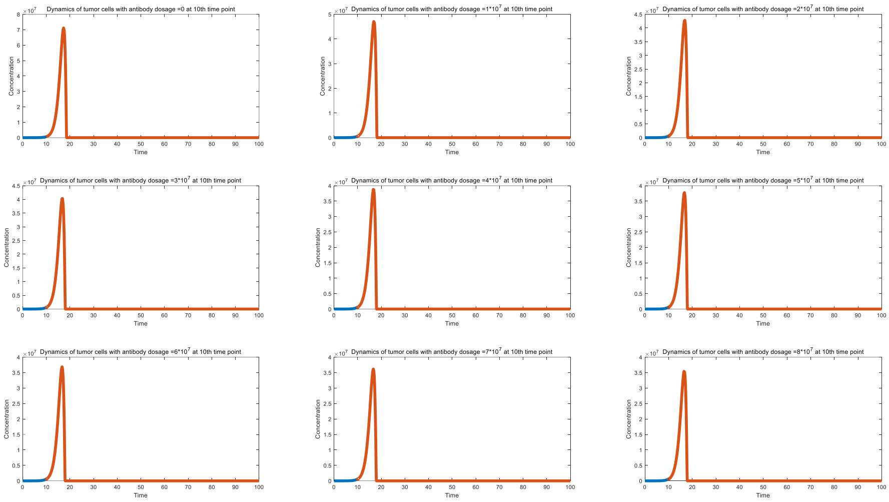


Figure S22: Effect of monoclonal antibody on tumor control

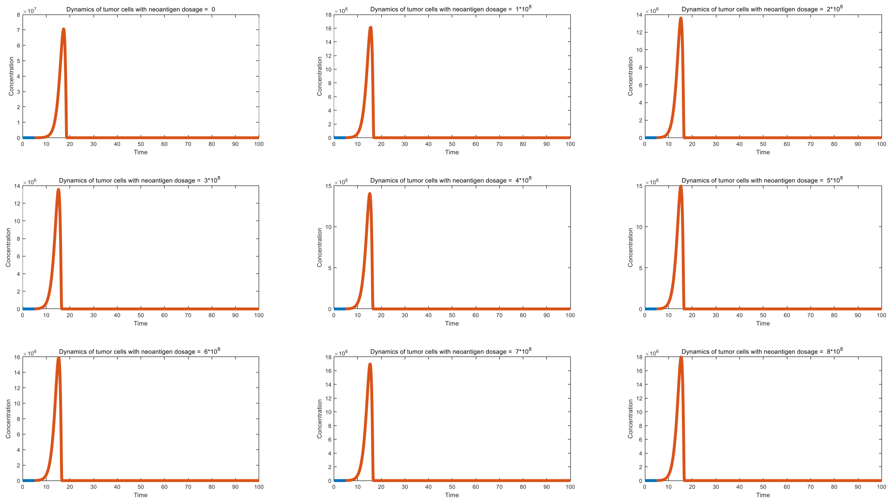


Figure S23_A: Effect of neoantigen addition on tumor control（vaccination at 5 th time point)

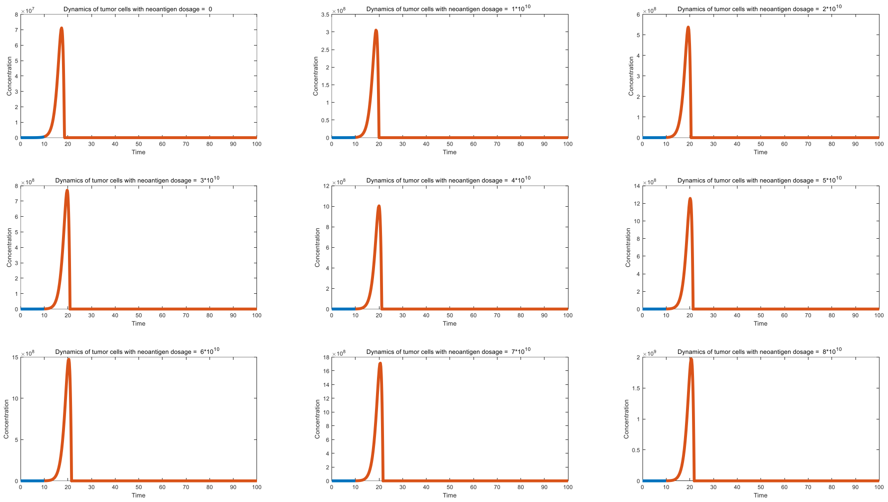


Figure S23_B: Effect of neoantigen addition on tumor control（vaccination at 10 th time point)

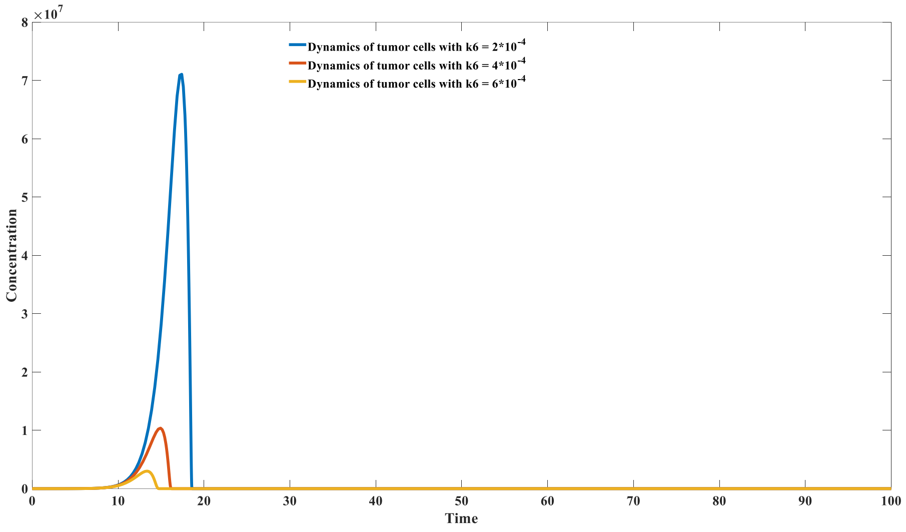


Figure S24: Mechanism of MHC-1 neoantigen vaccine on tumor control

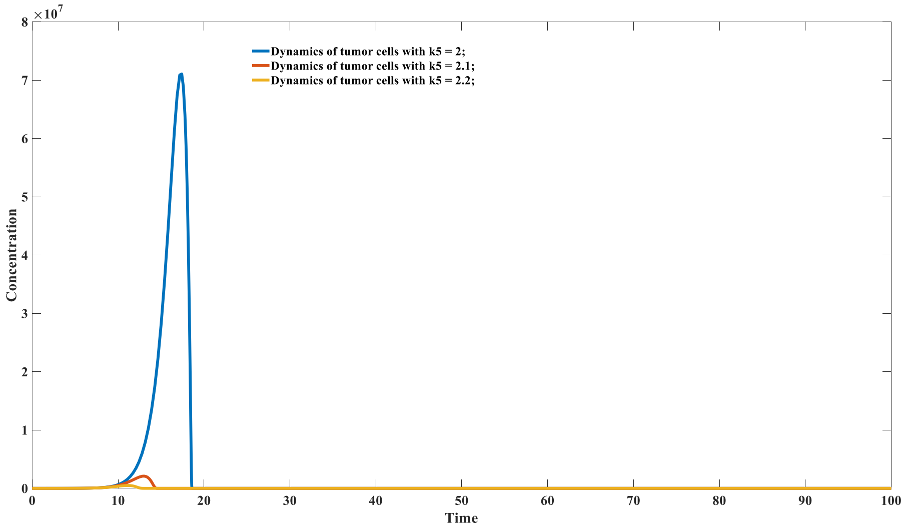


Figure S25: Mechanism of MHC-II neoantigen vaccine on tumor control
